# Multi-nodal regulation of olfactory dishabituation in Drosophila

**DOI:** 10.1101/2025.01.07.631716

**Authors:** A G. Charonitakis, S Pasadaki, E-M Georganta, K Foka, O Semelidou, EMC Skoulakis

## Abstract

Habituation adaptively filters repeated, inconsequential sensory input, while the response to such stimuli re-emerges upon appearance of novel or salient cues (dishabituation).However, the neural circuits underlying dishabituation remain poorly defined, particularly in the central nervous system. Using Drosophila olfactory habituation to the odorant 3-octanol (OCT), we dissect the circuit basis of intramodal (odor-odor) and cross-modal (footshock-odor) dishabituation. A brief yeast odor puff dishabituates intramodally, whereas footshock cross-modally and neither operate through sensitization. Genetic silencing and optogenetics demonstrate that Mushroom Body (MBs) output drives both dishabituation forms. αβ and γ Kenyon cells (KCs) mediate dishabituation, while α′β′ Kenyon cells mediate habituation. Dopaminergic neurons encode and PAM neurons mediating appetitive and PPL1 neurons aversive dishabituation, including that triggered by footshock and OCT itself. GABAergic APL neurons and specific MB output neurons tune the balance between habituation and dishabituation, relaying signals via MBONs to the Lateral Horn, a proposed decision node. Connectomic analysis reveals inhibitory interactions supporting this balance. Collectively, we reveal a multi-node circuit that dynamically overrides, rather than erases, habituation findings offering insight into habituation deficits in intellectual disability, autism, and schizophrenia.

**SIGNIFICANCE STATEMENT:** This study identifies a multi-node neural circuit spanning the Mushroom Bodies, dopaminergic and GABAergic modulatory neurons and the Lateral Horn, that governs intramodal and cross-modal dishabituation in Drosophila. Showing that dishabituation dynamically overrides rather than erases habituation and that distinct dopaminergic populations encode stimulus valence to bias this switch, the work reveals a general logic for how the brain flexibly regulates sensory filtering.

## INTRODUCTION

Animals must continuously filter sensory input, selectively attending to stimuli relevant for survival and reproduction, while suppressing responses to repetitive, inconsequential ones. Habituation is the behavioral manifestation of salience-filtering processes mediating reduced responses to repeatedly encountered percepts lacking biological significance driven by attenuated output from the Stimulus-Response (S-R) circuits mediating attraction or avoidance (Groves and Thompson 1979, Rankin, Abrams et al. 2009, Ramaswami 2014). Upon removal of the habituated percept, the attenuated response recovers spontaneously toward its naïve baseline over a relatively short time (Thompson and Spencer 1966, Rankin, Abrams et al. 2009, Ramaswami 2014, Cooke and Ramaswami 2020). Responsiveness to the habituated stimulus can also re-emerge through exposure to a novel stimulus (Thompson and Spencer 1966, Rankin, Abrams et al. 2009), or a shift in the behavioral relevance of the habituated percept sufficient to recruit attentional processing (Kato, Gillet et al. 2015). Processes underlying this response recovery are collectively termed dishabituation. Critically, dishabituation distinguishes true habituation from sensory fatigue or peripheral adaptation and is a defining hallmark of behavioral flexibility (Rankin, Abrams et al. 2009, Ramaswami 2014). Dishabituation is essential to detect and respond to sudden, potentially threatening stimuli embedded within otherwise familiar or invariant contexts. This is consistent with observations that moderately intense, behaviorally salient stimuli serve as the most effective dishabituators (Thompson and Spencer 1966, Rankin, Abrams et al. 2009).

How do dishabituating stimuli reinstate the habituated response? The Dual Process Theory (Groves and Thompson 1979, Thompson 2009), developed primarily from studies of fast sensory-to-motor reflex circuits (Glanzman 2008, Thompson 2009), proposes two competing processes. Synaptic depression at specific S-R circuit synapses drives response decrement, while a suddenly appearing novel stimulus sensitizes broadly the entire interrogated circuit, or the nervous system at large, reinstating the decremented response. The observation that the reinstated response quickly returns to habituated levels was taken as evidence that the two processes operate independently and antagonistically (Groves and Thompson 1979). However, this working framework was challenged by work in *Aplysia* demonstrating that sensitization and dishabituation engage distinct molecular mechanisms (Hochner, Klein et al. 1986, Antonov, Kandel et al. 1999) and furthermore, differ in onset kinetics, stimulus intensity requirements, and developmental trajectories (Marcus, Nolen et al. 1988). Critically, rather than sensitization masking an intact habituated state, dishabituating stimuli were thought to actively disrupt habituation within the S-R circuit itself (Steiner and Barry 2014). Work across multiple experimental systems agreed with this conclusion (Ehrlich, Boulis et al. 1992, Steiner and Barry 2011, Steiner and Barry 2014). Evidence from *Drosophila* further dissociates the two processes genetically. Flies jump to escape a sudden aversive odor puff and this response habituates upon repeated stimulation and can be dishabituated by vortexing (Asztalos, Arora et al. 2007). Loss-of-function mutations in the cytoplasmic tyrosine kinase Btk selectively impair habituation in this assay, leave dishabituation intact, but abolish sensitization (Asztalos, Arora et al. 2007). Genetic dissociation of dishabituation from sensitization strongly supports their mechanistic independence.

A critical mechanistic difficulty with the synaptic depression model of habituation is that it cannot readily explain how a desensitized synapse recovers rapidly upon dishabituation (Cooke and Ramaswami 2020), nor how it scales to the complex, multi-layered circuitry of the CNS. In fact, the circuitry engaged by dishabituators in the Central Nervous System (CNS) and how it interfaces with circuits mediating habituation has remained poorly explored. However, the Negative Image model (Ramaswami 2014) offers a more tractable framework for the complex fully centralized CNS. It proposes that repeated exposure to inconsequential stimuli progressively potentiates an inhibitory trace that heterosynaptically selectively suppresses responses to those stimuli, producing habituation (Ramaswami 2014, Cooke and Ramaswami 2020). Neurotransmission, in response to the novel dishabituator does not reverse synaptic depression, but rather, independently suppresses the familiarity-encoding inhibitory trace, thus releasing the habituated circuit from inhibition via disinhibition (Ramaswami 2014), yielding reemergence of the attenuated response. This mechanism is directly instantiated in rat piriform cortex, where olfactory habituation is driven by type III metabotropic glutamate receptor-mediated synaptic depression, and dishabituation is triggered by loud auditory stimuli acting through β-adrenergic receptor activation to suppress the inhibitory trace (Smith, Shionoya et al. 2009). A parallel logic operates in a *Drosophila* paradigm. Repeated sucrose stimulation of GRN64-expressing gustatory receptor neurons habituates the proboscis extension reflex by potentiating GABAergic inhibitory neurons. Novel gustatory stimuli engaging GRNs other those responsive to sucrose, recruit dopaminergic neurons (DANs) that, directly or indirectly, suppress GABAergic activity, thus overriding the response attenuation and reinstating the habituated response, manifested as dishabituation (Trisal, Aranha et al. 2022).

Our previous work indicates that effective dishabituation depends on the relationship between the habituating stimulus and the dishabituating cue. We have established habituation paradigms to recurrent footshocks (Acevedo, Froudarakis et al. 2007, Roussou, Papanikolopoulou et al. 2019, Foka, Georganta et al. 2022) and to 4-minute continuous exposure to the odorant 3-Octanol (OCT) {Semelidou, 2019 #3; Fig 1A} in Drosophila. These paradigms present with modality-dependence of effective dishabituation in that footshock habituation is rescued by a brief OCT or yeast odor (YO) puff, but not by visual stimuli or vortexing, a mechanosensory stimulus likely akin to footshock (Acevedo, Froudarakis et al. 2007, Roussou, Papanikolopoulou et al. 2019). Reciprocally, a single footshock or brief vortexing fully reinstates the attenuated response following habituation to 4-min OCT habituation, while YO exposure can also dishabituate from 30-min OCT exposure {Semelidou, 2018 #4; Fig 1B}. Spontaneous recovery of naïve response occurs within 4-6 minutes after removal of the OCT {Semelidou, 2018 #4; Fig 1B}, or footshok (Acevedo, Froudarakis et al. 2007) habituators. Collectivelly these findings suggested that changes in sensory context through exposure to a different stimulus modality provide a critical signal for dishabituation and raised the possibility that dishabituation engages neural circuits distinct from those driving the habituated response. This cross-modality in itself may constitute the novelty signal considered essential for dishabituation (Groves and Thompson 1979, Rankin, Abrams et al. 2009) and implies that dishabituation recruits neuronal circuits non-overlapping with those driving the habituated response.

**Figure 1:**
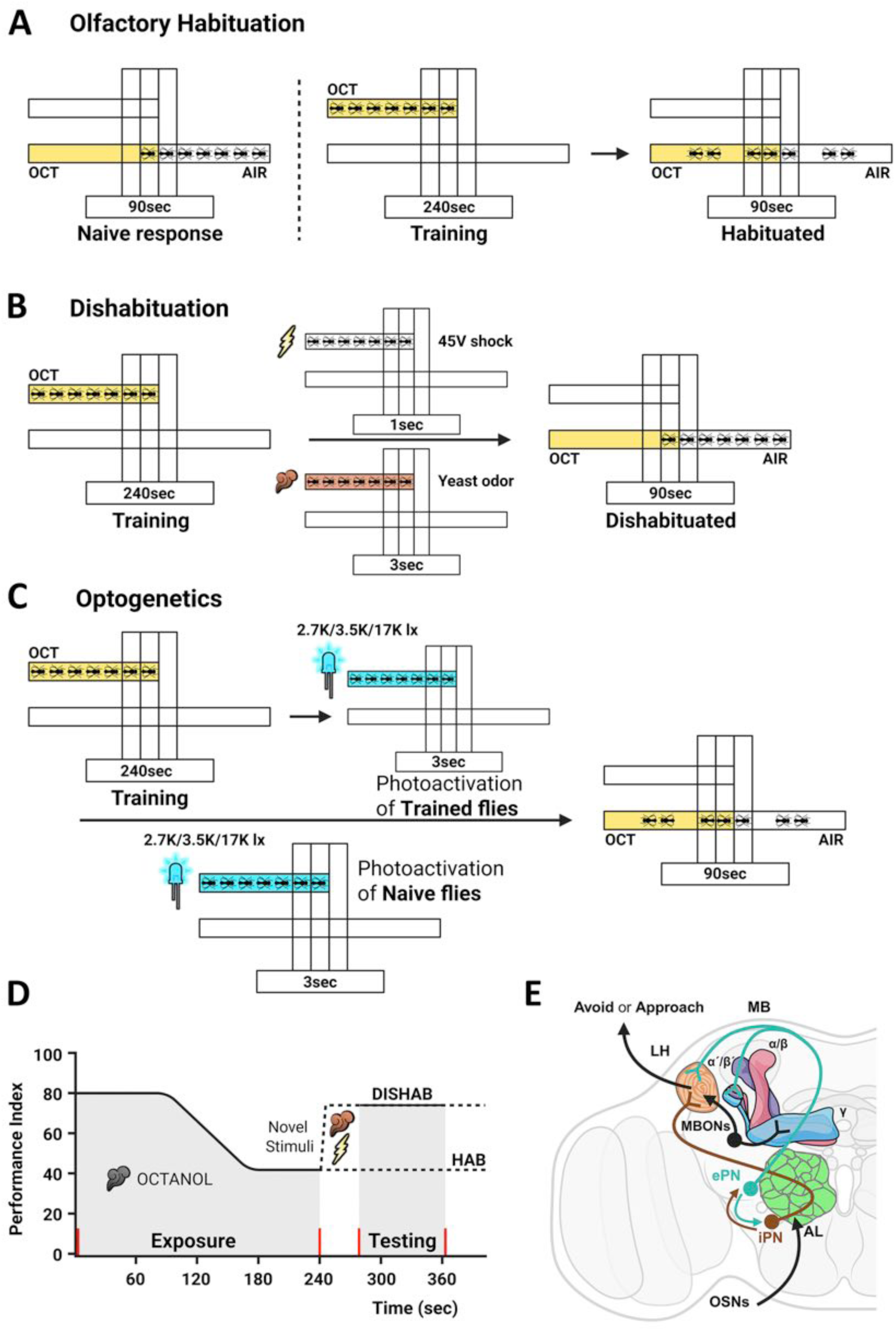
Olfactory habituation, dishabituation, and the optogenetic stimulation paradigm. Footshocks are represented by the lightning bolt and odor by the clouds of odor-specific color. **A)** Naïve flies exhibit an aversive osmotactic response to Octanol (OCT). After 4 min of continuous OCT exposure, the flies show reduced avoidance to OCT, indicative of habituation. **B)** Presentation of a novel stimulus (electric shock or yeast odor), following 4 min of OCT exposure results in reinstatement of the naïve response to OCT, demonstrating dishabituation. **C)** Optogenetic activation was delivered with a continuous 3-sec light pulse at 460nm, of three different light intensities. Stimulation was applied either after OCT exposure to assess its effect on dishabituation or in naïve flies to detect the effect on baseline avoidance. **D)** Schematic representation of the olfactory habituation – dishabituation behavioral paradigm showing the time lines of exposure and testing and expected OCT avoidance levels **E)** The model of short exposure olfactory habituation (Semelidou, Acevedo et al. 2018), establishing that habituation requires inhibition of the LH-driven innate response to OCT. Model created with Biorender.com https://BioRender.com/0qwwz1v.

However, this interpretation is complicated by the finding that olfactory and visual stimuli fail to dishabituate gustatory habituation in *Drosophila*, while in fact a gustatory stimulus engaging distinct sensory neurons is an effective dishabituator (Trisal, Aranha et al. 2022). This argues that effective dishabituators must share neuronal elements, or converge on circuits engaged by the habituating stimulus, rather than operating through entirely orthogonal pathways. Resolving this contradiction demands direct interrogation of the circuits recruited by dishabituating stimuli and how they interface with habituation circuits in the CNS. Such questions have generally remained poorly addressed despite recent progress (Smith, Shionoya et al. 2009, Trisal, Aranha et al. 2022), and the answers are essential towards mechanistic insights to the neural basis of adaptive sensory filtering, dishabituation mechanisms and requirements.

Here, we address these questions by dissecting the circuit mechanisms underlying both intramodal and cross-modal dishabituation in Drosophila olfactory habituation. Drosophila is uniquely positioned to address these questions, enabling circuit-level dissection of dishabituation with precision inaccessible in most other systems through its powerful genetic toolkit, combined with a largely elucidated and functionally validated brain connectome (Winding, Pedigo et al. 2023, Schlegel, Yin et al. 2024). We utilized genetic and optogenetic tools to selectively manipulate neurotransmission in adult *Drosophila* brain circuits and determine their contributions to intramodal and cross-modal dishabituation of 4-minute OCT habituation. As habituation and dishabituation are conserved (Cooke and Ramaswami 2020), mechanistic insights from Drosophila will likely be directly relevant to human pathology. Habituation deficits are a broadly shared feature of intellectual disabilities, autism, and schizophrenia (Ramaswami 2014, McDiarmid, Bernardos et al. 2017, Coll-Tane, Krebbers et al. 2019, Fenckova, Blok et al. 2019, McDiarmid, Belmadani et al. 2020, Foka, Georganta et al. 2022), highlighting the impact of identifying the neural circuits that regulate and override habituation.

## RESULTS

### The habituation-dishabituation paradigms

To investigate the circuitry and mechanism(s) of intramodal and cross-modal dishabituation, we employed the olfactory habituation to OCT paradigm (Fig 1A and (Semelidou, Acevedo et al. 2018)). The paradigm consists of modifying the naïve aversion to OCT by exposing 40-60 mixed sex Drosophila to this odor for 4 minutes, which attenuates significantly the naïve avoidance of this odor (Fig 1A) (Semelidou, Acevedo et al. 2018) and conforms to habituation characteristics (Thompson and Spencer 1966, Rankin, Abrams et al. 2009).

The S-R route engaged by the 4-min OCT exposure involves Olfactory Receptor Neurons (OSNs), which via excitatory Projection Neurons (ePNs) reach the Lateral Horn (LH) and the Mushroom Bodies (MBs). In addition, inhibitory Projection Neurons (iPNs) relay the information exclusively to the LH (Fig 1E). The MBs comprise bilateral clusters of about 2000 neurons, the Kenyon Cells (KCs) in the dorsal posterior of the adult Drosophila brain with distinct tripartite axonal morphology (Crittenden, Skoulakis et al. 1998) and are essential for olfactory associative learning and memory (Modi, Shuai et al. 2020, Davis 2023). The MBs receive major inputs from the olfactory system represented by sparse activation of particular intrinsic neurons (sparse coding, (Honegger, Campbell et al. 2011)). They also receive inputs from other sensory modalities and gate responses mediated by the LH (Modi, Shuai et al. 2020, Vrontou, Groschner et al. 2021). The LH is also a bilateral paired neuropil in the protocerebrum, which mediates innate behavioral responses (Schultzhaus, Saleem et al. 2017, Das Chakraborty, Chang et al. 2022), and is comprised of multiple types of LH interneurons (LHNs), thought to mediate odor discrimination and valence among other functions. LH output neurons (LHONs) mediate behavioral responses by impinging on premotor and motor neurons (Dolan, Frechter et al. 2019, Mohamed, Hansson et al. 2019). Importantly, a good number of LHONs are proximal to DAN dendrites and converge with Mushroom Body Output neurons (MBONs) indicating that they can modulate DANs and integrate MB-mediated informational outputs (Dolan, Frechter et al. 2019).

Avoidance in naïve animals is mediated by LHNs encoding valence information (Dolan, Frechter et al. 2019, Mohamed, Hansson et al. 2019). To maintain the naïve avoidance response to OCT during the initial minutes of exposure (habituation latency) requires GABAergic inhibitory activity of local interneurons in the antennal lobe and excitatory projection neurons (ePNs) impinging on the LH and the MBs ((Semelidou, Acevedo et al. 2018) and Fig 1E), suggesting that excitatory inputs to the LH drive OCT avoidance. In contrast, habituation to OCT requires neurotransmission from both excitatory and inhibitory projection neurons converging on the LH. The increased negative impact of the 4-min OCT exposure and increased activation of the LH-only-projecting iPNs drives feedforward inhibition (Wang, Gong et al. 2014), suppressing the innate avoidance of OCT, likely by inhibiting LHON signals to relevant motor neurons (Semelidou, Acevedo et al. 2018).

Dishabituation in this paradigm is achieved by a single 1-sec 45V footshock after a 4-min OCT exposure to the odorant and is manifested as re-emergence of OCT avoidance (Fig 1B, D). The temporal sequence of events in the assay and expected avoidance levels post the 4-min exposure to OCT and after experiencing the dishabituators are summarized schematically in Figure 1D. In accord with Thomson and Spencer tenets (Thompson and Spencer 1966, Rankin, Abrams et al. 2009), and prior results (Semelidou, Acevedo et al. 2018), undiluted (100%) OCT did not dishabituate flies habituated to 0.1-100% of the same odorant, whereas a single footshock was an efficient dishabituator (Suppl Fig 1A,B). Therefore, strong odor stimulation is not an intramodal dishabituator for flies habituated to lower concentrations of the same odorant.

### Yeast puff is an efficient intramodal dishabituator

Prior results indicated that a 3-sec YO stimulus dishabituated flies from 30-min exposure to OCT (Das, Sadanandappa et al. 2011, Semelidou, Acevedo et al. 2018). Hence, initially we asked whether YO can dishabituate flies after the reduced in timescale OCT exposure of 4 min (Semelidou, Acevedo et al. 2018). Indeed, after 4-min exposure to OCT, a 3sec YO exposure resulted in re-emergence of OCT avoidance, as effectively as for the footshock cross-modal dishabituator in two different control strains (Fig 2A, B). Therefore, YO is an efficient dishabituator of habituation to 4-min OCT, as is for 30-minute exposure to the same odor (Semelidou, Acevedo et al. 2018). Because the results with Canton S and Cantonized-*w^1118^* animals were identical, only the latter strain was used as a wild type control in all subsequent experiments.

**Figure 2:**
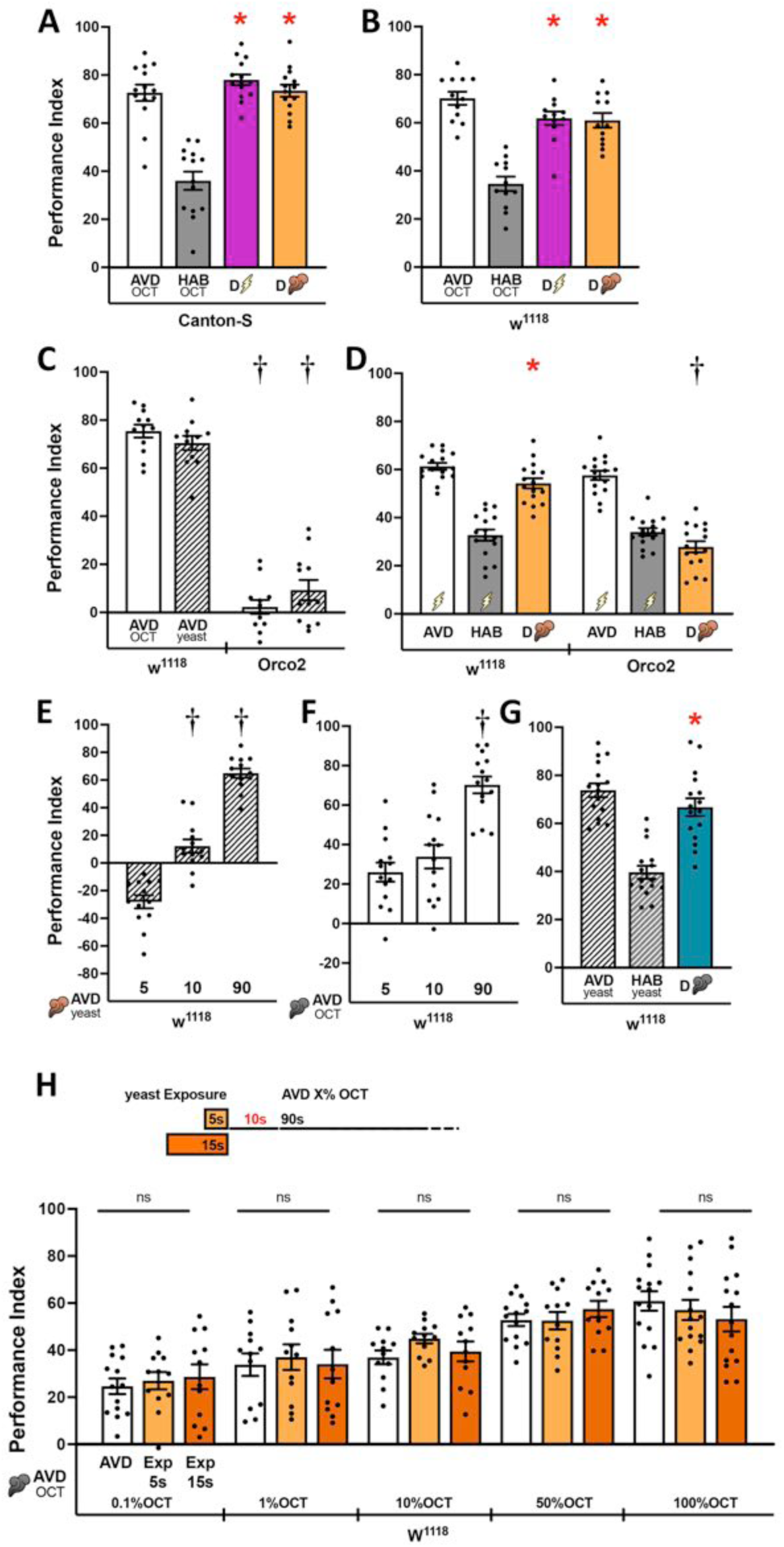
Intramodal and cross-modal dishabituation from OCT habituation. **A-H)** Mean Performance Indices ± SEM are shown in all panels. The naïve avoidance response (AVD) is represented by the white bar for OCT and white striped bar for yeast odor (YO). The habituated response (HAB) is shown by the grey bars for OCT and the striped, grey bar for YO. The dishabituated response is shown in magenta for the footshock dishabituator, in orange for YO and in cyan for OCT. Pre-exposures of 5sec and 15sec to YO are shown in orange and dark orange respectively. Stars indicate significant differences (p<0.005) from the habituated response and daggers significant difference (p<0.005) from naïve avoidance. All statistical details are presented in Supplemental Table 1. **A, B)** Naïve responses of Canton-S and *w^1118^* flies to OCT (AVD) and habituated (HAB) responses to OCT, dishabituated either by a single 45V electric shock (bolt) or a puff of yeast odor (plume). n=14 for B and n=12 for C. **C)** Naïve response of *w^1118^* and *Orco^2^* homozygotes to OCT and YO. Daggers indicate significant differences between the respective naïve responses to OCT and YO of Orco^2^ homozygotes controls. n=12 **D)** Footshock habituation cannot be dishabituated by YO presentation in *Orco^2^* homozygotes, unlike in control animals. n=16. **E)** Time-dependent changes in the response of *w^1118^* flies to YO. Daggers indicate significant differences (p<0.005) between the responses after 10 and 90 sec compared to the shortest exposure (5 sec) to YO. n=13 **F)** Time-dependent changes in the response of *w^1118^* flies to OCT. Daggers indicate significant differences (p<0.005) between the response after 90 sec compared to the shorter exposures (5 and 10 sec) to OCT. n=14 **G)** Habituation of control flies to YO and dishabituation by OCT presentation. n=16. **H)** Pre-exposure of *w^1118^* flies to YO for the indicated durations, followed by a stimulus-free interval of 10 sec, does not affect subsequent responses to a range of OCT dilutions. n≥12

Is in fact the YO puff an intramodal dishabituator? In flies, yeast as a tastetant was shown to dishabituate the proboscis extension reflex to sucrose (Trisal, VijayRaghavan et al. 2023). Although the flies do not come in contact with the yeast solution, we wondered whether YO is perceived as an odor in our assay, hence an intramodal dishabituator, or as a gustatory cross-modal dishabituator. If YO is perceived as an olfactory stimulus, then it should be perceived by control animals, but not anosmic *Orco* mutant flies (Larsson, Domingos et al. 2004). In fact, a 90-sec exposure to YO was as aversive to control *w^1118^* flies, as OCT was (Fig 2C, Suppl Fig 2A). In contrast, *Orco* mutant animals remained unresponsive to both odors (Fig 2C and Suppl Fig 2A). However, similar to *w^1118^* flies, *Orco* mutants habituated readily to recurrent footshocks (Acevedo, Froudarakis et al. 2007), but failed to dishabituate with a YO puff in contrast to control animals (Fig 2D and Suppl. Fig 2B). Therefore, like OCT, YO is a *bona fide* olfactory stimulus and thus, it acts as an efficient intramodal dishabituator of habituation to 4 min OCT.

Given that YO is a bona fide olfactory stimulus, it should also engage the canonical olfactory information processing pathway described above. However, YO is comprised of a bouquet of odorants that include sulfur-containing (ie Dimethyl sulfide), buttery (2,3 butanedione), nutty (furfural compounds), acids (isovaleric acid) and alcohols (Scheidler, Liu et al. 2015). Although we do not know the precise constitution of YO volatiles of the preparation we use, it nevertheless is unlikely that any of these odorants engage the same olfactory receptors as OCT. Therefore, the intramodal dishabituator is differentiated from the habituated stimulus initially at the sensory level likely by engaging distinct OSNs, which most likely result in distinct sparse coding signatures (Honegger, Campbell et al. 2011) than those signifying OCT at the level of the MBs.

Provided the nutritional significance of yeast for Drosophila, is YO always an aversive stimulus as indicated above (Fig 2C)? To determine its valence, the innate responses of control flies to YO were assessed at different lengths of exposure. Surprisingly, under our experimental conditions, whereas a 5-sec YO pulse was mildly attractive, a 10-sec pulse appeared neutral, while at 90 sec the odor was clearly aversive (Fig 2E). In contrast, both a 5 or a 10-sec puff of OCT were aversive, becoming highly aversive upon a 90-sec exposure (Fig 2F). Therefore, unlike the consistently aversive OCT, YO has a rewarding/appetitive valence at the 5-sec exposure typically used as a dishabituator in our experiments ((Semelidou, Acevedo et al. 2018, Foka, Georganta et al. 2022), Fig 2A, B).

The appetitive valence of brief YO exposure raised the question of whether effective dishabituation requires a change in stimulus valence relative to the habituated percept. However, following habituation to the aversive 90-sec YO exposure, flies readily dishabituated by the also aversive 5-sec OCT puff (Fig 2G). Importantly then, intramodal dishabituation does not require the habituated percept and the dishabituator to be of opposing valence. Two aversive odorants, engaging a similar overall sensory route to higher brain areas (Fig 1D and (Semelidou, Acevedo et al. 2018)), can be an effective habituator/dihabituator pair. These results suggest that the novelty of dishabituator rather than its valence relative to that of the habituated stimulus impacts its efficacy.

Is the re-emergence of OCT avoidance the result of sensitization by YO in accord to the predictions of Dual Process Theory (Carew, Castellucci et al. 1971, Groves and Thompson 1979)? To address this possibility, control animals were exposed to YO for the attraction-yielding 5-sec or the mildly aversive 15-sec exposure. After the 10-sec interval required to transfer the flies to the choice point of the maze (Fig 1A, B), avoidance of different dilutions of OCT was assessed for 90-sec. If YO exposure led to sensitization, post-exposure OCT avoidance should be enhanced compared to that of naïve animals, especially for the higher dilutions of the odorant. However, OCT avoidance over the range of dilutions tested remained unchanged after either 5 or 15 sec YO exposure (Fig 2H). Similar results were obtained when the interval between YO exposure and testing subsequent OCT avoidance was increased to 30- sec (Suppl Fig 2C). Therefore, sensitization does not underlie YO-mediated dishabituation from 4-min OCT exposure. In addition, previous results (Acevedo, Froudarakis et al. 2007), are not consistent with the possibility that footshocks, or exposure to another odor sensitize or suppress subsequent OCT avoidance. Collectively then, flies dishabituate their attenuated response to OCT by experiencing novel non-sensitizing stimuli that can be either cross-modal, or importantly, also intramodal olfactory stimuli independent of their valence.

### Neurotransmission from the Mushroom Body mediates cross-modal and intramodal dishabituation

Each odor generates a unique, sparse activity pattern in the KCs resulting in an activity code for odor identity (Turner, Bazhenov et al. 2008, Lin, Bygrave et al. 2014). Therefore, OCT and YO will be represented by such unique patterns within KCs (Campbell, Honegger et al. 2013), which are then expected to play a role, at least in intramodal dishabituation. Hence, we asked whether the MBs are required for intramodal and cross-modal dishabituation.

Prior work demonstrated that silencing all KCs (Fig 3A), impairs dishabituation with a single footshock (Semelidou, Acevedo et al. 2018). This conclusion was independently validated by conditionally silencing all KCs with expression of the temperature sensitive dynamin Shibire^ts^ (Shi^ts^) (Kitamoto 2001) under the pan-MB driver dncGal4. At the permissive 25°C (UN), Shi^ts^ expression did not impair habituation or dishabituation either by footshock or YO (Fig 3B). In contrast, attenuation of KC neurotransmission at the restrictive temperature (IN), suppressed both intramodal and cross-modal dishabituation manifested as suppressed OCT avoidance (Fig 3B). Moreover, since OCT and YO have distinct chemical composition and most likely engage distinct OSNs, it is highly unlikely that flies generalize the habituator with the intramodal dishabituator by overlapping KC activity patterns (Campbell, Honegger et al. 2013), thus failing to discriminate between them resulting in dishabituation failure (Fig 3B). Control heterozygotes exhibited normal YO and fooshock-mediated dishabituation at the restrictive temperature indicating that the induction regime did not interfere with the normal behavioral output (Fig 3E). Therefore, neurotransmission from KCs is necessary for both intramodal and cross-modal dishabituation.

**Figure 3:**
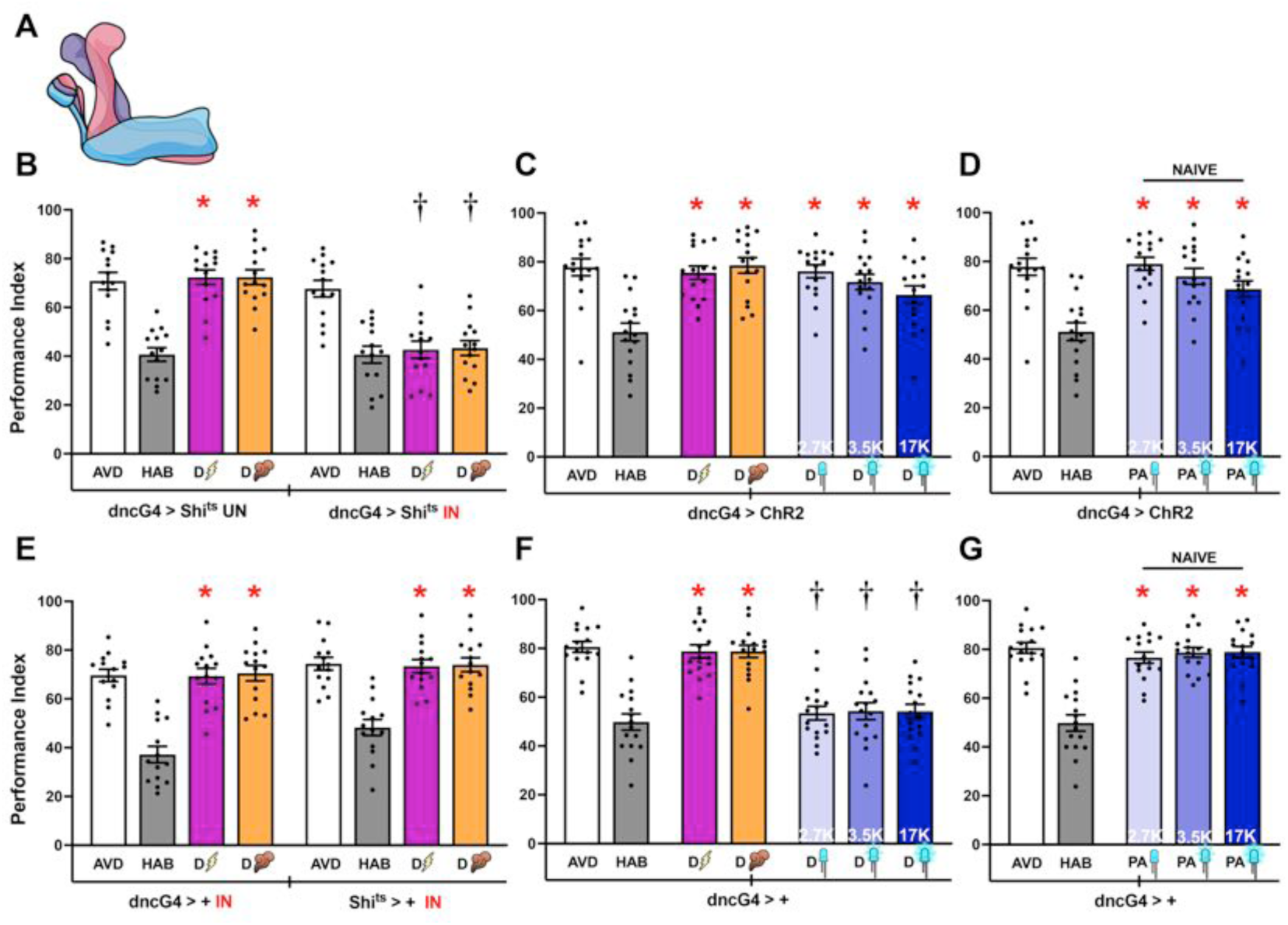
The mushroom bodies are essential for intramodal and cross-modal dishabituation. **A)** Schematic of the MB, illustrating its tripartite structure. **B-F)** Mean Performance Indices ± SEM are shown in all figures. The naïve response for OCT is shown in white bars and is always significantly different from the habituated response (grey bars). Cross-modal (footshock) dishabituation is indicated by the bolt (magenta bar) and intramodal (YO), by the plume (orange bar). Exposure to blue light is indicated by the LED bulbs and increasing intensity (as indicated by the respective Klx) by deepening shades of blue. Stars in all figures indicate significant differences (p<0.005), from the habituated response and daggers significant difference (p<0.005) from naïve avoidance. All statistical details are presented in Supplemental Table 1. **B)** Functional silencing of the MB neurons (IN) with UAS-shibire^ts^ under dncGal4 resulted in the expected habituation after 4 min of OCT exposure, but no dishabituation after a 45V electric shock (magenta bar) or YO (orange bar). At the permissive conditions (UN), habituation and both types of dishabituation were not affected. n=14 **C)** Flies expressing ChR2XXL in their MBs avoid, habituate and dishabituate with footshock or YO normally. Photoactivation of MB neurons in OCT-habituated flies with three different light intensities of 2.7K, 3.5K and 17K, restored OCT avoidance at all intensities. n=16 **D)** Photoactivation of MB neurons in naïve flies did not affect OCT avoidance significantly. n=16. **E)** Control dncGal4>+ and UAS-shibire^ts^>+ flies were subjected to the same induction conditions as the experimental group and did not present habituation or dishabituation impairments. n=14. **F)** Control dncGal4>+, not expressing ChR2XXL did not dishabituate upon photoactivation eliminating the possibility of dishabituation via visual stimulation. n=16. **G)** Blue light exposure at all intensities used did not affect OCT avoidance in naïve flies. n=16.

To determine whether KCs are also sufficient to drive dishabituation, we elicited neurotransmission from these neurons immediately after the 4-min habituating exposure to OCT by photostimulating flies expressing therein the light-sensitive Channel Rhodopsin (Fig 1C). To ascertain that photoactivation intensity did not yield artifacts due to subthreshold or excess activation, we assessed aversion to OCT after exposure to three different light intensities of 2.7 Klx, 3.5 Klx and 17 Klx. Photoactivation of the MBs with all three intensities resulted in recovery of OCT avoidance, behaviorally indistinguishable from that elicited either by the footshock or YO dishabituators (Fig 3C). Because photoactivation did not alter OCT avoidance of ChR2-expressing naïve flies (Fig 3D), we posit that it effectively emulates dishabituation rather than generally enhancing avoidance behavior (Fig 3C). However, as photoactivation of MBs, or any other neuronal type assayed below induces neurotransmission, it does not inform on the inputs that activate these neurons.

The blue light used to activate the ChR2 protein is visible to the flies (Morante and Desplan 2008). Therefore, we addressed the possibility that the observed dishabituation was not consequent of MB activation, but rather due to the light stimulus itself. To control for this possibility, flies heterozygous for the driver, hence not expressing ChR2, were tested for light-induced reversal of the attenuated OCT avoidance. While these flies habituated to OCT and dishabituated both by footshock and YO, blue light exposure did not alter the attenuated response to OCT (Fig 3F), or impact OCT avoidance in naïve flies (Fig 3G). Similar results were obtained with control strains and sibling flies that carry, but do not express the ChR2 transgenes (Suppl Fig 3A-H). Therefore, the blue light used to photoactivate ChR2-expressing neurons is not itself a dishabituator.

Taken together with the outcome of silencing experiments, the results strongly support the conclusion that neurotransmission from the MBs is necessary and sufficient for cross-modal and intramodal dishabituation, without affecting OCT a voidance *per se*. Finally, the results support the hypothesis that dishabituators recruit the MBs and are in accord with evidence that these neurons process and gate responses to stimuli from different sensory modalities (Modi, Shuai et al. 2020, Vrontou, Groschner et al. 2021)

### Differential actions of KC subsets in olfactory habituation and dishabituation

Each of the two bilaterally symmetrical MBs consists of ∼990 αβ, ∼350 α΄β΄and ∼675 γ KCs (Crittenden, Skoulakis et al. 1998, Aso, Hattori et al. 2014). Neurons of each group are further subdivided into functional compartments (Aso, Hattori et al. 2014), which are differentially engaged in processes underlying olfactory learning and memory (Yu, Akalal et al. 2006, Krashes, Keene et al. 2007, Blum, Li et al. 2009, Zhang and Roman 2013, Aso, Hattori et al. 2014). Furthermore, KC subsets have been reported to play distinct roles in latency to habituate to OCT (Semelidou, Acevedo et al. 2018), as well as, in latency and habituation to recurrent footshocks (Acevedo, Froudarakis et al. 2007, Roussou, Papanikolopoulou et al. 2019, Foka, Georganta et al. 2022). Since KCs are necessary for dishabituation with footshock and YO (Fig 3), we wondered whether distinct KC subsets are differentially engaged by cross-modal and intramodal stimuli to drive the process.

To address this question, each KC subset was silenced conditionally via Shi^ts^, or photoactivated by Chr2 expression therein. Initially we determined that all driver heterozygotes presented with normal habituation and dishabituation both with footshocks and YO (Suppl Fig 4). Blocking neurotransmission from αβ neurons (Fig 4A), did not affect habituation or dishabituation either via footshock, or YO (Fig 4B). Surprisingly however, photostimulation of these KCs reinstated OCT aversion in habituated flies irrespective of light intensity (Fig 4C), but did not affect OCT avoidance in naïve animals (Fig 4D). Similar results were obtained with a second αβ KC driver, c739Gal4 (Suppl Fig 5B-D). These results indicate that αβ KCs are not necessary for dishabituation, likely because another KC subset is also able to drive the process. Alternatively, αβ neurons may serve a cooperative or supporting role to another KC subset in mediating dishabituation. This interpretation is supported by the robust αβ activation by photostimulation (Fig 4C), which whether cooperative or supportive, reinstated OCT avoidance, bypassing the activation state of other KCs potentially contributing to the process.

**Figure 4:**
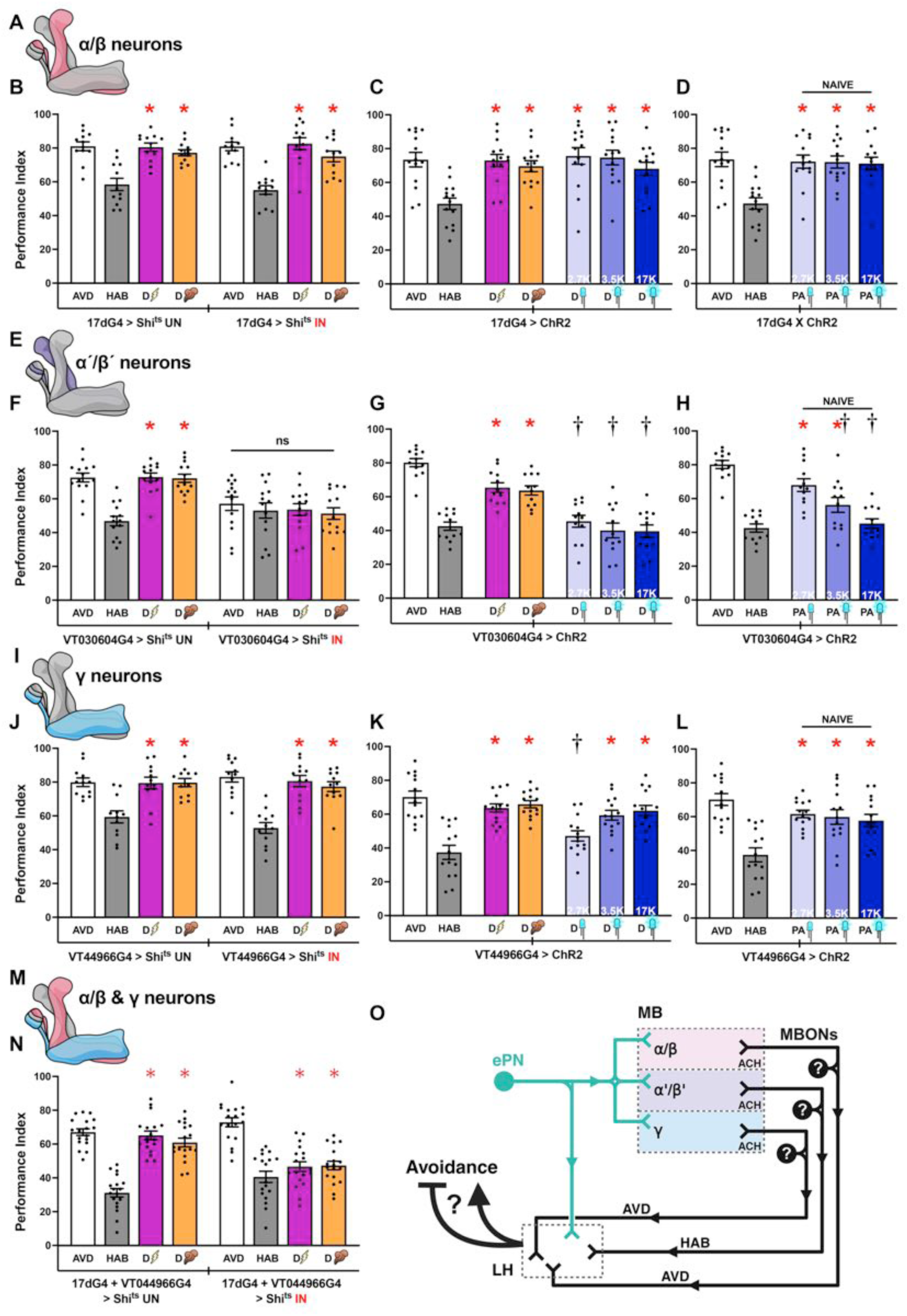
Distinct MBns regulate olfactory habituation and dishabituation. Except for the schematics, mean Performance Indices ± SEM are shown in all figures. The naïve response for OCT is shown in white bars, which are always significantly different from the habituated response (grey bars). Cross-modal (footshock) dishabituation is indicated by the bolt (magenta bar) and intramodal (YO), by the plume (orange bar). Exposure to blue light is indicated by the LED bulbs and increasing intensity (as indicated by the respective Klx) by deepening shades of blue. Stars in all figures indicate significant differences (p<0.005), from the habituated response and daggers significant difference (p<0.005) from naïve avoidance. All statistical details are presented in Supplemental Table 1. **A)** Schematic of αβ KCs highlighted in light red among the remaining KC types (gray). **B)** Functional silencing (IN) of αβ neurons with UAS-shibire^ts^ under 17dGal4 compared to control animals carrying, but not expressing the transgene at the permissive conditions (UN). Dishabituation with a 45V electric shock (magenta) and YO (orange), remained unaffected in both conditions. n=12. Additional yoked controls are presented in Supplemental Table 1. **C)** Photoactivation of αβ neurons with 3 light intensities in OCT-habituated flies restored avoidance of the odorant. n=14. **D)** Photoactivation of αβ neurons in naïve flies with 3 light intensities did not affect OCT avoidance. n=14. **E)** Schematic of α΄β΄ KCs highlighted in purple among the remaining KC types (gray). **F)** Functional silencing (IN) of α΄β΄ neurons with UAS-shibire^ts^ under VT030604Gal4 compared to control animals carrying but not expressing the transgene (UN). Silencing α΄β΄ MBns results in abrogation of habituation to OCT. Avoidance, habituation and dishabituation with a 45V electric shock (magenta) or YO (orange) were not significantly different. n=14. Additional controls run in parallel are presented in Supplemental Table 1. **G)** Photoactivation of α΄β΄ neurons under VT030604Gal4 in OCT-habituated flies with 3 light intensities did not restore avoidance, while a single footshock and YO exposure did. n=12. **H)** Photoactivation of α΄β΄ MBns in naïve flies resulted in light intensity dependent decrement in OCT avoidance. n=12. **I)** Schematic of γ KCs highlighted in blue among the remaining KC types (gray). **J)** Functional silencing (IN) of γ neurons with UAS-shibire^ts^ under VT044966Gal4 compared to control animals carrying, but not expressing the transgene (UN). Dishabituation with a 45V electric shock (magenta) and YO (orange), remained unaffected in both conditions. n=12. Additional controls run in parallel are presented in Supplemental Table 1. **K)** Photoactivation of γ neurons under VT044966Gal4 in OCT-habituated flies with 3 light intensities reinstated avoidance in light intensity dependent manner. n=14. **L)** Photoactivation of γ MBns in naïve flies did not affect OCT avoidance. n=14. **M)** Schematic of both αβ and γ KCs highlighted in light red and blue respectively. **N)** Functional silencing (IN) of αβ & γ neurons with UAS-shibire^ts^ under both 17dGal4 and VT044966Gal4 drivers compared to control animals carrying, but not expressing the transgene (UN). Dishabituation was abolished following both a 45 V electric shock (magenta) and YO (orange) stimulation. n=18. Additional controls run in parallel are presented in Supplemental Table 1. **O)** Schematic representation of the axonal projections of excitatory projection neurons (ePNs), transmitting olfactory information from the antennal lobes to the MB. Dishabituators (shock & YO) alter the sparse coding representation of the odor within the MB, leading to the activation of MBns. Although both α/β and γ neurons are required for encoding the novel stimulus, their activation is sufficient to drive dishabituation and reinstate the avoidance response. In contrast, activation of α΄β΄neurons is required for habituation and is sufficient to suppress/reduce avoidance to habituated levels, suggesting that these neurons act antagonistically to α/β and γ. Mushroom Body output neurons (MBONs) project to the Lateral Horn (LH), where they relay and distribute information processed in the MB.

In contrast to αβ KCs and in agreement with our previous results (Semelidou, Acevedo et al. 2018), silencing specifically their α΄β΄ counterparts under two different Gal4 drivers, VT030604Gal4 (Fig 4F) and c305aGal4 (Suppl. Fig 5F), attenuated avoidance, such that habituation and therefore dishabituation could not be assessed. To bypass this issue, we expressed ChR2 in α΄β΄ KCs, which did not interfere with habituation to OCT and dishabituation either by footshock or YO (Fig 4G and Suppl Fig 5G). Importantly, α΄β΄ KC photostimulation did not suffice to reinstate OCT avoidance in habituated flies (Fig 4G and Suppl Fig 5G), suggesting that these neurons do not contribute to the process. On the other hand, α΄β΄ photoactivation in naïve animals, especially at the higher intensities, reduced OCT avoidance, consistent with a role for these neurons in mediating habituation (Fig 4H and Suppl Fig 5H), rather than dishabituation.

Similar to results from αβ neurons, silencing γ KCs (Fig 4I) did not affect habituation to OCT, or footshock and YO-driven dishabituation (Fig 4J, Suppl Fig 5J). Conversely, their photoactivation in habituated flies reinstated OCT avoidance at the higher intensities (Fig 4K, Suppl Fig 5K), but did not affect avoidance in naïve flies (Fig 4L, Suppl Fig 5L). Taken together, the results suggest that neurotransmission from any single KC subset is not sufficient to drive dishabituation to 4 min-OCT exposure.

Since silencing all KCs impairs dishabituation and their photoactivation reinstates OCT avoidance (Fig 3) we hypothesized that αβ and γ KCs have redundant or cooperative roles, each sufficient to drive dishabituation, possibly with different activation thresholds (Fig 4C *versus* Fig 4K). To address this possibility directly, we generated flies harboring both the 17d and the VT044966 Gal4 drivers (Fig 4M). In agreement, with our hypothesis, simultaneous silencing of αβ and γ KCs blocked dishabituation with footshock and YO (Fig 4N), in effect emulating the effect of inhibiting pan-KC neurotransmission (Fig 3B). Collectively the results demonstrate that intramodal and cross-modal dishabituating stimuli converge on the MBs and mediate dishabituation via αβ and γ neurotransmission.

### The valence of intramodal and cross-modal dishabituators results in differential engagement of distinct Dopaminergic neurons (DANs)

How do dishabituating stimuli reach the MBs? Does the route depend on the valence of the dishabituator? DANs innervate the MBs, converging on specific MB compartments (Fig 5A) and convey reward or punishment information (Mao and Davis 2009, Waddell 2013, Aso, Hattori et al. 2014, Davis 2023). DANs also form axonoaxonic synapses with KCs, modulating their activities reciprocally (Cervantes-Sandoval, Phan et al. 2017). This is instructive on KC activity by changing the strength of particular KC synapses with their output neurons (MBONs) (Aso, Hattori et al. 2014), thus modulating the impact of stimuli. In addition, DANs receive inputs from MBONs (Dylla, Raiser et al. 2017), honing dynamic interactions with KCs such that errors in expected incoming stimuli can be predicted (Hancock, Rostami et al. 2022). Therefore, DANs are well-poised to play essential roles in dishabituation.

**Figure 5:**
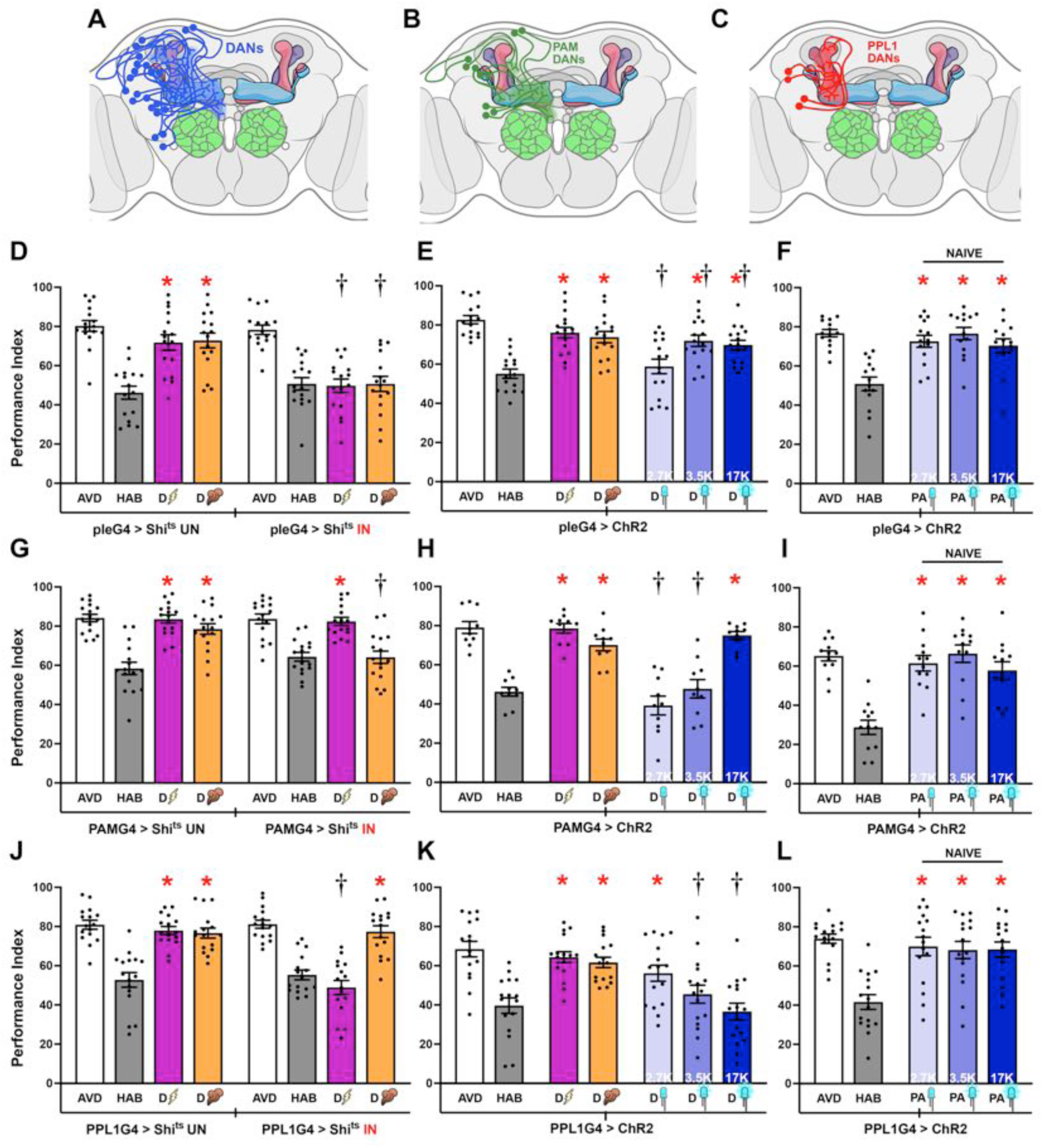
Dopaminergic Neurons drive cross-modal and intramodal dishabituation in Drosophila. Except for the schematics, mean Performance Indices ± SEM are shown in all figures. The naïve response for OCT is shown in white bars, which are always significantly different from the habituated response (grey bars). Cross-modal (footshock) dishabituation is indicated by the bolt (magenta bar) and intramodal (YO), by the plume (orange bar). Exposure to blue light is indicated by the LED bulbs and increasing intensity (as indicated by the respective Klx) by deepening shades of blue. Stars in all Figures indicate significant differences (p<0.005), from the habituated response and daggers significant difference (p<0.005) from naïve avoidance. All statistical details are presented in Supplemental Table 1. **A, B, C)** Schematics of all Dopaminergic neurons (DANs), the PAM-DAN cluster and the PPL1 DAN cluster, innervating the MB as indicated. **D)** Functional silencing of all DANs (IN), with UAS-shibire^ts^ under pleGal4 did not affect habituation after 4 min of OCT exposure, but dishabituation after stimulation with a 45V electric shock (magenta bar) and YO (orange bar) were impaired. At the permissive conditions (UN), habituation and both types of dishabituation were not affected. n=16. Additional yoked controls are presented in Supplemental Table 1. **E)** Photoactivation of all DANs in OCT-habituated flies with three different light intensities restored OCT avoidance in light intensity dependent manner. n=16 **F)** Photoactivation of all DANs in naïve flies without prior exposure to OCT did not affect OCT avoidance. n=14. **G)** Functional silencing (IN) of PAM-DANs with UAS-shibire^ts^ under 0273Gal4 was permissive to habituation after 4 min of OCT exposure and shock dishabituation (magenta bar), but not dishabituation by YO stimulation (orange bar). At the permissive conditions (UN) YO dishabituation was not affected. n=16. Additional yoked controls are presented in Supplemental Table 1. **H)** Photoactivation of PAM-DANs in OCT-habituated flies restored OCT avoidance only at the higher light intensity. n=10. **I)** Photoactivation of PAM-DANs in naïve flies without any exposure to OCT did not affect OCT avoidance. n=12. **J)** Functional silencing (IN) of PPL1-DANs with UAS-shibire^ts^ under MB504BGal4 did not affect habituation after 4 min of OCT exposure and dishabituation with YO (orange bar), but dishabituation by a 45 V electric shock stimulation (magenta bar) was impaired. At the permissive conditions (UN) shock dishabituation was not affected. n=16. Additional controls run in parallel are presented in Supplemental Table 1. **K)** Photoactivation of PPL1-DANs in OCT-habituated flies restored OCT avoidance only at the lower light intensity. n=16. **L)** Photoactivation of PPL1-DANs in naïve flies without any exposure to OCT did not affect OCT avoidance. n=16.

To determine whether DANs are involved in dishabituation, we initially conditionally silenced all dopaminergic neurons under the pan-DAN driver pleGal4 (Fig 5A). This manipulation did not affect avoidance or habituation to OCT, but both footshock and YO-driven dishabituation were disrupted (Fig 5D). Therefore, dopaminergic signals are critical for intramodal and cross-modal dishabituation. Conversely, photoactivating all DANs in habituated animals resulted restored OCT avoidance at the higher light intensities (Fig 5E), but did not alter OCT avoidance in naïve flies (Fig 5F). Habituation and dishabituation were also unaffected in yoked driver heterozygote controls (Suppl Fig 6A). Therefore, both footshock and YO dishabituators are relayed to the MBs by dopaminergic neurotransmission.

Because a 5-sec YO puff is attractive (Fig 2Ε) and a food related odor, we hypothesized that it might activate PAM neurons (Fig 5B), DANs known to transmit reward and positive valence signals to the MBs in conditioning tasks (Waddell 2013, Davis 2023). YO could activate PAM-DANs directly as an appetitive food-related cue (Lewis, Siju et al. 2015), or indirectly via the MBs or other sensory information integrating circuits (Cervantes-Sandoval, Phan et al. 2017, Boto, Stahl et al. 2019, Boto, Stahl et al. 2020). PAM-DANs can be effectively targeted using the 0273Gal4 driver (henceforth PAMGal4), which is expressed in all 130 neurons within this cluster (Burke, Huetteroth et al. 2012).

Conditional silencing of PAM-DANs did not affect avoidance or habituation to OCT and footshock mediated dishabituation, but specifically attenuated YO-mediated dishabituation (Fig 5G). In contrast, habituation and dishabituation were unaffected in driver and Shi^ts^ heterozygotes assayed in yoked experiments (Suppl Fig 6B). Interestingly, photoactivation of PAM-DANs in habituated flies restored OCT avoidance at the highest activation intensity (Fig 5H). The requirement for high-intensity activation to elicit this recovery further suggests that additional neural subsets may normally contribute to the process and are bypassed by direct strong PAM-DAN photoactivation. This increase in OCT avoidance in habituated animals emulated dishabituation as photoactivation at all intensities did not impact OCT avoidance in naïve animals (Fig 5I). These results indicate that PAM-DAN neurotransmission is necessary for intramodal dishabituation and suffices to promote recovery of the naïve osmotactic response. Therefore, MB neurotransmission from αβ and γ KCs driving intramodal dishabituation require instructive inputs from, or synergism with PAM-DAN activity to modulate MBON outputs (Aso, Sitaraman et al. 2014, Aso and Rubin 2016). In this scenario, silencing PAM-DANs is likely equivalent to attenuating MB output in response to the intramodal dishabituator.

Conversely, are the PPL1-DANs which convey the footshock stimuli for associative learning (Davis 2023), differentially involved in cross-modal dishabituation? To conditionally manipulate PPL1-DANs (Fig 5C), we used the MB504BGal4 driver (Aso, Hattori et al. 2014, Otto, Pleijzier et al. 2020), henceforth PPL1Gal4, which is expressed specifically in PPL1-γ1pedc, PPL1-γ2α′1, PPL1-α2α′2, and PPL1-α3 neurons (Feng, Weng et al. 2021). Attenuation of neurotransmission from these four types of PPL1 neurons did not affect avoidance or habituation to OCT, but selectively abrogated cross-modal dishabituation (Fig 5J).

Habituation and both types of dishabituation remained unaffected in driver and Shi^ts^ heterozygotes in parallel yoked experiments (Suppl Fig 6C). Surprisingly, photoactivation of PPL1-DANs in OCT-habituated animals resulted in full restoration of avoidance only at the lower photostimulation intensity (Fig 5K), but OCT avoidance in naïve animals remained unaffected (Fig 5L). The results demonstrate that PPL1-DAN activity is necessary and sufficient for dishabituation via the aversive cross modal footshock. This in turn suggests the instructive role of PPL1-DANs on the MBs regarding the novelty of the footshock dishabituator.

Do PPL1-DANs mediate dishabituation specific to the aversive cross-modal footshock dishabituator on OCT habituated animals, or are they also engaged by aversive intramodal dishabituators? To address this question, flies were habituated by 4-min exposure to the also aversive at this time scale YO and dishabituated either with a 3-sec OCT puff or a footshock. Silencing PPL1 neurons under MB504BGal4, suppressed dishabituation both with footshock and significantly also with OCT (Fig 6A). Conversely, photoactivation of PPL1-DANs partially reinstated YO avoidance at the lowest light intensity (Fig 6B). Interestingly however, silencing PAM-DANs did not affect dishabituation with OCT and footshock in YO-habituated flies (Fig 6C). These results indicate that PPL1 neurons convey aversive valence information to the MBs, irrespective of stimulus modality, this being necessary for aversive stimulus mediated intra- or cross-modal dishabituation. PPL1-DANs also receive excitatory inputs from KCs (Cervantes-Sandoval, Phan et al. 2017), but although they engage all three types of KCs, their inputs appear to augment or synergize with outputs specifically from the αβ and γ subsets to drive cross-modal dishabituation (Fig 6D).

**Figure 6:**
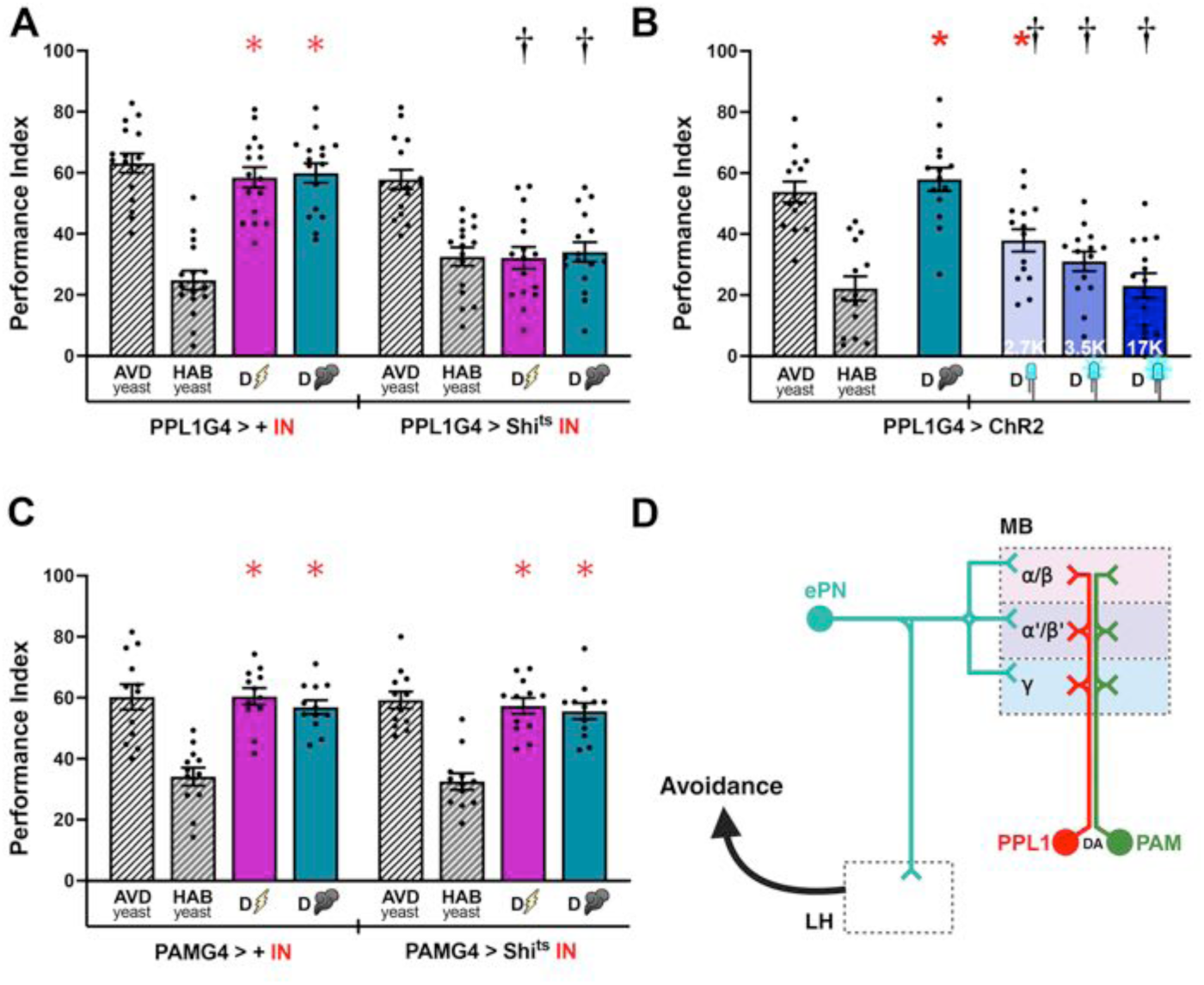
PPL1 Dopaminergic Neurons drive intra-modal and cross-modal dishabituation to yeast odor in Drosophila. Mean Performance Indices ± SEM are shown in all figures. The naïve response to yeast odor (YO) is shown in white striped bars, and is always significantly different from the habituated response (striped grey bars). The dishabituated response is shown in magenta for the footshock dishabituator, and in cyan for OCT. Exposure to blue light is indicated by the LED bulbs and increasing intensity (as indicated by the respective Klx) by deepening shades of blue. Stars in all Figures indicate significant differences (p<0.005), from the habituated response and daggers significant difference (p<0.005) from naïve avoidance. All statistical details are presented in Supplemental Table 1. **A)** Functional silencing (IN) of PPL1-DANs with UAS-shibire^ts^ under MB504BGal4 did not affect habituation after 4 min of YO exposure, but dishabituation by a 45 V electric shock stimulation (magenta bar) and OCT (cyan bar), were impaired. n=16. **B)** Flies expressing ChR2XXL in their MBs avoid and habituate to YO, and dishabituate with OCT (cyan bar) normally. Photoactivation of PPL1-DANs in YO-habituated flies restored YO avoidance only at the lower light intensity. n=14. **C)** Functional silencing (IN) of PAM-DANs with UAS-shibire^ts^ under 0273Gal4 was permissive to habituation after 4 min of YO exposure and dishabituation to both shock (magenta bar) and OCT (cyan bar). n=12. **D)** Schematic representation of the axonal projections of ePNs and dopaminergic neurons (PPL1 & PAM) to the MB. Dopaminergic reinforcement relays information about a novel stimulus and its associated valence, depending on the specific neuronal subset activated.

The collective results indicate that the novelty and the valence of the dishabituators is transmitted to the MB via DANs irrespective of their modality. Both aversive dishabituators engage the PPL1 neurons, whereas the intramodal but attractive YO engages PAM-DANs. We hypothesize that robust PPL1 signals to the MBs may lead to synaptic depression as described for negatively reinforced associative learning (Hige, Aso et al. 2015, Perisse, Owald et al. 2016), a condition that could suppress dishabituation as suggested by their robust photo activation (Fig 5K). Alternatively, distinct neurons within the PPL1 cluster could promote dishabituation, while others with higher activation thresholds may suppress it. Such PPL1 neurons may project to different MB compartments (Aso, Siwanowicz et al. 2010, Aso, Hattori et al. 2014), facilitating circuit-level regulation.

### Differential engagement of neurons within the PPL1 cluster in dishabituation

The PPL1-DANs are functionally diverse (Sareen, McCurdy et al. 2021) and modulate behavioral outputs in a context-specific manner. To address the abovementioned hypothesis and investigate whether distinct PPL1 subpopulations underlie the intensity-dependent effects of PPL1 photoactivation on dishabituation (Fig 5K), we capitalized on the specificity of extant Gal4 drivers that selectively target individual PPL1 compartments (Feng, Weng et al. 2021). Activation of γ2α΄1 and γ1pedc PPL1-DANs was shown to suppress avoidance in naïve flies (Aso, Sitaraman et al. 2014, Perisse, Owald et al. 2016) and promote associative memory, while neurotransmission from α2α΄2 and α3 PPL1-DANs to interfere with memory consolidation (Feng, Weng et al. 2021). Because the two sets of PPL1-DANs apparently promote different outcomes, but both arborize on the α and γ KCs, which contribute to dishabituation (Fig 4B, J, N), we investigated whether activation of one or more of these DANs suffices to drive aversive stimulus-mediated dishabituation.

Each DAN subset was photostimulated in naïve and OCT-habituated flies under their respective specific Gal4 drivers, MB630B, MB058B, MB320C and MB296B (Feng, Weng et al. 2021). Consistent with the results obtained using the broader PPL1 driver MB504BGal4, post-habituation photoactivation of PPL1-γ2α΄1 (Fig 7A) and to a lesser extend PPL1-γ1pedc neurons (Fig 7D), resulted in significant recovery of OCT avoidance (Fig 7B, E), especially at the lower intensity. In contrast, photostimulation of α2α΄2 (Fig 7G) and α3 (Fig 7J) PPL1-DANs in habituated animals did not alter their attenuated OCT avoidance at any intensity (Fig 7H, K). Therefore, neurotransmission from γ2α΄1 and γ1pedc, but not α2α΄2 and α3 PPL1-DANs to KCs is required for dishabituation with footshock and likely other aversive dishabituators. The proximity of α2α΄2 and α3 with γ2α΄1 and γ1pedc PPL1 neurons (Feng, Weng et al. 2021), suggests that these results are unlikely due to inefficient photoactivation of the former due to their location in the CNS.

**Figure 7:**
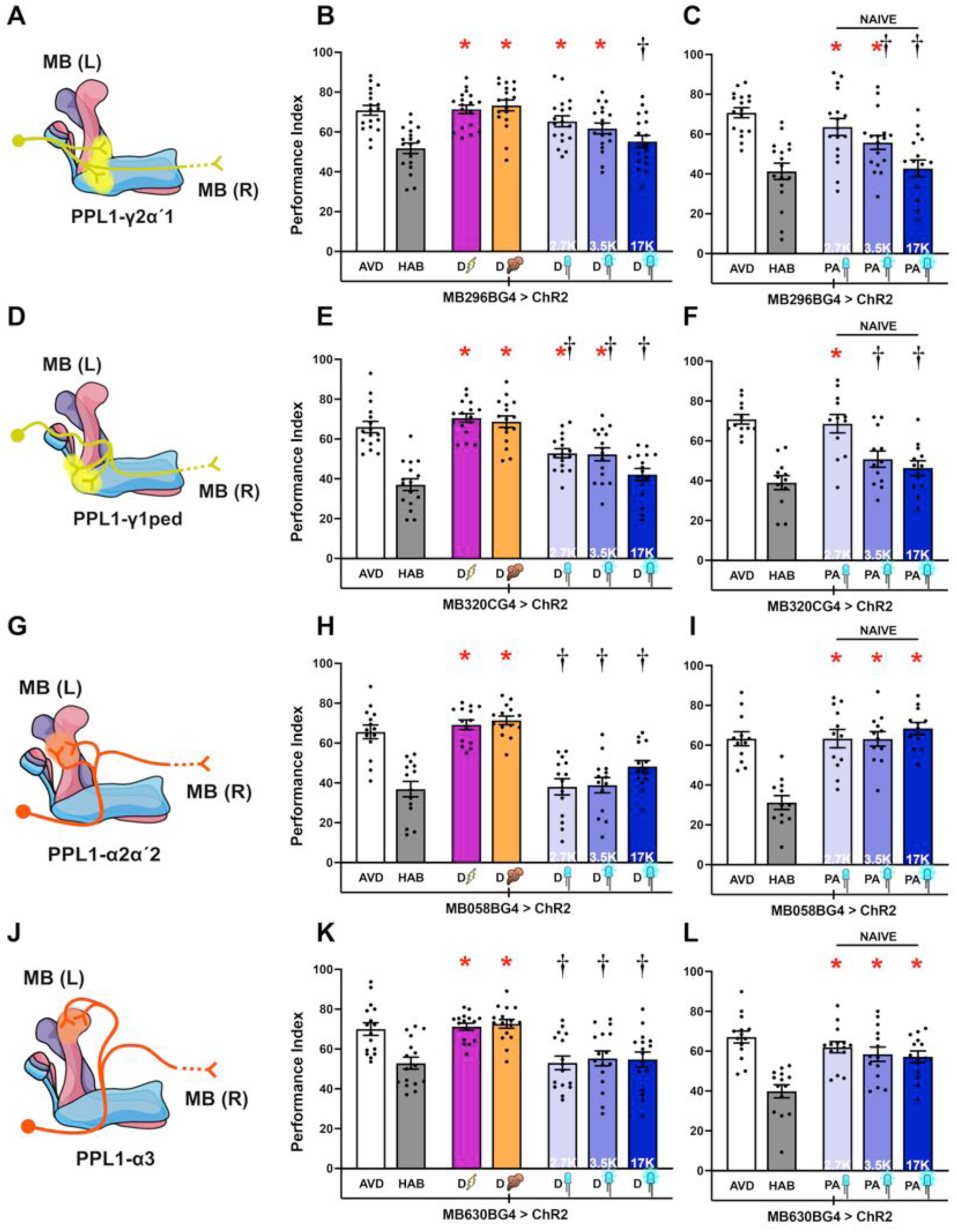
The role of PPL1 neuronal subsets in cross-modal dishabituation. Mean Performance Indices ± SEM are shown in all figures. The naïve response for OCT is shown in white bars and is always significantly different from the habituated response (grey bars). Cross-modal (footshock) dishabituation is indicated by the bolt (magenta bar) and intramodal (YO), by the plume (orange bar). Exposure to blue light is indicated by the LED bulbs and increasing intensity (as indicated by the respective Klx) by deepening shades of blue. Stars in all Figures indicate significant differences (p<0.005), from the habituated response and daggers significant difference (p<0.005) from naïve avoidance. All statistical details are presented in Supplemental Table 1. **A, D)** Schematics of PPL1-γ2α΄1 and PPL1-γ1pedc neurons and their ipsilateral (MB L) and contralateral (MB R) projections. **B)** Photoactivation of PPL1-γ2α΄1 neurons under MB296BGal4 in OCT-habituated flies restored OCT avoidance at the low, but not the stronger light intensities (p=0.0087 *versus* the habituated response and p=0.0162 *versus* naïve avoidance). n=18 **C)** Photoactivation of PPL1-γ2α΄1 neurons in naïve flies resulted in light intensity-dependent decrement in OCT avoidance from naïve avoidance (p=0.0064 at 3.5Klx and p<0.0001 at 17Klx). n=17. **E)** Photoactivation of PPL1-γ1pedc neurons under MB320CGal4 in OCT-habituated flies restored OCT avoidance at the low and strong light intensities respectively. n=16. **F)** Photoactivation of PPL1-γ1pedc neurons in naïve flies resulted in light intensity-dependent decrement in OCT avoidance with performance at naïve levels only at 2.7K (p=0.6747). n=12. **G, J)** Schematics of PPL1-α2-α΄2 and PPL1-α3 neurons and their ipsilateral (MB L) and contralateral (MB R) projections. **H)** Photoactivation of PPL1-α2-α΄2 neurons under MB058BGal4 in OCT-habituated flies with 3 light intensities did not restore avoidance, while the response recovered after a single footshock and YO exposure. n=14. **I)** Photoactivation of PPL1-α2-α΄2 neurons in naïve flies with 3 light intensities did not affect OCT avoidance. n=12. **K)** Photoactivation PPL1-α3 neurons under MB630BGal4 in OCT-habituated flies with 3 light intensities did not restore avoidance, while the response recovered after a single footshock and YO exposure. n=16. **L)** Photoactivation of PPL1-α3 neurons in naïve flies with 3 light intensities did not affect OCT avoidance. n=14.

Interestingly, the γ2α΄1 and γ1pedc PPL1 neurons provide aversive reinforcement to γ KCs (Aso and Rubin 2016, Li, Lindsey et al. 2020). Indeed, photoactivation of γ2α΄1 and γ1pedc neurons in naïve flies suppressed OCT avoidance, especially at the higher, but not the lower light intensities (Fig 7C, F). In contrast, photostimulation of α2α΄2 and α3 PPL1-DANs in naïve animals did not affect OCT avoidance (Fig 7I, L).

Taken together, these results show that activation of γ2α΄1 and γ1pedc PPL1-DANs mediate dishabituation and suggest that dopamine released by these DANs modulates behavior in a prior experience-dependent manner (naïve *versus* habituated flies) by differential suppression of KC outputs mediating the habituated response. This implicates KC outputs onto aversion-driving MBONs leading to decreased avoidance (Aso, Sitaraman et al. 2014, Perisse, Owald et al. 2016). This situation is akin to suppression of avoidance in hungry flies, enabling conditioned approach to food upon PPL1-DAN activation (Perisse, Owald et al. 2016).

The differential effects of γ2α΄1, γ1pedc and α2α΄2, α3 PPL1-DANs on OCT avoidance and aversive stimulus-mediated dishabituation also indicate that photostimulation of neurons impinging on the MBs alone is not sufficient to drive dishabituation and underlies the specificity of our photoactivation approach. Furthermore, the arborizations of γ2α΄1, γ1pedc onto the KCs (Fig 7A, D) and the similar response profile upon photoactivation of the broader PPL1 cluster (Fig 5K), is consistent with the proposed importance of inputs to γ neurons for dishabituation from OCT.

### APL neurons regulate intramodal dishabituation through GABAergic neurotransmission

While DANs innervate specific compartments, additional afferent neurons impinge upon the MBs (Keene, Krashes et al. 2006, Liu and Davis 2009, Chen, Wu et al. 2012). We hypothesized that these inputs act synergistically with dopaminergic signals to fine-tune and modulate MB activity. Therefore, we asked whether additional neuronal subsets that ramify onto the MBs contribute to intramodal and cross-modal dishabituation. Preliminary screens demonstrated that inhibition of Dorsal Anterior Lateral (DAL) and Dorsal Paired Medial (DPM) neurons did not affect habituation to OCT or dishabituation (not shown). However, silencing the Anterior Paired Lateral (APL) interneurons (Fig 8A), impaired intramodal dishabituation without affecting OCT avoidance and habituation, or cross-modal dishabituation (Fig 8B).

**Figure 8:**
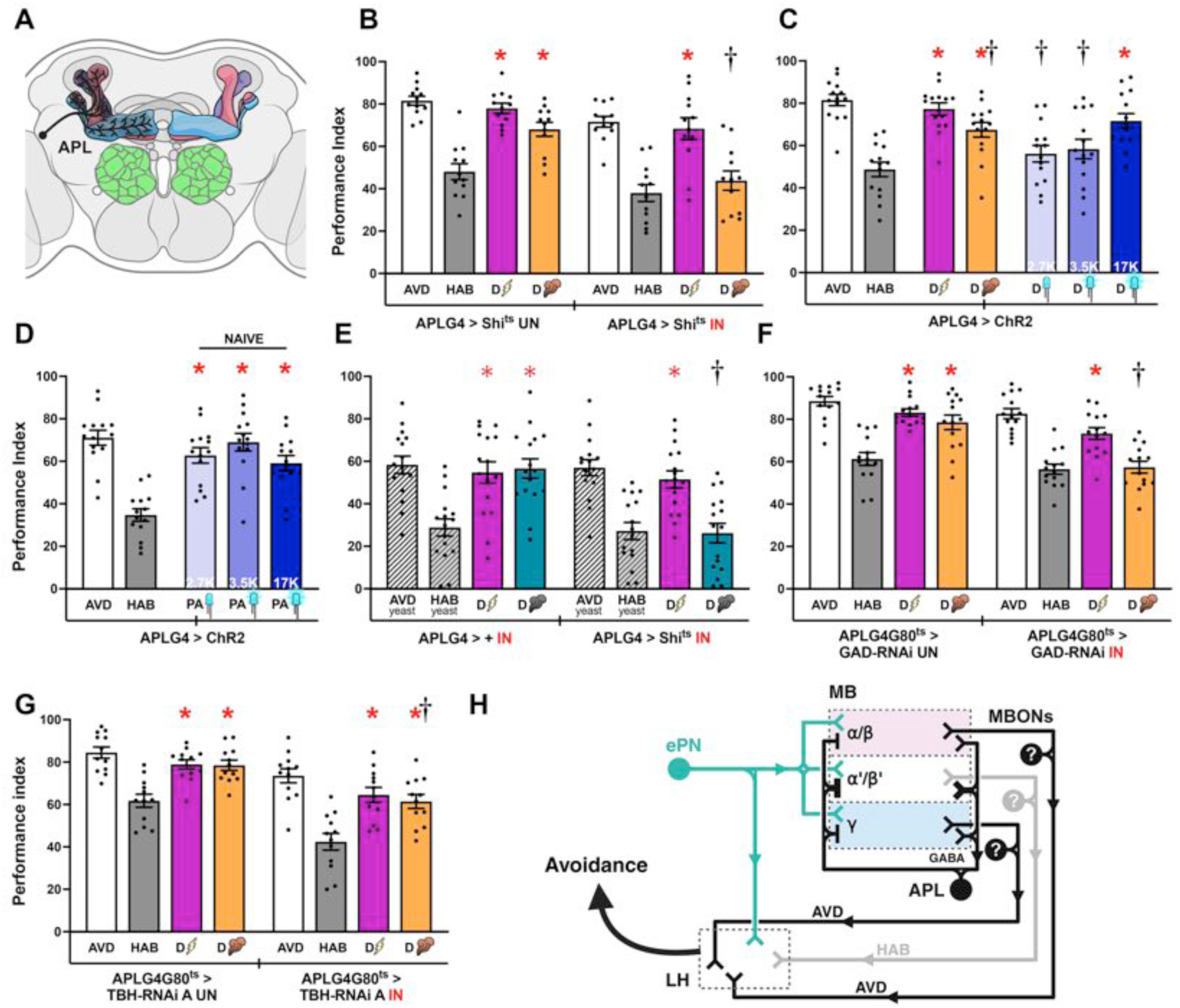
APL neurons regulate intramodal Dishabituation. Mean Performance Indices ± SEM are shown in all figures. The naïve avoidance response (AVD) is shown by the white bar for OCT and white striped bar for yeast odor (YO). The habituated response (HAB) is shown by the grey bars for OCT and the striped, grey bar for YO. The dishabituated response is shown in magenta for the footshock dishabituator, in orange for YO and in cyan for OCT. Exposure to blue light is indicated by the LED bulbs and increasing intensity (as indicated by the lx) by deepening shades of blue. Stars in all Figures indicate significant differences (p<0.005), from the habituated response and daggers significant difference (p<0.005) from naïve avoidance. All statistical details are presented in Supplemental Table 1. **A)** Schematic of APL neurons innervating the MB. **B)** Functional silencing of APL neurons with UAS-shibire^ts^ under APLGal4 was permissive to habituation after 4 min OCT exposure and shock dishabituation (magenta bar), but dishabituation by YO exposure (orange bar) was impaired. At the permissive conditions (UN), YO dishabituation was not affected. n=12. Additional controls run in parallel are presented in Supplemental Table 1. **C)** Photoactivation of APL neurons in OCT-habituated flies restored OCT avoidance only at the higher light intensity. n=14. **D)** Photoactivation of APL neurons in naïve flies with 3 light intensities did not affect OCT avoidance. n=14. **E)** Functional silencing of APL neurons with UAS-shibire^ts^ under APLGal4 was permissive to habituation after a 4 min YO exposure and shock dishabituation (magenta bar), but dishabituation following OCT exposure (cyan bar) was impaired. n=16. **F)** Abrogation of the GABAergic output of APL neurons with UAS-GAD-RNAi under APLGal4;Gal80^ts^ at 30°C did not affect habituation after 4 min OCT exposure or dishabituation with shock (magenta bar), but dishabituation by YO stimulation was impaired (orange bar). At the permissive conditions (UN), dishabituation with YO was not affected. n=14. **G)** Abrogation of the octopaminergic output of APL neurons via UAS-TBH-RNAi under APLGal4;Gal80^ts^ at 30°C, compared to control animals carrying, but not expressing the transgene (UN). Dishabituation with a 45V electric shock (magenta) and YO (orange), although partially reduced, was sufficient in both conditions. n= 12. **H)** Schematic representation of the dendritic inputs from the MB and axonal outputs to the MB of GABAergic APL neuron. Based on the Drosophila connectome and calculated synapses with the MB, APL forms a feedforward inhibitory loop. APL silencing selectively abrogates intramodal dishabituation, whereas habituation remains intact. This suggests that persistent activity of α′/β′ neurons, resulting from the loss of APL-mediated inhibition, continues to promote habituation. Concurrently, disrupted signaling to α/β and γ MBns, may compromise the encoding of information about novel odors, thereby preventing dishabituation.

The two bilaterally symmetrical APL neurons (APLNs) innervate broadly the MBs (Fig 8A), engaging all KC subsets. APLNs, form inhibitory reciprocal synapses targeting both projection neuron ramifications onto KC dendrites themselves and are thought to normalize odor-evoked representations therein (Driscoll, Buchert et al. 2021) by localized feedback inhibition, promoting the sparseness and decorrelation of olfactory representations (Lin, Bygrave et al. 2014, Amin, Apostolopoulou et al. 2020). Their activity regulates odor-evoked neurotransmission from the MBs (Prisco, Deimel et al. 2021). Given their critical modulatory role on olfactory inputs and MB outputs, it is not surprising that silencing APLNs impaired specifically YO-mediated intramodal dishabituation (Fig 8B).

To verify the results of the silencing experiments and determine whether APLN activity suffices to drive dishabituation, we photostimulated these neurons in habituated flies expressing ChR2 therein. Activation of APLNs restored OCT avoidance in OCT-habituated flies, but only at the highest light intensity (Fig 8C), without affecting naïve osmotactic responses (Fig 8D). Control driver heterozygotes performed normally (Suppl Fig 7A). This activation profile suggests either that the APLN activation threshold to drive intramodal dishabituation is high, or it requires synergism with additional MB-impinging neurons, such as the PAM-DANs, which present a similar activation profile (Fig 5H). This putative requirement for synergism is likely obviated with intense photostimulation, but nevertheless APLNs are necessary and sufficient for YO-mediated dishabituation.

Because the YO dishabituator also engages the PAM-DANs (FIG 5G, H), we wondered whether APLNs are engaged in intramodal dishabituation mediated by attractive stimuli, or even if the intramodal dishabituator is aversive. Hence, we habituated flies to YO and dishabituated with the PPL1-engaging (Fig 6A) aversive OCT. Significantly, silencing APLNs completely blocked OCT-mediated dishabituation in YO-habituated flies, whereas footshock-mediated dishabituation remained unaffected (Fig 8E). Therefore, neurotransmission from APLNs is required for intramodal dishabituation irrespective of the valence of the dishabituating odorant.

Interestingly, connectomic evidence indicates that APL neurons receive sparse inputs from PAM and PPL1 DANs (Takemura, Aso et al. 2017, Li, Lindsey et al. 2020). In addition, APL-driven inhibition is localized within MB compartments defined by DAN innervation (Amin, Apostolopoulou et al. 2020). Therefore, APLN activity can be modulated by incoming dopaminergic stimulation and in turn it modulates MB output (Amin, Apostolopoulou et al. 2020, Prisco, Deimel et al. 2021).

APLNs are reported to be both GABAergic and Octopaminergic (Wu, Shih et al. 2013), we asked which of the two neurotransmitters mediates the contribution of these neurons to intramodal dishabituation. Therefore, we used RNA interference to downregulate the rate-limiting enzymes in Octopamine and GABA biosynthesis within APLNs specifically in adult flies. Limiting GABA synthesis by attenuating of glutamic acid decarboxylase (Gad) under APLGal4;Gal80^ts^, did not affect OCT-habituation, but impaired YO-mediated dishabituation, whereas footshock dishabituation was unaffected (Fig 8F). Driver and RNAi-mediating transgene heterozygotes did not present with avoidance, habituation or dishabituation deficits in yoked experiments (Suppl Fig 7B, C). In contrast, downregulation of tyramine beta-hydroxylase (TBH) required for Octopamine biosynthesis, did not affect OCT avoidance and habituation or have a major impact on dishabituation (Fig 8G). The results were qualitatively recapitulated with a second independent TBH RNAi, presenting again a minor effect on YO-driven dishabituation (Suppl Fig 7E and Suppl Fig 7D for control heterozygotes). Therefore, APLN octopaminergic neurotransmission does not seem to contribute to dishabituation.

Collectively, the results demonstrate that APLNs mediate both appetitive and aversive intramodal dishabituation through GABAergic inhibition of the KCs suggesting that APL-mediated local inhibition on KCs (Amin, Apostolopoulou et al. 2020, Prisco, Deimel et al. 2021) is essential for intramodal dishabituation. As schematized in Figure 8H, we posit that this GABAergic activity is required to inhibit the activity at least of α΄β΄ KCs, which drive OCT habituation (Fig 4F, G). This notion is supported by connectome data indicating increased number of synapses from APLNs to α΄β΄ KCs (Suppl Fig 7H) and is represented in the schematic (Fig 8H) with heavier lines.

### MBONs are engaged in intramodal habituation and dishabituation

Because APL-driven GABAergic inhibition of the MBs is necessary for intramodal dishabituation (Fig 8F) and the role of efferent GABAergic innervation on MB function (Masuda-Nakagawa, Ito et al. 2014, Driscoll, Buchert et al. 2021, Georganta, Moressis et al. 2021), we investigated whether additional GABAergic neurons contribute to dishabituation. However, silencing of all GABAergic neurons under GADGal4 resulted in nearly paralyzed animals (not shown), we opted to ask whether GABAergic neurotransmission is implicated in dishabituation by photoactivating these neurons. Indeed, activation of all GABAergic neurons, in OCT-habituated animals resulted in dishabituation at the lowest light intensity (Suppl Fig 7F). In contrast, strong photostimulation in OCT-habituated flies had the opposite effect, resulting in decreased OCT avoidance to levels as low as the habituated response (Suppl Fig 7F). Accordingly, intense pan-GABAergic photoactivation in naïve flies also resulted in significantly reduced OCT avoidance (Suppl Fig 7G). These results indicate that in fact GABAergic activity is necessary for dishabituation promoting recovery of OCT avoidance. However, robust GABAergic activity emulated by the intense photostimulation suppresses OCT avoidance, potentially reflecting its the proposed and described role for inhibitory signaling in habituation (Ramaswami 2014, Semelidou, Acevedo et al. 2018).

OCT dishabituation at low light intensity, but suppressed avoidance upon intense photostimulation is consistent with the notion that high affinity GABAergic receptors drive dishabituation, while low affinity GABA receptors appear involved in attenuated odor avoidance. Both classes of GABA receptors have been described in Drosophila (Rosario, Barat et al. 1989) and in the MBs receive GABAergic signals from the APL (Liu, Krause et al. 2007, Ren, Li et al. 2012). If GABAergic neurotransmission to the MBs is necessary for dishabituation, then this activity must be inhibited to mediate habituation. This could occur either by KCs directly inhibiting the inhibitory APLN activity via their numerous reciprocal synapses (Suppl Fig 7H), or via MBONs impinging upon the APL. Therefore, we searched for MBONs receiving inputs from KCs and impinging upon APLNs.

MBON11 (Fig 9A), is a GABAergic feedforward neuron, receiving inputs from γ and via the pedunculus also from αβ KCs and arborises back on αβ neurons, the contralateral MB and the APL (Aso, Hattori et al. 2014, Aso, Sitaraman et al. 2014, Takemura, Aso et al. 2017, Li, Lindsey et al. 2020). Given the role of GABAergic neurotransmission in dishabituation and the role of αβ and γ KCs in this process (Fig 4N, O), we sought to determine the role of MBON11 in dishabituation.

**Figure 9:**
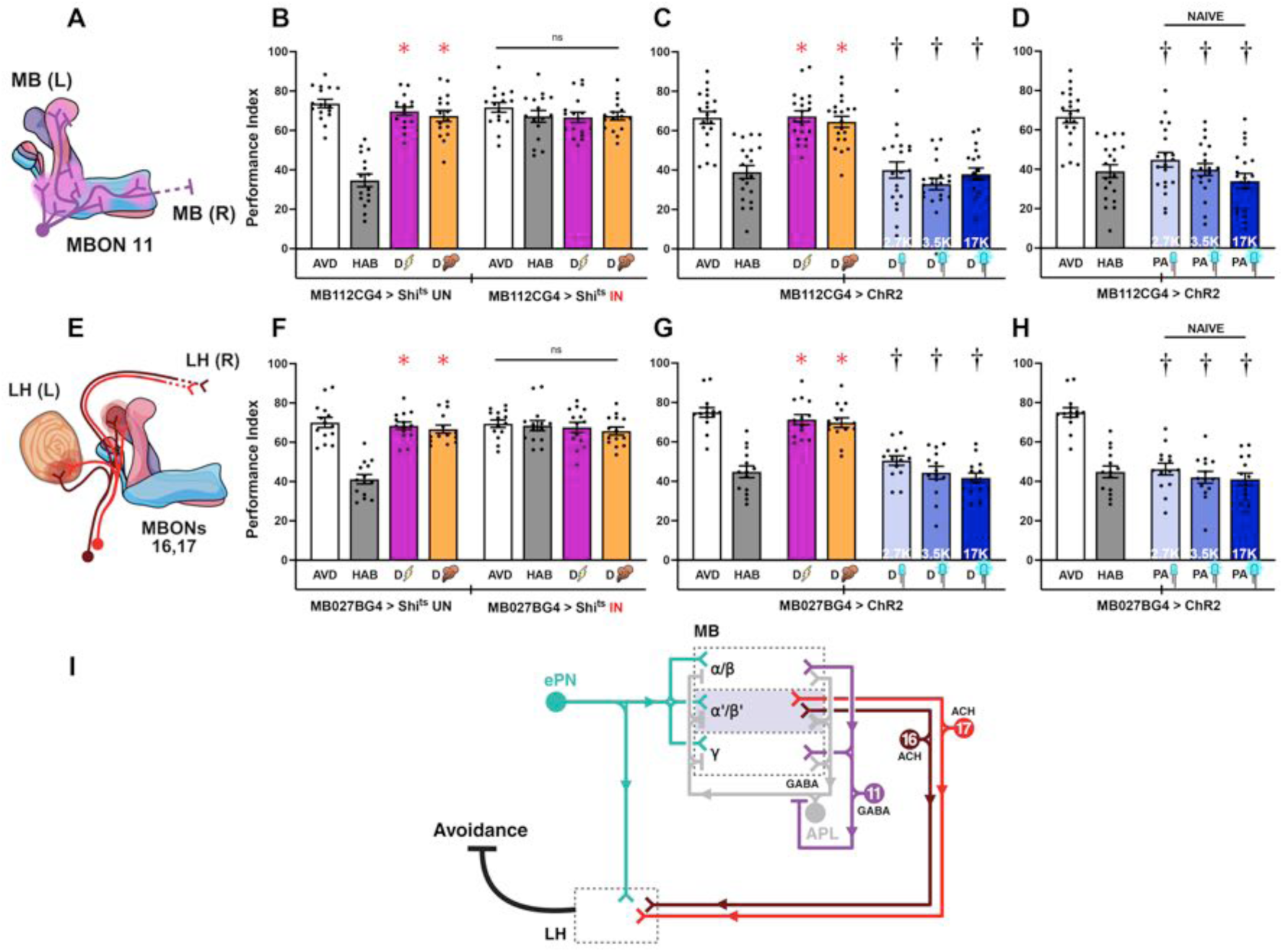
MBONs mediating olfactory habituation. Except for the schematics, mean Performance Indices ± SEM are shown in all figures. The naïve response for OCT is shown in white bars and is always significantly different from the habituated response (grey bars). Cross-modal (footshock) dishabituation is indicated by the bolt (magenta bar) and intramodal (YO), by the plume (orange bar). Exposure to blue light is indicated by the LED bulbs and increasing intensity (as indicated by the respective Klx) by deepening shades of blue. Stars in all Figures indicate significant differences (p<0.005), from the habituated response and daggers significant difference (p<0.005) from naïve avoidance. All statistical details are presented in Supplemental Table 1. **A)** Schematic of the ipsilateral (MB L) and contralateral (MB R) projections of MBON11 highlighted in violet. **B)** Functional silencing (IN) of MBON11 with UAS-shibire^ts^ under MB112CGal4 compared to control animals carrying, but not expressing the transgene (UN). Silencing MBON11 results in abrogation of habituation to OCT. However, avoidance, habituation and dishabituation with a 45V electric shock (magenta) or YO (orange), were not significantly different. n=16. Additional controls run in parallel are presented in Supplemental Table 1. **C)** Photoactivation of MBON11 in OCT-habituated flies with 3 light intensities did not restore avoidance, while a single footshock and YO exposure did. n=20. **D)** Photoactivation of MBON11 in naïve flies reduced OCT avoidance across all tested light intensities. n=20. **E)** Schematic of the ipsilateral (MB L) and contralateral (MB R) axonal projections of MBONs 16, 17 highlighted in maroon and scarlet. **F)** Functional silencing (IN) of MBONs 16, 17 with UAS-shibire^ts^ under MB027BGal4 compared to control animals carrying, but not expressing the transgene (UN). Silencing MBONs 16, 17 results in abrogation of habituation to OCT. Avoidance, habituation and dishabituation with a 45V electric shock (magenta) or YO (orange) were not significantly different. n=14. Additional controls run in parallel are presented in Supplemental Table 1. **G)** Photoactivation of MBONs 16, 17 in OCT-habituated flies with 3 light intensities did not restore avoidance, while a single footshock and YO exposure did. n=14. **H)** Photoactivation of MBONs 16, 17 in naïve flies reduced OCT avoidance across all tested light intensities. n=14. **I)** Schematic representation of the axonal projections of ePNs, which transmit olfactory information from the antennal lobes to the MB and MBONs relaying that information to LH to promote habituation. Activation of either α/β or γ KCs promote avoidance, reinstating the naïve response. GABAergic MBON 11 arborizes in α/β and γ KC axons where it receives excitatory cholinergic input. Silencing MBON 11 results in loss of habituation. Based on connectomic analyses showing that the APL neuron is the primary postsynaptic target of MBON11, it is possible that over a threshold activation of MBON11 by α/β and/or γ MBns inhibits APL activity. This, in turn, relieves APL-mediated inhibition within the Mushroom Body, particularly of α′/β′ neurons, thereby promoting habituation. Cholinergic MBONs 16 and 17 are also required for habituation, as their silencing abolishes this behavioral response. Their arborization within the α′/β′ lobes and projections to the LH position them as a critical link between the MB and the LH for the transmission of habituation-related signals

Silencing MBON11 via its specific MB112CGal4 driver eliminated habituation to OCT occluding dishabituation assessment of (Fig 9B). Importantly, MBON11 photoactivation in habituated flies did not restore OCT avoidance at any light intensity (Fig 9C), while its activation in naïve flies resulted in diminished OCT avoidance (Fig 9D). Control driver heterozygotes habituated and dishabituated efficiently with YO or footshocks as expected (Suppl Fig 8A). Therefore, MBON11 is necessary to modulate OCT avoidance and habituation, not dishabituation. Because it arborizes onto the APL, we hypothesize that MBON11 activation from αβ and γ neurons inputs inhibits APL activity. Because APLNs target preferentially α΄β΄ KCs (Suppl Fig 7H -Top output partners) and inhibit their habituation promoting activity negating this inhibition via MBON11 activation is necessary for habituation. Because MBON11 outputs also feedback onto αβ and γ KCs it also attenuates their activity. Therefre, it appears that MBON11-mediated inhibition of APLNs relieves its blockade of α΄β΄ activity, which combined with attenuation of the avoidance/dishabituation-promoting αβ and γ activity drives habituation to OCT.

To strengthen the hypothesis regarding the role of MBON11 in relation to disinhibition of α΄β΄ KC activity underlying olfactory habituation, we manipulated MBONs originating in α΄β΄ KCs and arborizing onto the LH. As an example, we chose driver MB027B Gal4 (Aso, Hattori et al. 2014), which marks the cholinergic MBONs 16 (α′3ap) and 17 (α′3m) (Fig 9E). Silencing MBONs 16 and 17 eliminated habituation, but did not affect OCT avoidance (Fig 9 F), as did silencing α΄and β΄KCs (Fig 4F). Conversely, photoactivation of these MBONs in habituated animals attenuated dishabituation at all light intensities (Fig 9G). However, photoactivation of these neurons in naïve flies suppressed OCT avoidance to levels exhibited by habituated animals. Hence robust cholinergic neurotransmission from α΄ KCs and most probably and their β΄ counterparts, to the LH mediates habituation to OCT. Control heterozygotes performed as expected (Supl Fig 8B). These results were strengthened further by silencing MBONs 16 and 17 with a different driver, MB543B Gal4, which also completely blocked habituation (Suppl Fig 8C).

Collectively these results support the model presented in Fig 9I, postulating that APLN GABAergic activity consequent to OCT exposure and most likely via reciprocal synapses with KCs suppresses α΄β΄ activity, promoting odor avoidance or dishabituation in habituated animals. MBON11-mediated GABAergic inhibition of the APLN inhibitory activity releases α΄β΄ neurotransmission, which via cholinergic MBONs activates LHNs mediating decreased avoidance of the odor after 4 min of exposure (Fig 9I).

Conversely from the habituation-promoting activity of MBON11 are additional MBONS projecting from αβ and/or γ KCs to the LH engaged in mediating OCT avoidance/dishabituation? The dendrites of the cholinergic MBON18, marked by driver MB050B (Fig 10A) arborize on α2 KCs and project to the LH (Aso, Sitaraman et al. 2014, Takemura, Aso et al. 2017, Li, Lindsey et al. 2020). Photoactivation of MBON 18 in OCT-habituated animals restored OCT avoidance especially at the higher intensities (Fig 10B), but did not impact naïve osmotaxis to that odor (Fig 10C). A caveat that may be reflected in the requirement for more intense activation to drive dishabituation is that MB050B also marks, albeit lightly ΜΒΟΝ α΄1, whose photoactivation may antagonize the effects of MBON18 neurotransmission to the LH in our model. Therefore, activation of the α2-efferent MBON18 is sufficient to induce dishabituation.

**Figure 10:**
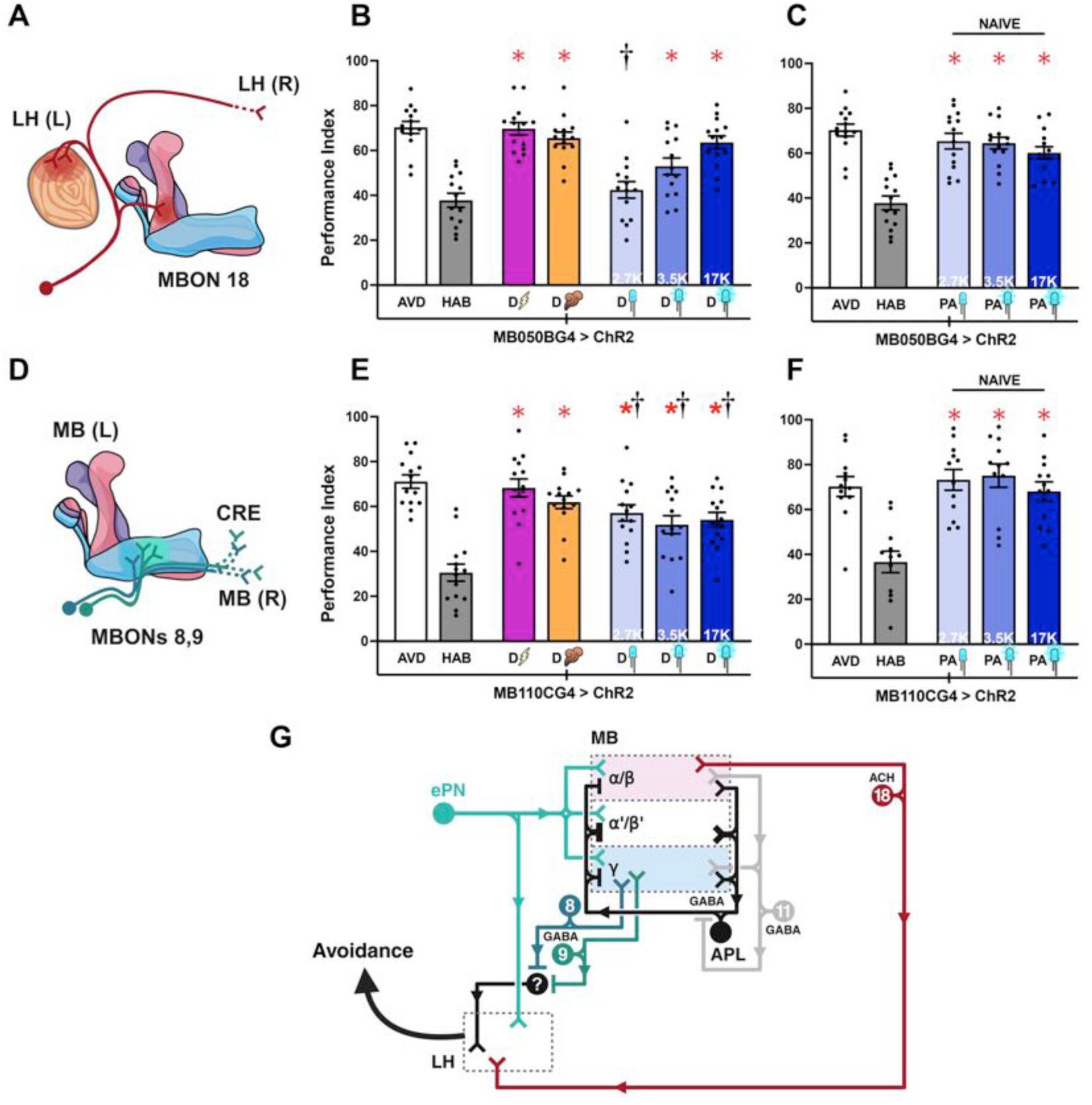
MBONs implicated in intramodal dishabituation. Except for the schematics, mean Performance Indices ± SEM are shown in all figures. The naïve response for OCT is shown in white bars and is always significantly different from the habituated response (grey bars). Cross-modal (footshock) dishabituation is indicated by the bolt (magenta bar) and intramodal (YO), by the plume (orange bar). Exposure to blue light is indicated by the LED bulbs and increasing intensity (as indicated by the respective Klx) by deepening shades of blue. Stars in all Figures indicate significant differences (p<0.005), from the habituated response and daggers significant difference (p<0.005) from naïve avoidance. All statistical details are presented in Supplemental Table 1. **A)** Schematic of the axonal projections of MBON 18 highlighted in burgundy. **B)** Photoactivation of MBON18 in OCT-habituated flies restored OCT avoidance in light intensity dependent manner. n=14. **C)** Photoactivation of MBON 18 in naïve flies without any exposure to OCT did not affect OCT avoidance. n=14. **D)** Schematic of the axonal projections of MBONs 08, 09 highlighted in blue-teal and green-teal. **E)** Photoactivation of MBONs 08, 09 under the MB110CGal4 driver in OCT-habituated flies restored OCT avoidance at all light intensities. n=14. **F)** Photoactivation of MBON 08, 09 in naïve flies did not affect OCT avoidance. n=12. **G)** Schematic representation of the axonal projections of ePNs, which transmit olfactory information from the antennal lobes to the MB and MBONs relaying that information to LH promoting dishabituation, reinstatement of the naïve response. Activation of α/β KCs promotes dishabituation via MBON18, which projects directly to LH. Activation of γ MBns promotes dishabituation via the GABAergic MBONs 8,9 which project to the Crepine neuropil. As the Crepine is connected to LH in the Drosophila connectome, this pathway may provide an indirect route for conveying dishabituation related signals to LH.

We further examined the role of representative MBONs that might relate dishabituation information from γ KCs, which synergize with their aβ counterparts to drive the process (Fig 4J, N). Driver MB110C marks two GABAergic MBONs, MBON-γ3 and ΜΒΟΝ-γ3β΄1 (or MBONs 8 and 9; Fig 10D) arborizing in the respective γ compartments and projecting to Crepine/Lateral Accessory Lobe area (Aso, Hattori et al. 2014). Photoactivation of these MBONs in OCT-habituated flies partially restored avoidance (Fig 10E), but did not affect the response to this odor in naïve animals (Fig 10F). These results suggest that MBONs 8 and 9 contribute to dishabituation, but not OCT avoidance *per se*, by relaying information from γ KCs to the LH.

Taken together these findings demonstrate that cholinergic MBONs downstream of the αβ and γ KCs, including MBONs 8, 9 and 18 are required to convey the output of the MBs during dishabituation to higher-order centers, including the Crepine and LH and are summarized in Fig 10G.

### Specificity of the circuit and neuronal activities driving dishabituation from OCT

Collectively, the results support a model where dishabituating stimuli converge onto the MBs, where with modulation from DANs and APLNs engage distinct MBONs mediate dishabituation. Surprisingly, photoactivation of most neuronal elements examined herein, apart from α2α΄2 and α3 PPL1 DANs, sufficed to restore avoidance in OCT-habituated flies. This raised the question of whether robust afferent neurotransmission from any CNS circuit, emulated by photoactivation might suffice to reverse habituation to OCT. Then dishabituation could simply reflect robust activation of CNS circuits, independent of the specific pathways involved, thereby mimicking a generalized sensitization state as proposed by the Dual-Process Theory (Groves and Thompson 1979, Thompson 2009).

To address this possibility, we focused on Ellipsoid Body neurons-EBns (Fig 11A), which do not appear to ramify onto and are not considered part of the broader MB circuitry. Conditional silencing of EBns did not affect OCT avoidance, habituation and dishabituation either with footshocks or YO (Fig 11B). Control heterozygotes also performed as expected (Suppl Fig 8D). Importantly, photoactivation of EBns did not drive dishabituation in OCT-habituated flies at any intensity (Fig 10C) and had no effect on the aversive osmotactic response of naïve animals (Fig 10D).

**Figure 11:**
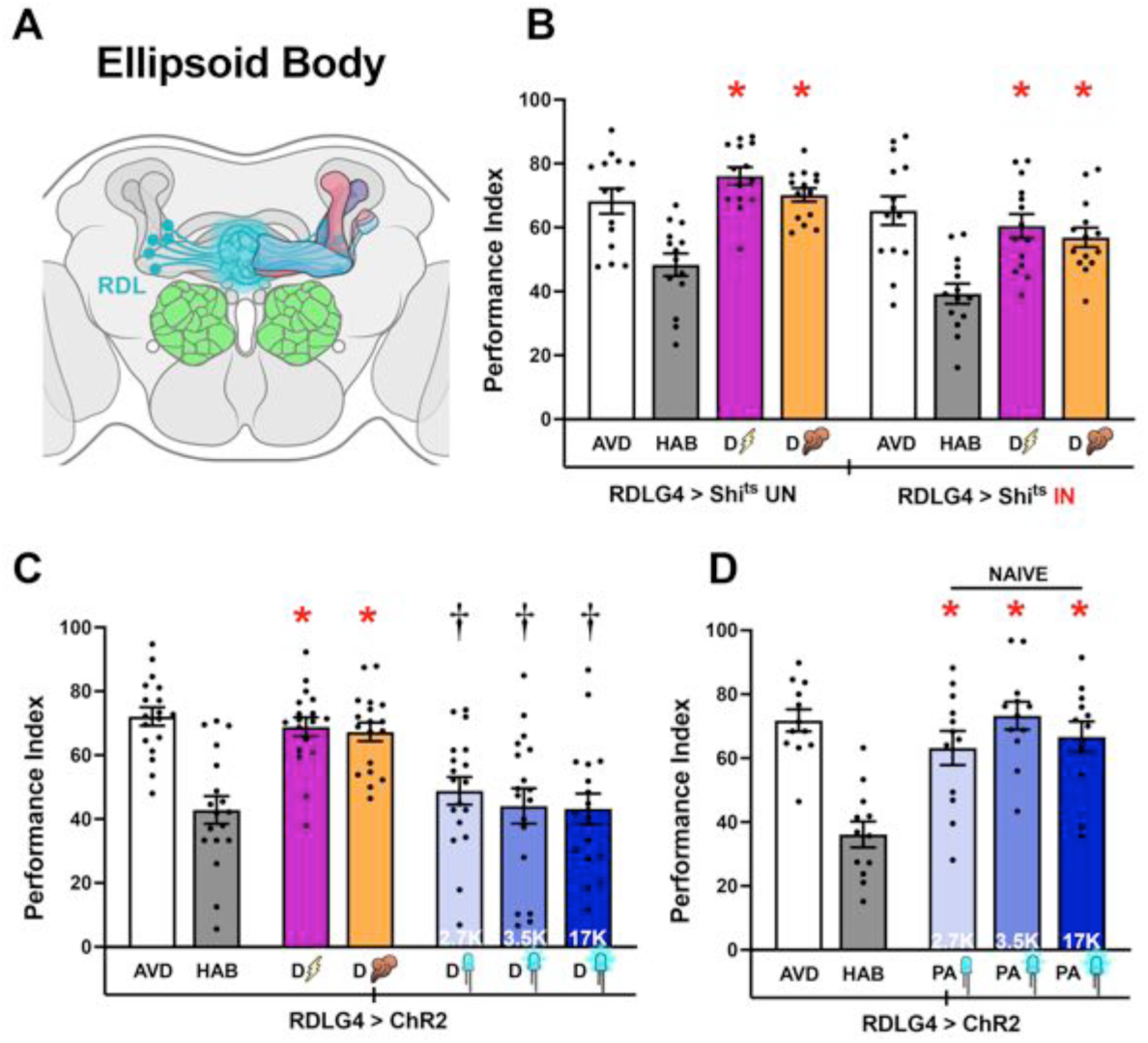
Ellipsoid Body Neurons do not participate in olfactory Habituation nor in Dishabituation. Mean Performance Indices ± SEM are shown in all figures. The naïve response for OCT is shown in white bars and is always significantly different from the habituated response (grey bars). Cross-modal (footshock) dishabituation is indicated by the bolt (magenta bar) and intramodal (YO), by the plume (orange bar). Exposure to blue light is indicated by the LED bulbs and increasing intensity (as indicated by the Klx) by deepening shades of blue. Stars in all Figures indicate significant differences (p<0.005), from the habituated response and daggers significant difference (p<0.005) from naïve avoidance. All statistical details are presented in Supplemental Table 1. **(A)** Schematic of Ellipsoid Body (EB) neurons. **(B)** Functional silencing of the EB neurons with UAS-shibire^ts^ under RDLGal4 compared to control animals carrying but not expressing the transgene (UN). Olfactory habituation and dishabituation with a 45V electric shock (magenta) and YO (orange), remained unaffected in both conditions. Additional controls run in parallel are presented in Supplemental Table 1. n=14. **(C)** Photoactivation of EB neurons under RDLGal4 in OCT-habituated flies with 3 light intensities did not restore avoidance, while a single footshock and YO exposure did. n=18. **(D)** Photoactivation of EB neurons in naïve flies with 3 light intensities did not affect OCT avoidance. n=12.

These results confirm our model, demonstrating that photoactivation of neurons that do not impinge on the MBs or the LH does not affect intramodal or cross-modal dishabituation in flies habituated to 4 min OCT exposure. Furthermore, emulating robust neuronal activity by photoactivation of any CNS circuit does not suffice to drive dishabituation through a generalized sensitization state.

## DISCUSSION

Here, we show in Drosophila that novel stimuli, whether presented within the same sensory modality as the habituating stimulus (intramodal) or through a different modality (cross-modal), reverse habituation through a MB-centered circuitry. Habituation in this olfactory paradigm manifests as attenuated responses to predictable, inconsequential 4-min OCT (Semelidou, Acevedo et al. 2018) or YO presentation (Fig 2G).

Addressing a key question on the role of sensitization in restoring habituated responses, we find no evidence that dishabituator-induced sensitization underlies intramodal dishabituation (Fig 2H, Suppl Fig 2C), nor that higher OCT intensities can dishabituate animals habituated to lower concentrations (Suppl Fig 1). Moreover, one or more 45V footshocks do not alter subsequent OCT avoidance (Acevedo, Froudarakis et al. 2007), indicating that cross-modal dishabituation is likewise unlikely the result of sensitization. Thus, neither broad CNS sensitization via footshock nor sensitization of the olfactory circuitry itself can account for dishabituation in this paradigm, in contrast to predictions of the Dual Process Theory (Groves and Thompson 1979, Thompson 2009).

Importantly however, we find that the habituating odorant and the dishabituator can in fact co-occur in the S-R circuitry. In this experiment, unlike all others reported here, we delivered the cross-modal footshock dishabituator at the end of the 4-min OCT habituating exposure but kept the OCT stimulus on. We measured OCT avoidance immediately after footshock and at 1-min intervals thereafter, up to 4 minutes (Fig 12). Although the habituator OCT was present, its avoidance emerged to naïve levels immediately (0) and the first minute post-dishabituator delivery and remained elevated, albeit partially, at 2 minutes. Therefore, the dishabituator’s neuronal activation pattern coexists with that established by the habituating stimulus and temporarily updates and overrides the reduced responsiveness to OCT.

**Figure 12:**
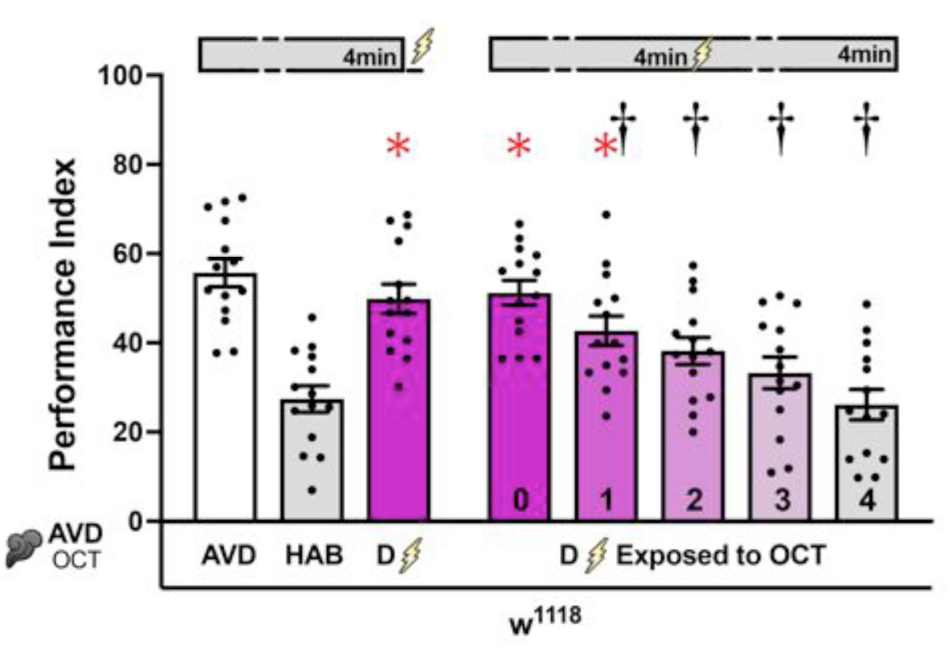
Continued exposure to Octanol during shock-induced dishabituation reinstates habituation. Mean Performance Indices ± SEM are shown in all figures. The naïve response for OCT is shown in white bar and is always significantly different from the habituated response (grey bar). Cross-modal (footshock) dishabituation is indicated by the bolt (magenta bar). Stars in all Figures indicate significant differences (p<0.005), from the habituated response and daggers significant difference (p<0.005) from naïve avoidance. All statistical details are presented in Supplemental Table 1. A) Naïve (AVD) and habituated (HAB) response to OCT of *w^1118^* flies, dishabituated by a single 45V electric shock. Exposure to OCT was overlapped with the dishabituator (footshock) and was maintained continuously for at least 4 minutes during which flies re-habituate normally, as indicated by the progressive transition from magenta to lighter grey bars. n=14.

Collectively our results demonstrate that habituation and dishabituation engage distinct yet interacting neuronal assemblies along the S-R circuitry, with differences in neurons conveying appetitive and aversive intramodal and cross-modal dishabituators. The neuronal elements engaged constitute distinct interacting nodes, which together mediate and regulate the level of OCT (or YO) avoidance depending on prior experience and context. These nodes and their functional contributions are discussed below.

### The MB node is central to both habituation and dishabituation

Odorants such as OCT and YO constituents must engage the canonical olfactory processing pathway with non-overlapping olfactory sensory neurons (OSNs) relaying information to the antennal lobe (AL) and subsequently to MBs and the LH. Odor identity is represented in the MBs by sparse activation of particular intrinsic neurons (sparse coding; (Honegger, Campbell et al. 2011)), which persists post odor exposure (Davis, 2011) whereas the LH is a major center for innate aversion and attraction odor-guided behavior (Jefferis, Potter et al. 2007, Dolan, Frechter et al. 2019, Das Chakraborty, Chang et al. 2022). Our findings place the MBs at the center of both habituation and dishabituation, where distinct KC subsets and their downstream output neurons regulate experience-dependent behavioral responses Habituation to 4-min OCT recruits inhibitory AL projection neurons (iPNs) that project to the LH, reducing naïve olfactory responses (Semelidou, Acevedo et al. 2018). Moreover, silencing the MBs and particularly the α΄β΄ KCs, results in premature habituation to OCT (Semelidou, Acevedo et al. 2018). Consistently, we demonstrate that optogenetic activation of α΄β΄ KCs suppresses OCT avoidance in naïve animals (Fig 4 H, Suppl Fig 5H). Likewise, silencing the cholinergic MBONs 16 and 17 (MBON-α΄1 and MBON-α΄3ap respectively), which arborize from α΄ KCs impairs habituation (Fig 9F), while their photoactivation in naïve animals attenuated OCT avoidance (Fig 9H). Together, these observations support a model in which α′β′ KCs and their downstream MBONs cooperate with iPN-mediated LH inhibition to establish the habituated behavioral state. The requirement for particularly strong α′β′ KC photoactivation to suppress naïve avoidance further suggests that this pathway normally acts synergistically with iPNs. This interpretation is in accord with the involvement of α΄β΄KCs in habituation to recurrent footshoks (Roussou, Papanikolopoulou et al. 2019, Foka, Georganta et al. 2022), suggesting that the same or similar circuit mediates habituation to multiple stimuli.

Dishabituation also engages the MBs but recruits distinct KC populations and downstream circuitry. The sudden appearance of a novel odor is expected to engage a different population of KCs from the habituating odor, apparently superimposed on that of the habituated odor as suggested by the data on Fig 12. Footshock likely reaches the MBs through ascending mechanosensory and nociceptive pathways and dopaminergic inputs (Yu, Akalal et al. 2006;Cervantes-Sandoval, Phan et al. 2017). Regardless of dishabituation stimulus modality, αβ and γ KCs emerge as the principal MB populations mediating both intramodal and cross-modal dishabituation (Fig 4N). Downstream, our results implicate MBONs 8, 9 and 18 as candidate output pathways relaying dishabituating signals toward higher-order centers. Information from the MBs is related to the LH, updating and adjusting the level of aversion or attraction in an experience-dependent manner (Benton 2022). Moreover, the inhibitory MBONs 8 and 9 innervate the Crepine area and may further modulate the activity of LH output neurons which arborize in this area (Dolan, Frechter et al. 2019).

Therefore, the collective results strengthen the previous model for habituation (Fig 1E) and extend it by positing the MB as a central hub where habituating and dishabituating signals converge. Rather than representing opposing processes within a single circuit, habituation and dishabituation appear to be mediated by partially distinct but interacting MB pathways, allowing novel stimuli to transiently override the behavioral consequences of prior experience while preserving the underlying sensory representation (Fig 12).

### The dopaminergic-APL node modulates the activity of the MB node

The interaction between KCs and MBONs is modulated by dopamine which modulates their synaptic output based on prior experiences (Aso, Hattori et al. 2014, Owald and Waddell 2015, Aso and Rubin 2016). Our results demonstrate that dopamine inputs are crucial for dishabituation, providing valence nuances and informing on the novelty of the dishabituating stimuli. In support of this view, collective activation of dopaminergic neurons (DANs) restored OCT avoidance in habituated animals (Fig 5E) but did not alter avoidance in naïve flies, Thie indicates that dopaminergic activity modulates specifically MB activity and not that of the LH driving innate responses (Jefferis, Potter et al. 2007, Dolan, Frechter et al. 2019, Das Chakraborty, Chang et al. 2022) .

DAN activity upon exposure to a dishabituator probably facilitates the recruitment of αβ and γ KCs, biasing MB afferent signaling towards dishabituation. In support of this view, collective activation of dopaminergic neurons (DANs) restored OCT avoidance in habituated animals (Fig 5E) but did not alter avoidance in naïve flies. Dopaminergic neurotransmission has also been implicated in gustatory dishabituation in Drosophila (Trisal, VijayRaghavan et al. 2023), but whether it engages the MBs and how in this paradigm remained unresolved.

Importantly, the results show that although dopamine mediates both intramodal and cross-modal dishabituation, it does so through the recruitment of distinct, valence-specific dopaminergic subsets. PAM-DANs are specifically required for intramodal dishabituation with the attractive YO (Fig 5G), while PPL1-DANs mediate dishabituation with the aversive cross-modal footshocks (Fig 5J) and importantly also with the aversive intramodal dishabituator OCT to YO habituated flies (Fig 6A). This segregation is congruent with the role of PAM-DANs in appetitive associative learning and memory (Burke, Huetteroth et al. 2012, Liu, Plaçais et al. 2012, Yamagata, Ichinose et al. 2015) and PPL1-DANs in aversive reinforcement of these processes (Claridge-Chang, Roorda et al. 2009, Aso, Siwanowicz et al. 2010, Perisse, Owald et al. 2016).

Thus, rather than simply conveying valence information (Claridge-Chang, Roorda et al. 2009, Aso, Hattori et al. 2014, Aso and Rubin 2016, Boto, Stahl et al. 2019), our results suggest that these dopaminergic pathways act instructively on activation of particular KCs/MBONs. This provides information about the behavioral significance of novel stimuli, allowing different dishabituators to access a common MB node through distinct dopaminergic subsets. This instructive activation apparently favors the activities of γ KCs and their afferent MBONs (Aso and Rubin 2016), such as the GABAergic γ3 (MBON8) and γ3β΄1 (MBON9) (Fig 10A, D). This view is supported by the differential engagement of γ-projecting PPL1-DANs in cross and intra-modal aversive dishabituators (Fig 5J, Fig 6A). Although PPL1 photoactivation does not result in full avoidance recovery (Fig 6B), they are clearly involved in naïve avoidance of intramodal stimuli.

How these DAN populations become selectively engaged remains uncertain. However we suggest that odors activate the MBs and the LH, with the latter relaying the signals to DANs (Otto, Pleijzier et al. 2020). It is likely then that aversive odors preferentially activate PPL1 and attractive ones their PAM counterparts, as a consequence of selective connectivity with aversion or attraction mediating LHONs (Cervantes-Sandoval, Phan et al. 2017, Otto, Pleijzier et al. 2020). Similarly, footshock avoidance mediated by the LH likely activates selectively PPL1 neurons which in turn modulate MB activity via their reciprocal synapses, which are known to engage largely γ KCs (PPL1-γ1pedc) forming a feedback loop (Cervantes-Sandoval, Phan et al. 2017, Li, Lindsey et al. 2020). Interestingly, PAM DANs also form connections with γ KCs, at least in the larva (Lyutova, Selcho et al. 2019), and may also do so in the adult by the connectome (Lin, Yang et al. 2024).

The differential effects of PAM and PPL1 activation further suggest that dopamine does not uniformly excite the MB circuit, but instead biases activity toward distinct KC populations and their downstream MBONs. Apart from the MB compartment-specific PAM and PPL1-DANs innervation (Aso, Grübel et al. 2009, Aso, Hattori et al. 2014, Aso and Rubin 2016), the MBs express both excitatory and inhibitory dopamine receptors in partially overlapping KC-type-specific patterns with distinct temporally specific roles, at least for associative learning (Aso, Ray et al. 2019; Handler, Graham et al. 2019). These receptors also exhibit different affinities for dopamine (Karam, Jones et al. 2020). This evidence suggests dynamic balance of the excitatory/inhibitory state of the different KC types upon dopamine release from PAM and PPL1 neurons. Such a mechanism may explain the different photostimulation thresholds required for PAM and PPL1-mediated dishabituation (Fig 5H, K).

We propose that dopaminergic modulation of KC activity is honed and modulated by the inhibitory activity of APLNs. The GABAergic APLNs arborize preferentially onto the α΄β΄ KCs (Suppl Fig 7H) but also form multiple reciprocal connections with all KC types (Liu and Davis 2009, Lin, Bygrave et al. 2014). Unlike PAM-DANs, APLNs regulate intramodal dishabituation irrespective of the valence of the dishabituator (Fig 8B, E). APLN inputs may also contribute to differentiation of intramodal habituating and dishabituating stimuli. Interestingly, even though APLs also respond to footshocks (Liu and Davis 2009, Zhou, Chen et al. 2019), they are not necessary for cross-modal dishabituation. These results suggest that APLNs participate in distinguishing the habituating odor from a novel odor rather than encoding stimulus valence. This notion is supported by data showing that APLNs respond with exposure length-dependent increased localized inhibition back to the KCs (Amin, Apostolopoulou et al. 2020), sparsening further the odor-specific KC pattern and limiting the impact of dopaminergic inputs on the MB (Prisco, Deimel et al. 2021).

Finally, our data suggest that MBON11 contributes to this regulatory node. Habituation requires, at least in part, release of the α΄β΄ inhibition and our data indicates that this is achieved by activation of the GABAergic MBON11 (Fig 9A), which projects broadly on the APL and appears to have the broadest KC input convergence (Li, Lindsey et al. 2020). We suggest that continued OCT exposure activates this odor excitable MBON, possibly by tonic dopamine release by the aversive odor-activated PPL1-γ1pedc (Perisse, Owald et al. 2016). PPL1-γ1pedc activation attenuates APL-mediated MB inhibition, which affects primarily the more heavily innervated α΄β΄ neurons. This favors neurotransmission from α΄β΄ neurons, leading to habituation. This view is strongly supported by abrogation of habituation upon silencing MBON11 and suppression of OCT avoidance upon its robust photoactivation in naïve animals (Fig 9B, D), a pattern also mirrored by the response of α΄β΄ KCs to these manipulations (Fig 4F, H).

Collectively, the results support a model in which DANs, APLNs and MBON11 form a critical node that instructively modulates and is modulated by the MBs. Distinct DAN populations convey the valence of novel stimuli, whereas APL-mediated inhibition shapes odor-specific KC representations and, together with MBON11, gates habituation upon odor exposure. Together, these modulatory signals bias the balance of KC output toward either habituation or dishabituation, allowing behavioral responses to be rapidly updated in response to changes in incoming stimuli

### The Lateral Horn constitutes the penultimate decision node of the habituation/dishabituation circuit

Although we did not directly interrogate the LH functionally, its critical role in mediating innate responses to olfactory and other stimuli, is well established (Schultzhaus, Saleem et al. 2017, Das Chakraborty and Sachse 2021, Das Chakraborty, Chang et al. 2022). We have shown that iPN inputs to the LH mediate olfactory habituation, suggesting a role for this region in experience-driven behavioral responses (Semelidou, Acevedo et al. 2018). In addition, functional interactions with the MB were shown to modulate bidirectionally experience-dependent behavioral outputs (Dolan, Belliart-Guérin et al. 2018), corroborating our findings. Here we show that MBONs impinging on the LH contribute both to habituation (Fig 9) and dishabituation (Fig 10), providing additional functional support for the role of LH in experience-driven responses.

Based on the behavioral output, we propose that dishabituation also occurs at the LH, resulting from engagement of neurons distinct from those whose activity is attenuated by iPN inhibitory inputs driving habituation (Semelidou, Acevedo et al. 2018). This hypothesis is supported by investigating the predicted synaptic outputs of the habituation-promoting MBON 16, 17 and the dishabituation-promoting MBON 18 in the LH. Interrogating the connectome revealed reciprocal inhibition between these MBON groups, which although limited, we posit it reflects the broader logic in the functional organization of the LH (Suppl Fig 9).

MBONs provide one instructive input route to the LH from the MBs the other being the iPNs from the AL. The habituation-promoting MBON 17 synapses on the GABAergic LHCENT8, which projects to the dishabituation linked MBON 18 and one of its targets, LHPV5e1. Furthermore, the cholinergic MBON16 and 17 share synaptic connections on the GABAergic mALB1 which project to LHPV10d1, another target of MBON 18. Our data support this hypothesis, with MBON16 and 17 photoactivation attenuating OCT avoidance in naïve flies (Fig 9H), probably bypassing iPN inputs due to theartificially string activation.

Conversely, the dishabituation-promoting cholinergic MBON18 synapses on the GABAergic LHCENT6, which projects to MBON16 (Suppl Fig 9). This suggests an inhibitory loop, which conditionally, in response to the dishabituator, inhibits the avoidance attenuating activity of MBON16. Hence, activation of MBON18 contributes to dishabituation by inhibiting MBON16 activity. In fact, activation of MBON18 in naïve flies results in strong OCT avoidance (Fig 10C).

Taken together, the functional and connectome evidence are consistent with the notion that the LH constitutes the penultimate decision node of the habituation/dishabituation circuit, which incorporates instructive inputs from the MBs via MBONs and the iPNs. The contribution of MBONs, even with current representative sample of these neurons, is evident in communication MB activity to the LH. During habituation, the 4 min odor exposure engages cholinergic MBONs that innervate inhibitory LH neurons. This instructive input from the MBs along convergent iPN activity result in inhibitory potentiation that suppresses avoidance mediating circuits yielding habituation. Conversely, novel dishabituating stimuli recruit instructive excitatory input to the LH, which engages distinct inhibitory LHNs, suppressing habituation-activity of the avoidance suppressing circuit. This model (Suppl Fig 10 and Fig 13B), along with the temporary efficacy of the dishabituator (Fig 12), is consistent with the previously published notion that dishabituation overrides, rather than eliminates habituation (Trisal, Aranha et al. 2022).

**Figure 13.**
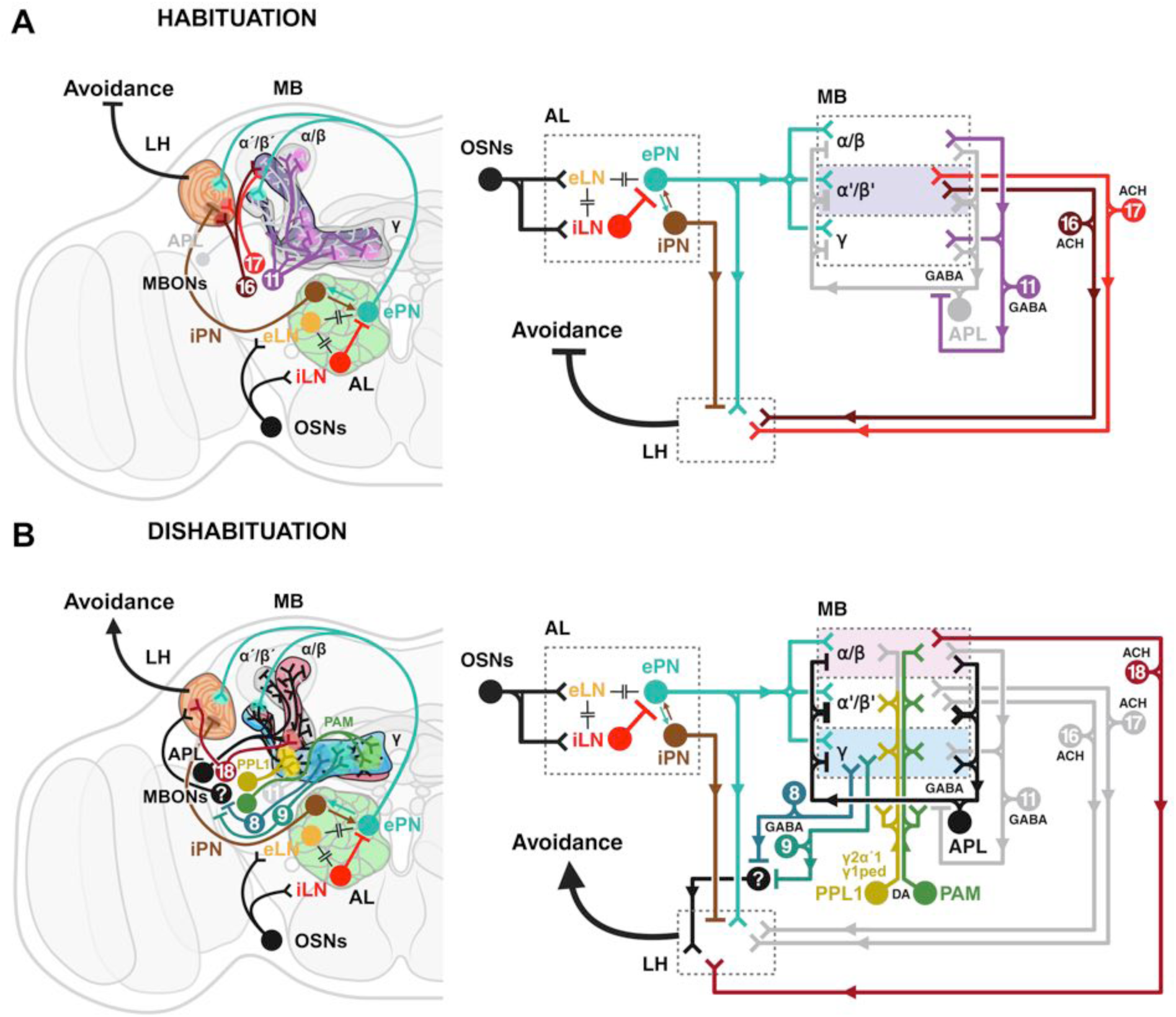
A working model of the neuronal circuits underlying (A) Habituation and (B) Dishabituation. The principle neuronal components mediating the behavior comprise the antennal lobe (AL-light green), the lateral horn (LH-orange) and the mushroom body (MB), divided in three lobes (α/β-pink, α΄/β΄-purple, γ-blue). In the stick figure the same neuronal compartments are presented as boxes oriented by a dashed line. Model created with Biorender.com Distinct neuronal subsets are marked with different colors; Olfactory sensory neurons (OSNs-black), excitatory local interneurons (eLN-light orange), inhibitory local Interneurons (iLN-red), excitatory projection neurons (ePN-dark cyan), inhibitory local Interneurons (iLN-brown), mushroom body output neurons (MBON8-blue-teal, MBON9-green-teal, MBON11-violet, MBON16-maroon, MBON17-scarlet red, MBON18-burgundy red), anterior paired lateral neuron (APL-black), PAM dopaminergic neurons (PAM-dark green), PPL1 dopaminergic neuronal subsets (PPL1-γ2α’1 & PPL1-γ1pedc both in mustard yellow). Neurons not involved in the progression from habituation to dishabituation (reinstatement of avoidance levels), are colored light grey for clarity. Synapses thought to be electrical connecting interneurons of the AL such as the eLN, iLN & ePN are indicated by -||-. Synapses show the dendritic / axonal projections of neurons and arrows display the information path in the circuit. Blunted axonal projections indicate inhibition.

### Habituation and dishabituation from 4-min OCT exposure requires dynamic internodal interactions

Habituation and dishabituation to 4-min OCT exposure constitute experience-dependent adaptive changes to the innate aversive osmotaxis mediated by the LH (Schultzhaus, Saleem et al. 2017, Das Chakraborty and Sachse 2021). Habituation and dishabituation emerge from dynamic interactions between three major circuit nodes: the MBs, the dopaminergic/APL modulatory network, and the LH, where innate olfactory behavior is ultimately expressed (Schultzhaus, Saleem et al. 2017, Das Chakraborty and Sachse 2021). We propose that these nodes do not function independently, but continuously interact to determine whether avoidance is maintained, suppressed, or reinstated.

During habituation (Fig 13A) olfactory information reaches both the LH and the MB through excitatory projection neurons. Previous work demonstrated that delayed recruitment of inhibitory projection neurons suppresses LH circuits mediating innate avoidance and is required for habituation (Semelidou, Acevedo et al. 2018). In this view, the default state of the circuit favors continued avoidance, but sustained OCT or YO sensory input gradually biases network activity toward the habituated state. Our results indicate that this process is complemented by activity within the MBs. Specifically, α′β′ KCs together with their downstream MBONs promote habituation, while APL-mediated inhibition shapes odor-specific KC activity and MBON11 likely regulates this inhibition through feedback onto the APL. MBON11-mediated disinhibition of α΄β΄ KC neurotransmission in then relayed to the LH though MBONs16 and 17, and acts in synergy with delayed inputs from iPNs that suppress the innate OCT avoidance and establish habituation.

We posit therefore that the default state of the MB and DAN/APL nodes favor continued avoidance of the aversive odor stimulus for the time we have defined as latency to habituate, mediated by excitatory ePN input on the LH and MBs (Semelidou, Acevedo et al. 2018). To attenuate this default avoidance behavior requires instructive signals from both nodes to the LH, along with inputs from the AL/iPN node. Persistent activity of KCs post odor exposure has been reported under different contexts (Yu, Ponomarev et al. 2004, Davis 2011). Therefore, the OCT pattern of KC activation persists even after odor exposure has ceased. This period is likely reflected in the 4-6 minutes required for spontaneous recovery (Rankin, Abrams et al. 2009) of OCT habituation (Semelidou, Acevedo et al. 2018).

The sudden presentation of a novel odor, or footshock initiates a fundamentally different sequence of events. This information is relayed to the MBs and generates its unique activation signature therein (Honegger, Campbell et al. 2011). We show that dopaminergic subsets mediate differentiation of dishabituator valence, most likely based on inputs from the LH (Otto, Pleijzier et al. 2020). Importantly, in a manner analogous to their role in signaling unexpected outcomes during olfactory learning (Zhao, Widmer et al. 2021), we posit that DAN activation detects the error between the predicted OCT stimulus and the actual dishabituator input, even if they are contemporary (Fig 12). Together with APL-mediated inhibition of α΄β΄ neutotransmission necessary for habituation, these modulatory neurons promote dishabituation. This incongruence temporarily enhances APL activity, which preferentially attenuates α΄β΄ neutotransmission, which is necessary for habituation.

Instructive signals onto the MBs via PPL1-γ1pedc generate conditions favoring neurotransmission from γ and αβ KCs, which is relayed to the LH of both hemispheres via the cholinergic MBON18 and GABAergic inputs to the Crepine by MBONs 8 and 9 (Fig 10 A, D). These inputs to the LH and convergent iPN inputs result in temporary attenuation of the habituation promoting MBON16 activity, most likely among other such neurons (Suppl Fig 9), resulting in suppression of the habituated response and emergence of increased avoidance manifested as dishabituation.

In the proposed model (Fig 13), the MBs are the node where the expected olfactory stimulus is compared to an unexpected dishabituator through dopaminergic and GABAergic modulation. Their output to MBONs is modulated by DANs and the APL. This tripartite interaction is ostensibly reflected in the similar effects on habituated flies of artificially emulating this activity by photoactivation. However, the MBs do not directly generate the behavioral transition from habituation to dishabituation. The resulting shift in MB output is transmitted through distinct MBON pathways to the LH, where competing avoidance-suppressing and avoidance-promoting circuits determine the behavioral outcome. In this model, dishabituation does not erase the habituated state but transiently overrides its behavioral expression, consistent with the temporary nature of dishabituation observed in our behavioral experiments.

Although the detailed LH circuitry remains to be experimentally validated, the proposed multi-nodal model is in accord with work in vertebrates suggesting distinct circuitries are driving habituation and dishabituation and propose that the hippocampus processes and mediates responses to novel stimuli, differentiating predicted habituating stimuli from the unexpected ones driving dishabituation (Thinus-Blanc, Save et al. 1991, Wang and Arbib 1992). Understanding these interactions in the genetically tractable Drosophila brain may therefore provide significant insights into the neural mechanisms of attention and cognition in other species, including humans (Kavsek and Bornstein 2010).

Finally, we posit that understanding habituation and dishabituation and its underlying mechanisms in model systems such as Drosophila is likely to have significant implications in studies of attention and cognitive processing in other species including humans (Kavsek and Bornstein 2010). Processes underlying habituation and dishabituation, such as those described herein, offer models that reveal how the brain filters updates and prioritizes information and how distinct, interacting neuronal circuits underlie these processes. This has broader significance for understanding memory storage and updating in general, as well as translational relevance to conditions involving sensory gating deficits, such as schizophrenia and autism.

## MATERIALS AND METHODS

### Drosophila strains

*Drosophila melanogaster* strains were cultured in standard wheat-flour-sugar food supplemented with soy flour and CaCl_2_ at 25°C, unless specified otherwise. For efficient RNA interference (RNAi)-mediated abrogation with the temperature-sensitive TARGET system, flies were raised at 18°C until hatching, to keep the transgenes silent and then placed at 30°C for 2 days prior to testing to induce the transgenes. Control *w^1118^* flies had been backcrossed to the Canton-S wild type strain for at least 10 generations (*w^1118^* strain). All other strains had been backcrossed to the Cantonised-*w^1118^* for four to six generations prior to use in behavioral experiments.

The UAS-Shibire^ts1^ transgene (BDRC #44222) bearing a temperature-sensitive mutation in Dynamin attenuating re-uptake of synaptic vesicles (Kitamoto 2001) was kindly provided by RL Davis. For the optogenetic activation experiments the robustly expressed UAS-ChR2-XXL transgene encoding for the D156C mutant of Channelrhodopsin-2 (BDRC #58374) was used, courtesy of André Fiala. ChR2-XXL presents an extended open state of the channel and can be efficiently photoactivated with without requiring retinal supplementation (Dawydow, Gueta et al. 2014).

The Dopaminergic system was targeted with the pleGal4 driver (Aso, Hattori et al. 2014) (BDRC #68302). The PAM subset of dopaminergic neurons was targeted with the 0273Gal4 driver (Burke, Huetteroth et al. 2012), a kind gift of S. Waddell (Oxford University, Oxford, UK). Conversely, the PPL1 subset of DANs was targeted with MB504BGal4 (Vogt, Schnaitmann et al. 2014) (BDRC #68329), obtained from R. Tanimoto (Tohoku University, Sendai, Japan). Drivers for subsets of the PPL1-DANS, PPL1-γ1pedc, PPL1-γ2α΄1, PPL1-α2α΄2, PPL1-α3 (Aso, Hattori et al. 2014, Aso and Rubin 2016), were obtained from BDRC (#68253, #68308, #68278, #68334 respectively), as was the MB110C driver (#68262), targeting the MBONs 8, 9 afferent from γ3 and γ3β΄1 compartments(Aso, Hattori et al. 2014). The MB543B driver (BDRC #68335), targeting MBONs 15,16,17 afferent from α′1, α′3ap and α′3m compartments respectively, driver MB027B (BDRC #68301) targeting the MBONs 16 and 17, driver MB050B (BDRC #68365) targeting the MBONs 15 and 18 and finally, driver MB112C (BDRC #6MBON 11, were also obtained from BDRC (Aso, Hattori et al. 2014).

The dncGal4 driver (BDRC #48571) was used to target all KCs (Foka, Georganta et al. 2022). The αβ KCs were targeted with two drivers: c739Gal4 (Pavlopoulos, Anezaki et al. 2008, Georganta, Moressis et al. 2021) (BDRC #7362) and 17dGal4 (Akalal, Wilson et al. 2006), provided by RL Davis (Scripps Institute, University of Florida, USA). Drivers targeting the α΄β΄ KCs, were the c305aGal4 (Krashes, Keene et al. 2007) and VT030604Gal4 (Roussou, Papanikolopoulou et al. 2019) (VDRC #200228), kind gifts from S. Waddell (Oxford University, Oxford, UK). To mark the γ KCs, two drivers were used: MB131BGal4 (Aso, Hattori et al. 2014) (BDRC #68265), kindly provided by Y. Aso (Janelia Research Campus, Howard Hughes Medical Institute, Ashburn, VA) and VT044966Gal4 (VDRC #203571), a gift of K. Keleman (Howard Hughes Medical Institute).

The APLGal4 driver has been described previously (Wu, Shih et al. 2013) and the transgenic lines encoding interfering RNA (RNAi lines) were obtained from VDRC: UASGad RNAi (VDRC #32344), UASTbh RNAi B (VDRC #51667) and BDRC: UASTbh RNAi A (BDRC #27667) respectively. The previously described GadGal4 (BDRC #51630) (Buchner 1991, Georganta, Moressis et al. 2021) and RdlGal4 (Jenett, Rubin et al. 2012) (BDRC #49620) were obtained from BDRC. The Olfactory Receptor Co-receptor (Orco) mutants were sourced from BDRC (Or83b^1^ #23129, Or83b^2^ #23130).

Finally, TubGal80^ts^ transgenes were obtained from RL Davis (Scripps Institute, University of Florida, USA) and were introduced to Gal4 driver strains by standard genetic crosses or recombination as indicated.

### Behavioral experiments

To obtain animals for behavioral experiments, Gal4 driver homozygotes were crossed to UAS-Shibire^ts^ and UASChR2-XXL *en-masse* to produce the experimental groups. Similarly, Gal4 driver homozygotes, UAS-Shibire^ts^ and UAS-ChR2-XXL were mass-crossed to *w^1118^* flies, to obtain isogenic heterozygous controls. For the RNAi experiments APLGal4;Gal80^ts^ homozygotes were crossed to the RNAi transgene-bearing lines and to *w^1118^*or *y^1^;v* accordingly to produce matching genetic backgound isogenic heterozygous controls. All flies used in behavioral experiments were tested 3–5 days after emergence. The flies were transferred to clean vials with fresh food one hour before conducting behavioral experiments. All experiments were performed under dim red light at 25°C with humidity maintained between 60% and 70%.

The responses to 3-Octanol (habituator) and the Shock and Yeast Odor dishabituators were set up as described before (Semelidou, Acevedo et al. 2019), and were re-optimized with Canton-S and *w^1118^* animals. Strains expressing Shibire^ts^ were systematically assessed for OCT avoidance under permissive and non-permissive conditions. Although variable among genotypes, OCT avoidance was not significantly different between permissive and non-permissive conditions for all strains tested (Suppl. Fig 7). Naïve Canton-S and *w^1118^* flies were subjected to optogenetic activation to ascertain that the treatment does not affect OCT avoidance *per se* (Suppl Fig 2B, D).

For neuronal silencing experiments, UAS-Shibire^ts^ -expressing strains were placed in a 32°C incubator for 30 min prior to the start of the “training phase” to inactivate the transgenic temperature-sensitive Shibire protein, which recovers its full activity within 15 min after removal from the restrictive temperature (Kitamoto 2001, McGuire, Le et al. 2001). The isogenic heterozygous controls were subjected to the same treatment, while the uninduced control group remained at the original experimental temperature of 25°C.

Flies expressing RNAi-mediating transgenes were transferred to vials with fresh food one hour prior the experiment and placed again at 30°C to sustain the silencing effect. The isogenic heterozygous controls were subjected to the same treatments as the experimental group, while the uninduced control group remained at 18°C after hatching for two days to avoid transgene expression, transferred to fresh food one hour before the experiment and remained at the experimental temperature of 25°C.

For optical modulation of neuronal activity, the flies were stimulated with a continuous 3-sec light pulse at 460nm, of three different light intensities generated by modulating the electric potential of the diode to 4.5V, 5V, and 6Vwhich yielded 2.7, 3.5 and 17Klx measured with a UNI-T model UT383S photometer. To emulate a dishabituator, flies expressing the UAS-ChR2-XXL transgene were photoactivated after the 4-minute OCT exposure inside the training arm. After a 30-second rest phase, they were then given the choice between air and OCT for 90 seconds. The different photoactivation intensities were meant to ascertain that neurons would not be unresponsive because of subthreshold stimulation. To test the role of the photoactivated neurons in mediating avoidance *per se,* naïve flies expressing the UAS-ChR2-XXL transgene were exposed to light without prior exposure to OCT and then tested for avoidance of the odor.

### Olfactory habituation and dishabituation

Olfactory habituation experiments were performed as described previously (Semelidou, Acevedo et al. 2018). For “habituation training”, approximately 60-65 flies were exposed in the upper arm of a standard T-maze to an airstream (500 ml/min) drawn over a 5.7cm^2^ meniscus of the aversive odor DL-3-Octanol (Thermo Scientific) for 4 minutes. Immediately after odor exposure the flies were treated with one of the dishabituators, a single 1.25 sec 45V DC electric shock, or 3 sec of air drawn over a meniscus of 1.8 cm^2^ of brewer’s yeast solution (0.3 gr brewer’s yeast/1ml dH_2_O) we name “yeast odor”, YO. Electric shock is an aversive mechanosensory stimulus and as previously described, the number of shocks does not have a significant effect on dishabituation (Semelidou, Acevedo et al. 2018). The 45V footshock used as a dishabituator (Fig 1B, C) is weaker than that used for associative conditioning (Georganta, Moressis et al. 2021), but was avoided robustly despite the variability among genetic backgrounds (Suppl Fig 7A, B) and weak stimuli can be at least as effective dishabituators as stronger ones (Marcus, Nolen et al. 1988). Hence, the footshock dishabituator (Fig 1B, C) represents a behaviorally salient novel stimulus and failure to drive dishabituation is not due to limited ability to sense and respond to it.

In contrast, extending the exposure to YO, shifts the response of control flies from initial attraction to an aversive response. This observation prompted implementation of the appetitive 3-second yeast puff as a dishabituator. YO was also used as the habituator and a puff of OCT as the dishabituator following the same paradigm, to assess whether dishabituation efficiency is driven by stimulus novelty rather than by the valence of the dishabituator relative to the habituated stimulus (Fig 6).

The aversive odor OCT was also used as a dishabituator in control files habituated to different dilutions of OCT (0.1, 1, 10, 20, 50,100%) to test for within modal dishabituation driven by changes in the intensity to the same stimulus (Suppl Fig 1).

After training and a 30-second rest, the animals were positioned in the choice point of the maze to assess their osmotactic response. They were presented with a choice between air and OCT for 90 seconds. Subsequently, the flies were confined to the arm they selected and tallied. The Performance Index (PI) was derived by subtracting the percentage of flies favoring the scented arm (OCT) from the percentage congregating in the unscented arm (air). Habituation to OCT is manifested as post-exposure reduced avoidance compared to that of naïve animals.

In the paradigm described above, habituator and dishabituator presentations are sequential. To assess if olfactory habituation is reinstated following dishabituation, exposure to OCT continued through dishabituator (footshock) delivery and was maintained continuously for at least 4 minutes, during which flies habituated efficiently (adapted from Groves & Thompson, 1970). More specifically, flies were exposed to 4 min of OCT, dishabituated with a footshock and following 0, 1, 2, 3 and 4 minutes of continuous OCT exposure were positioned in the choice point of the maze to assess their osmotactic response (Fig 12).

### Shock habituation and dishabituation

Habituation to footshocks was performed as described previously (Foka, Georganta et al. 2022). For exposure to the habituator, approximately 60 flies were exposed in the upper arm of a standard T-maze to 15 recurrent 45 V DC footshocks via an electrified copper grid for 90 seconds. Immediately after shock treatment, the flies were presented with the YO dishabituator described above, followed by positioning them on the center of the maze to assess their subsequent avoidance of an electrified maze arm, *versus* an arm carrying an inert grid. Following a 90 second choice period, the flies were trapped and tallied. The Performance Index (PI) was calculated by subtracting the percentage of flies favoring the inert grid from those congregating in the arm with the electrified grid.

### Odor pre-exposure on Odor Avoidance

To assess the effects of prior sensory exposure on odor avoidance, flies were pre-exposed to either YO or OCT, and their subsequent avoidance was measured to different OCT dilutions. Flies were pre-exposed to YO for the indicated durations of 5sec and 15 sec, where the valence of the stimulus is attractive and aversive respectively, followed by a stimulus-free interval of either 10, 30, or 90 sec. Then flies were tested for avoidance of different dilutions of OCT (0.1, 1, 10,5 0, 100%) and tallied (Fig 2H & Suppl Fig 2C).

### Statistical analyses

For all experiments, both control and genetically matched experimental groups were tested together within the same session, employing a balanced experimental design. The sequence of training and testing between the different groups was randomized to eliminate temporal bias. In cases where two or three genetic controls were utilized, significant differences from all the controls were required for results from experimental animals to be considered significantly different. For all experiments employing expression of Shi^ts^ only the experimental genotypes (UN *versus* IN) are shown in the relevant figures for simplicity. However, all control genotypes were run in parallel as indicated in the cumulative Statistics Table (Suppl Table 1) and were included in the statistical analysis of the experiment, estimating ANOVAs and p values.

The raw data were analyzed using parametric methods after ascertaining normality with the JMP statistical software package from SAS Institute. Bonferonni-corrected ANOVA tests were conducted, followed by planned comparisons (least square mean contrast analyses) to assess significant differences among genotypes. The significance level was adjusted for experiment-wise error rates, following the approach suggested by Sokal and Rohlf (Sokal and Rohlf 1981). Detailed statistical comparisons are provided in the text and summarized in the collective statistics table (Suppl Table 1).

### Artwork

All models and artwork were created with Biorender.com.

## Supporting information

all supplemental data

## Acknowledgements

The authors would like to thank Y. Aso, R. Davis, A. Fiala, R. Tanimoto, S. Waddell and the Bloomington Drosophila Stock Center for fly strains. M. Loizou for the endless supply of fly food and fly husbandry and all members of the lab for fruitful debates and discussions. This work was supported by a grant from the Hellenic Foundation for Research and Innovation (HFRI FM17-ΤΔΕ2961) to EMCS.

