## Supplementary material for "Multi-nodal regulation of olfactory dishabituation in Drosophila": all supplemental data

### SUPPLEMENTAL FIGURES AND LEGENDS

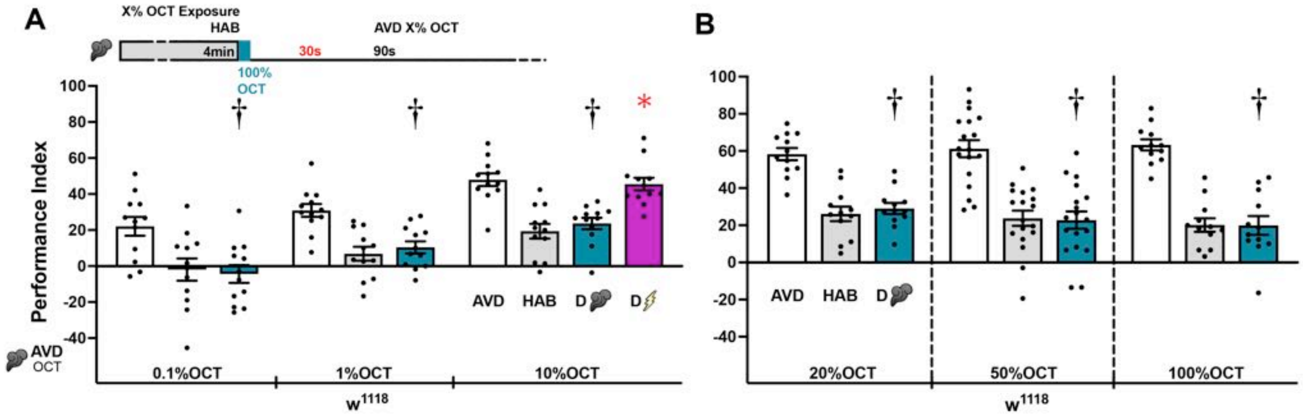

**Supplemental Figure 1: Flies habituated flies by a range of Octanol dilutions cannot be dishabituated by concentrated Octanol.**

**A-B)** Mean Performance Indices ± SEM are shown in all panels. The naïve avoidance response (AVD) is shown by the white bar for the different dilutions of OCT (AVD X% OCT). The habituated response (HAB) is shown by the grey bars for the different dilutions of OCT (HAB X% OCT). The dishabituated response is shown in magenta for the footshock dishabituator and in cyan for 100% OCT. Stars indicate significant differences (p<0.005) from the habituated response and daggers significant difference (p<0.005) from naïve avoidance. All statistical details are presented in Supplemental Table 1.

**A)** Naïve response of $w^{1118}$ flies to 0.1%, 1% & 10% OCT (AVD). The habituated (HAB) response to the stated OCT dilutions cannot be dishabituated with a puff of 100% OCT (plume), whereas a single 45 V electric shock (bolt) reinstates the naïve response. n=12 for all dilutions.

**B)** Naïve response of $w^{1118}$ flies to 20%, 50% & 100% OCT (AVD). The habituated (HAB) response to the stated OCT dilutions cannot be dishabituated with a puff of 100% OCT (plume). Experiments were conducted separately for each dilution. n=12 for 20%, n=18 for 50% and n=12 for 100% of OCT.

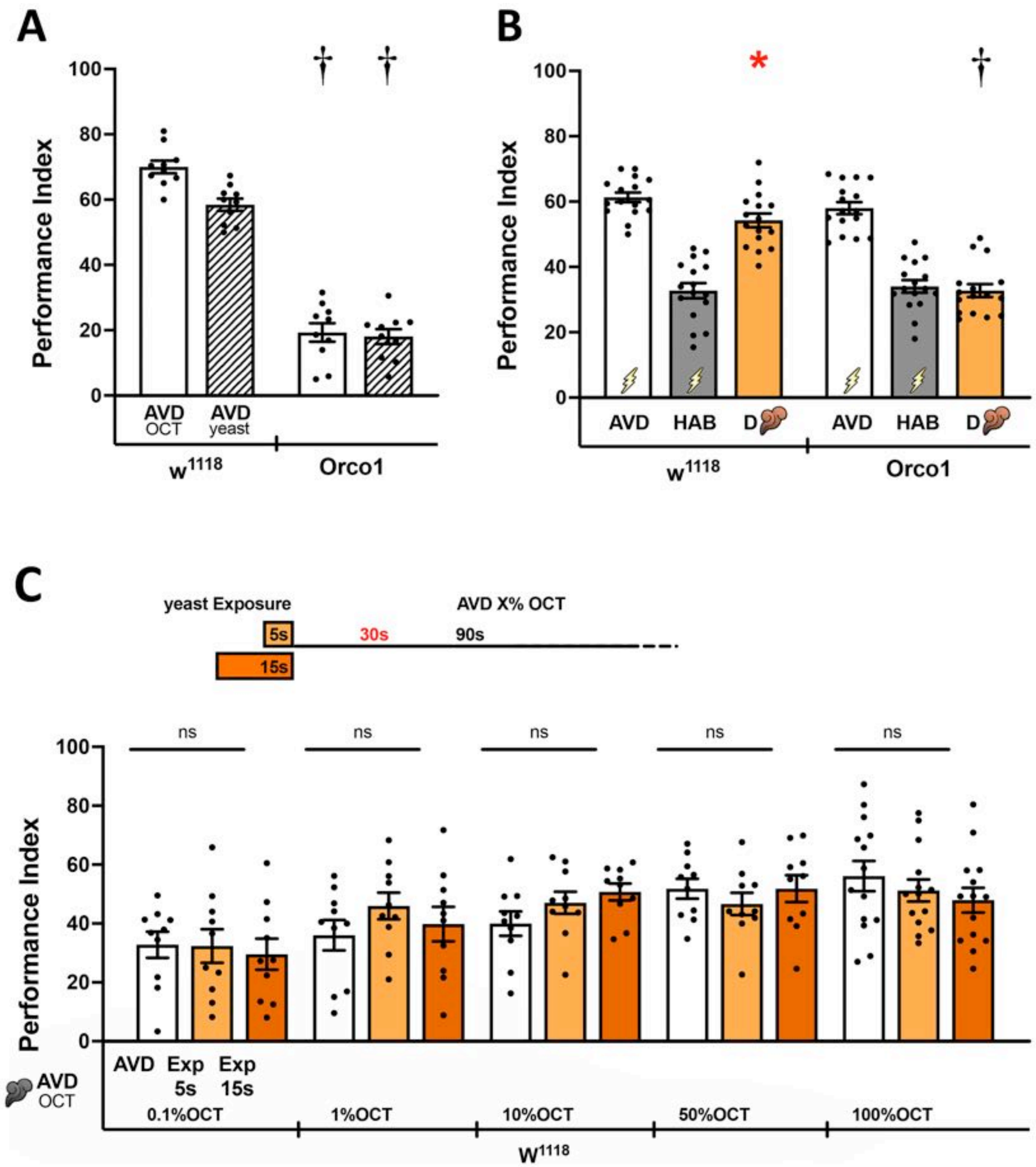

**Supplemental Figure 2: *Orco[1]* mutants do not respond to YO stimulation, which is not an does not drive dishabituation by sensitization.**

Mean Performance Indices ± SEM are shown. The naïve avoidance response (AVD) is shown by the white bar-OCT and white striped bar-yeast odor (YO). The habituated response (HAB) to footshock is shown by the grey bars and the dishabituated response by YO exposure (plume) in orange. Pre-exposures of 5sec and 15sec to YO are shown in orange and dark orange respectively. Stars indicate significant differences ($p<0.005$) from the habituated response and daggers significant difference ($p<0.005$) from naïve avoidance. All statistical details are presented in Supplemental Table 1.

**A)** Naïve response of $w^{1118}$ and $Orco^1$ homozygotes to OCT and YO. Daggers indicate significant differences between the respective naïve responses to OCT and YO of $Orco^1$ homozygotes and controls. n=10.

**B)** Footshock habituation cannot be dishabituated by YO presentation in $Orco^1$ homozygotes, unlike in control animals. n=16.

**C)** Pre-exposure of $w^{1118}$ flies to YO for the indicated durations, followed by a stimulus-free interval of 30 sec, does not alter subsequent avoidance of a range of OCT dilutions. n≥10

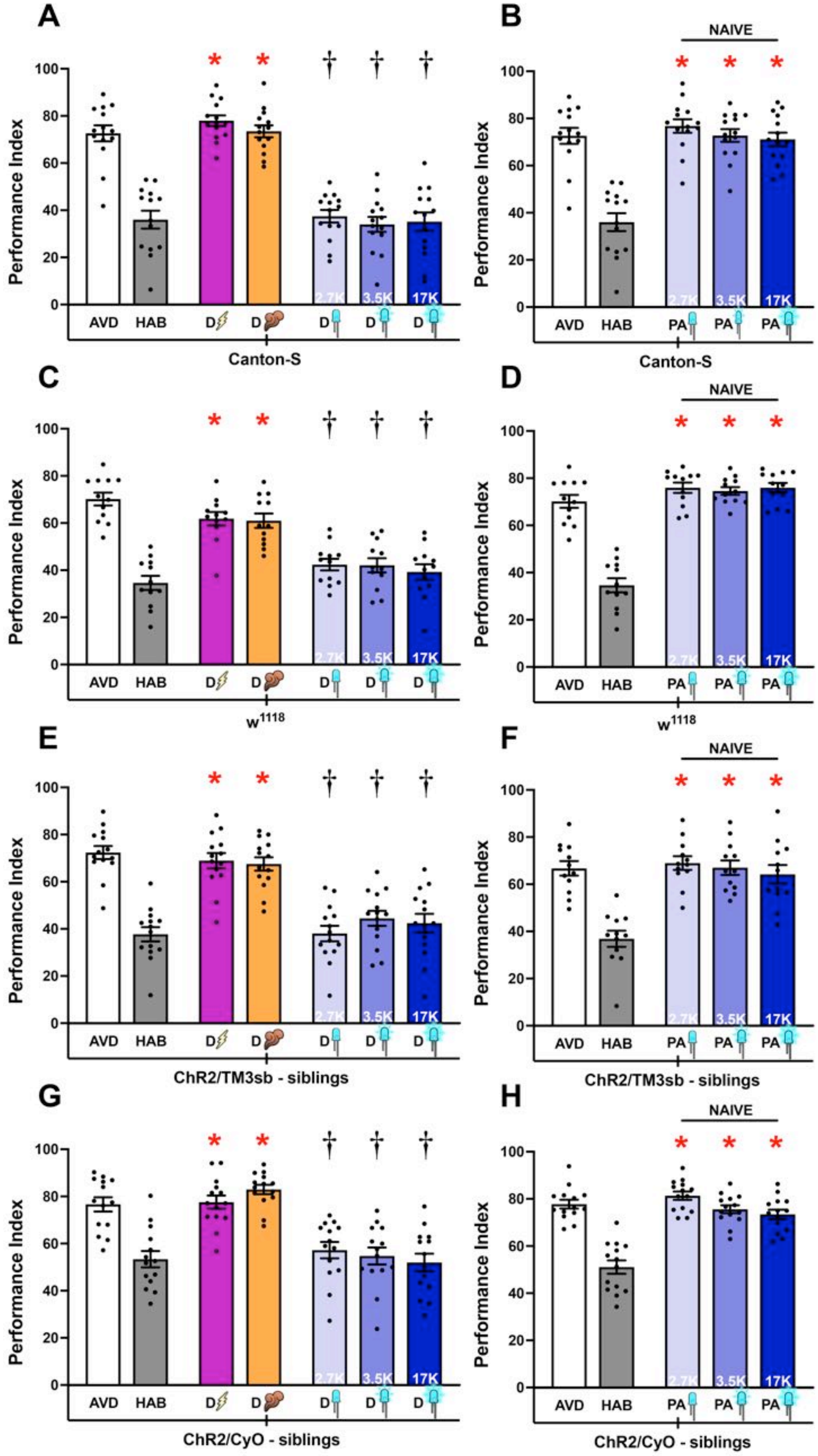

**Supplemental Figure 3: Photoactivation Controls.**

Mean Performance Indices ± SEM are shown in all figures. The naïve response for OCT is shown in white bars and is always significantly different from the habituated response (grey bars). Cross-modal (footshock) dishabituation is indicated by the bolt (magenta bar) and intramodal (YO), by the plume (orange bar). Exposure to blue light is indicated by the LED bulbs and increasing intensity (as indicated by the lx) by deepening shades of blue. Stars in all Figures indicate significant differences ($p<0.005$), from the habituated response and daggers significant difference ($p<0.005$) from naïve avoidance. All statistical details are presented in Supplemental Table 1.

**A)** Canton-S and **C)** $w^{1118}$ flies did not dishabituate upon photoactivation eliminating the possibility that the blue light was in fact the dishabituator. n=14 for A and n=12 for C.

**B)** Canton-S and **D)** $w^{1118}$ naïve flies upon photoactivation presented normal avoidance levels to OCT. n=14 for B and n=12 for D.

**E-H)** Similarly, the UAS-ChR2XXL/TM3Sb and UAS-ChR2XXL/CyO sibling flies of non-balanced animals expressing ChR2XXL also remained unresponsive to optogenetic activation (n=14 for E,G,H and n=12 for F).

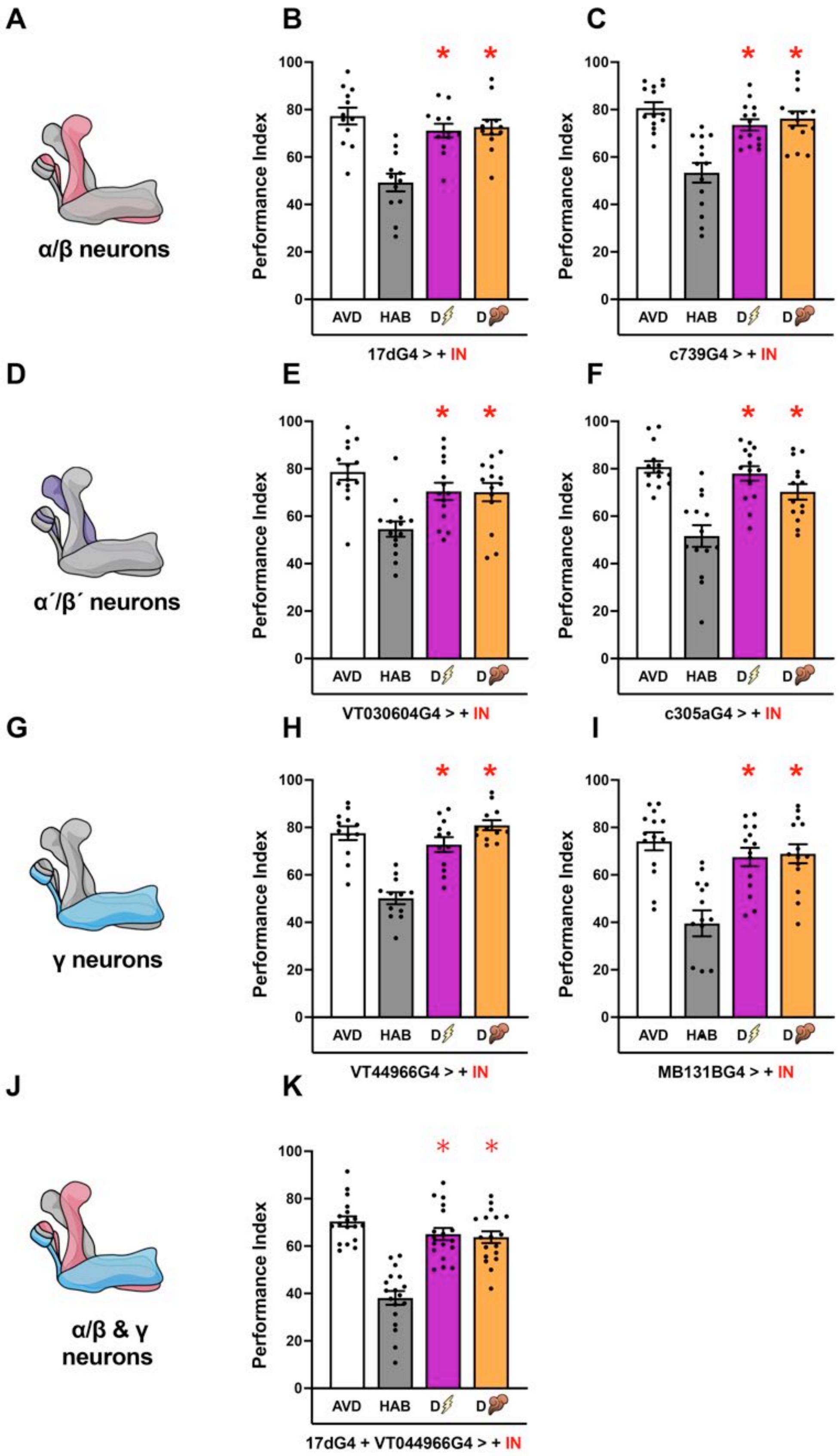
A
α/β neurons
B
Performance Index
AVD
HAB
D
17dG4 > + IN
C
c739G4 > + IN
D
α′/β′ neurons
E
VT030604G4 > + IN
F
c305aG4 > + IN
G
γ neurons
H
VT44966G4 > + IN
I
MB131BG4 > + IN
J
α/β & γ neurons
K
17dG4 + VT044966G4 > + IN

**Supplemental Figure 4: Sibling Controls for MB neuronal subsets.**

Mean Performance Indices ± SEM are shown in all figures. The naïve response for OCT is shown in white bars and is always significantly different from the habituated response (grey bars). Cross-modal (footshock) dishabituation is indicated by the bolt (magenta bar) and intramodal (YO), by the plume (orange bar). Stars indicate significant differences ($p<0.005$) from the habituated response. All statistical details are presented in Supplemental Table 1.

**A, D, G, J)** Schematic of the types of KCs (αβ, α΄β΄, γ and αβ & γ).

**B, C)** Controls for αβ KCs (n=12 for B and n=14 for C).

**E, F)** Controls for α΄β΄ KCs (n=14 for E and n=14 for F)

**H, I)** Controls for γ KCs (n=12 for H and n=14 for I).

**K)** Control for αβ & γ KCs (n=18).

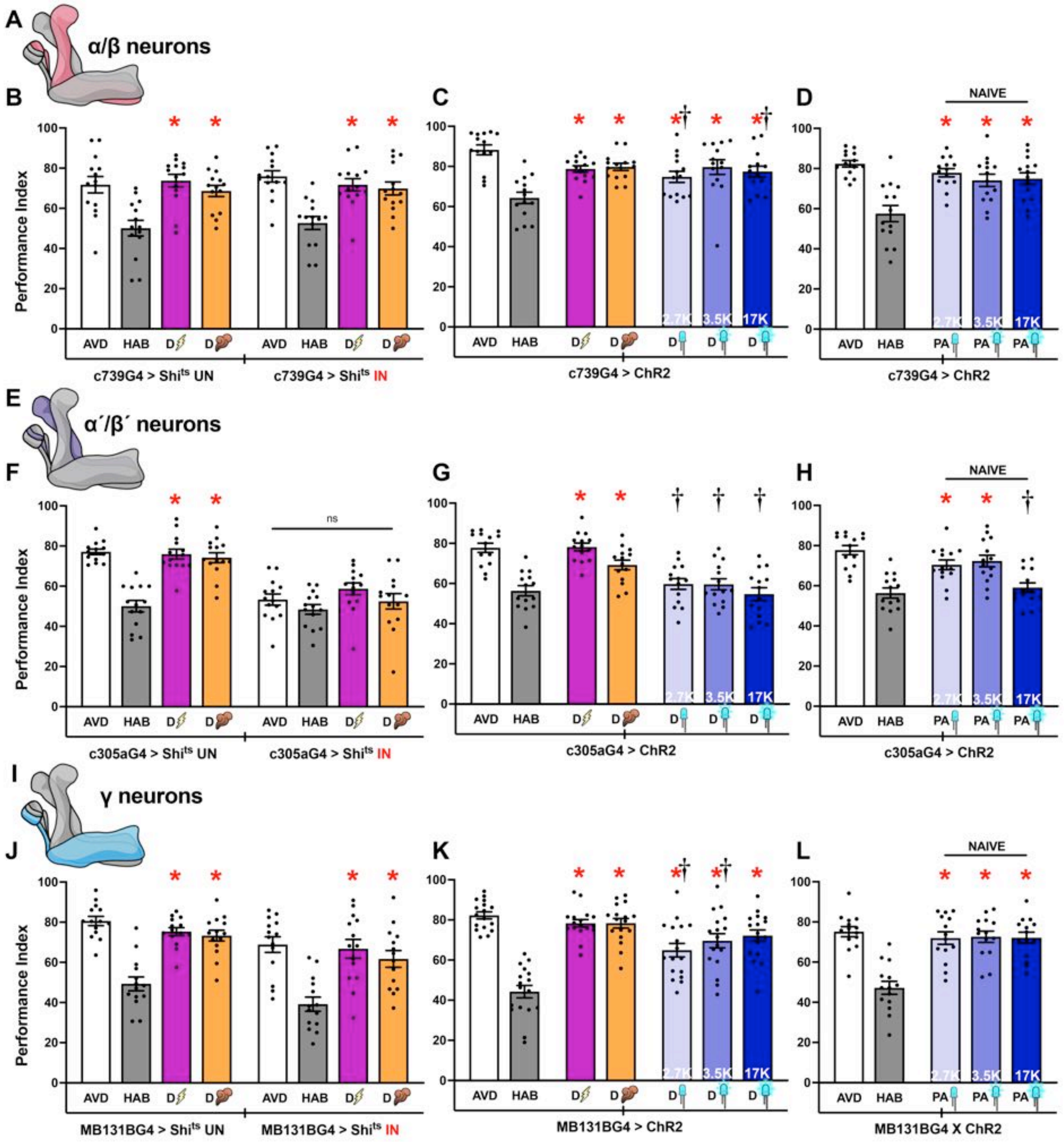

**Supplemental Figure 5: Distinct KCs regulate olfactory habituation and dishabituation.** Mean Performance Indices ± SEM are shown in all figures. The naïve response for OCT is shown in white bars and is always significantly different from the habituated response (grey bars). Cross-modal (footshock) dishabituation is indicated by the bolt (magenta bar) and intramodal (YO), by the plume (orange bar). Exposure to blue light is indicated by the LED bulbs and increasing intensity (as indicated by the lx) by deepening shades of blue. Stars indicate significant differences

($p<0.005$) from the habituated response and daggers significant difference ($p<0.005$) from naïve avoidance. All statistical details are presented in Supplemental Table 1.

**A, E, I)** Schematics of the KC subsets (αβ, α′β′and γ).

**B)** Functional silencing (IN), of αβ neurons with UAS-shibirets under c739Gal4 compared to control animals carrying, but not expressing the transgene (UN). Dishabituation with a 45V electric shock (magenta) and YO (orange), remained unaffected in both conditions. n=14. Additional controls run in parallel are presented in Supplemental Table 1.

**C)** Photoactivation of αβ neurons with 3 light intensities via UAS-ChR2XXL in OCT-habituated flies, restored avoidance of the odorant at the higher light intensities but marginally at 2.7Klx ($p=0.0058$). n=14

**D)** Photoactivation of αβ neurons in naïve flies with 3 light intensities did not affect OCT avoidance. n=14.

**F)** Functional silencing (IN) of α′β′ neurons with UAS-shibirets under c305aGal4 compared to control animals carrying, but not expressing the transgene (UN). Silencing α′β′ MBns results in abrogation of habituation to OCT. n=14. Additional controls run in parallel are presented in Supplemental Table 1.

**G)** Photoactivation of α′β′ neurons under c305aGal4 in OCT-habituated flies with 3 light intensities did not restore avoidance, as a single footshock and YO exposure did. n=14.

**H)** Photoactivation of α′β′ MBns in naïve flies resulted in light intensity dependent decrement in OCT avoidance. n=14.

**J)** Functional silencing (IN) of γ neurons with UAS-shibirets under MB131BGal4 compared to control animals carrying, but not expressing the transgene (UN). Dishabituation with a 45V electric shock (magenta) and YO (orange), remained unaffected in both conditions. n=14. Additional controls run in parallel are presented in Supplemental Table 1.

**K)** Photoactivation of γ neurons under MB131BGal4 in OCT-habituated flies with 3 light intensities reinstated avoidance in a light intensity dependent manner. n=17.

**L)** Photoactivation of γ neurons in naïve flies did not affect OCT avoidance. n=14.

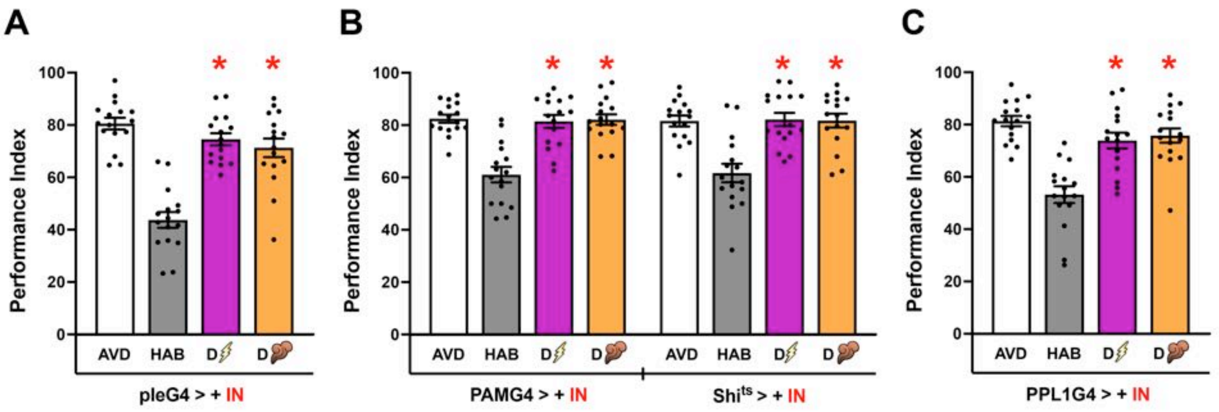

**Supplemental Figure 6: Sibling Controls for Dopaminergic Neurons.**

Mean Performance Indices ± SEM are shown in all figures. The naïve response for OCT is shown in white bars and is always significantly different from the habituated response (grey bars). The dishabituated response is shown in magenta for the footshock dishabituator and in orange for YO. Stars in all Figures indicate significant differences ($p<0.005$), from the habituated response and daggers significant difference ($p<0.005$) from naïve avoidance. All statistical details are presented in Supplemental Table 1.

**A, B, C)** Isogenic Control flies for all DANs (pleGal4>+), for PAM-DANs (0273Gal4>+) and UAS-Shits>+ and PPL1-DANs (MB504BGal4>+) respectively were subjected to the same 32°C induction conditions as the experimental group, but did not present with habituation or dishabituation impairments. n=16 for all genotypes.

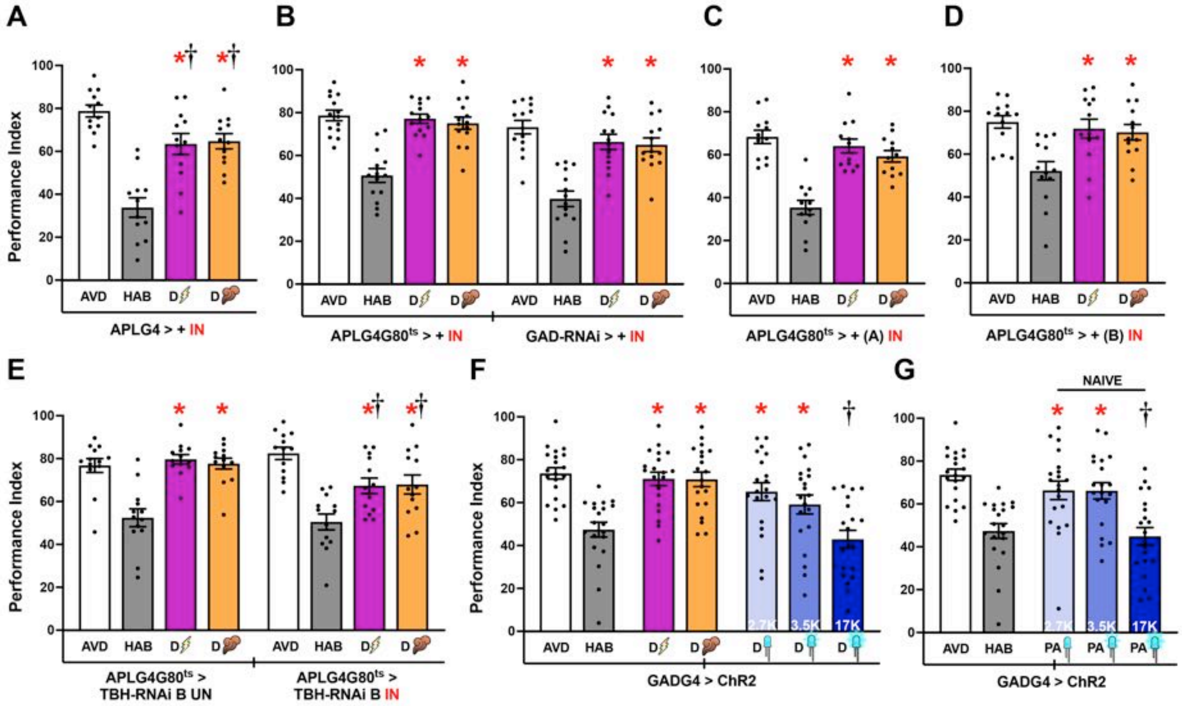

H

| Top **Input** Types | | | | | | | |
|---|---|---|---|---|---|---|---|
| APL (L+R) ♀ | Synapses | Partners | Syns/Partners | APL (L+R) ♂ | Synapses | Partners | Syns/Partners |
| KCαβ | 10.608 | 298 | 35,6 | KCαβ | 45.366 | 980 | 46,3 |
| KCα'β' | 19.919 | 703 | 28,3 | KCα'β' | 15.798 | 334 | 47,3 |
| KCγ | 13.847 | 338 | 41,0 | KCγ | 24.743 | 657 | 37,7 |

| Top **Output** Types | | | | | | | |
|---|---|---|---|---|---|---|---|
| APL (L+R) ♀ | Synapses | Partners | Syns/Partners | APL (L+R) ♂ | Synapses | Partners | Syns/Partners |
| KCαβ | 50.608 | 1.771 | 28,6 | KCαβ | 73.187 | 2.015 | 36,3 |
| KCα'β' | 33.895 | 916 | 37,0 | KCα'β' | 31.002 | 490 | 63,3 |
| KCγ | 62.261 | 2.484 | 25,1 | KCγ | 88.517 | 1.547 | 57,2 |

**Supplemental Figure 7: Isogenic Controls for APL Neurons.**

Mean Performance Indices ± SEM are shown in all figures. The naïve response for OCT is shown in white bars and is always significantly different from the habituated response (grey bars). Cross-modal (footshock) dishabituation is indicated by the bolt (magenta bar) and intramodal (YO), by the plume (orange bar). Exposure to blue light is indicated by the LED bulbs and increasing intensity (as indicated by the lx) by deepening shades of blue. Stars in all Figures indicate significant differences ($p<0.005$), from the habituated response and daggers significant difference ($p<0.005$) from naïve avoidance. All statistical details are presented in Supplemental Table 1.

**A)** Isogenic Control flies APLGal4>+ in $w^{1118}$ genetic background were subjected to the same 32°C induction conditions as the experimental group, did not present habituation or dishabituation impairments. n=12.

**B)** Isogenic Control flies APLGal4;Gal80ts>+ and GAD-RNAi>+ were subjected to the same 30°C induction conditions as the respective experimental group, but habituation or dishabituation were not impaired. n=14.

**C)** Isogenic Control flies APLGal4;Gal80ts>+ with the appropriate genetic background ($y^1v^1$), distinct from that in B, were subjected to the same 30°C induction conditions as the respective experimental group and did not present habituation or dishabituation impairments. n=12.

**D, E)** Functional silencing of APL neurons with another UAS-TBH-RNAi, strain B, under APLGal4;Gal80ts at 30$^o$C, compared to control animals carrying, but not expressing the transgene (UN). Dishabituation with a 45V electric shock (magenta) and YO (orange), although partially reduced, sufficed to reverse OCT habituation. Isogenic Control flies APLGal4;Gal80ts>+ with the appropriate genetic background ($w^{1118}$) dishabituated normally. n=13.

**F)** Photoactivation of GABAergic neurons in OCT-habituated flies restored OCT avoidance at 2.7K and 3.5K light intensities. n=20.

**G)** Photoactivation of GABAergic neurons in naïve flies resulted in light intensity dependent decrement in OCT avoidance. n=20.

H) Total synaptic inputs from KCs to the APL neuron and its synaptic outputs back to KCs, in the female and male Drosophila connectomes. Normalized over the number of synaptic partners the inhibitory effect from APL is stronger to the α′β′ MB neurons.

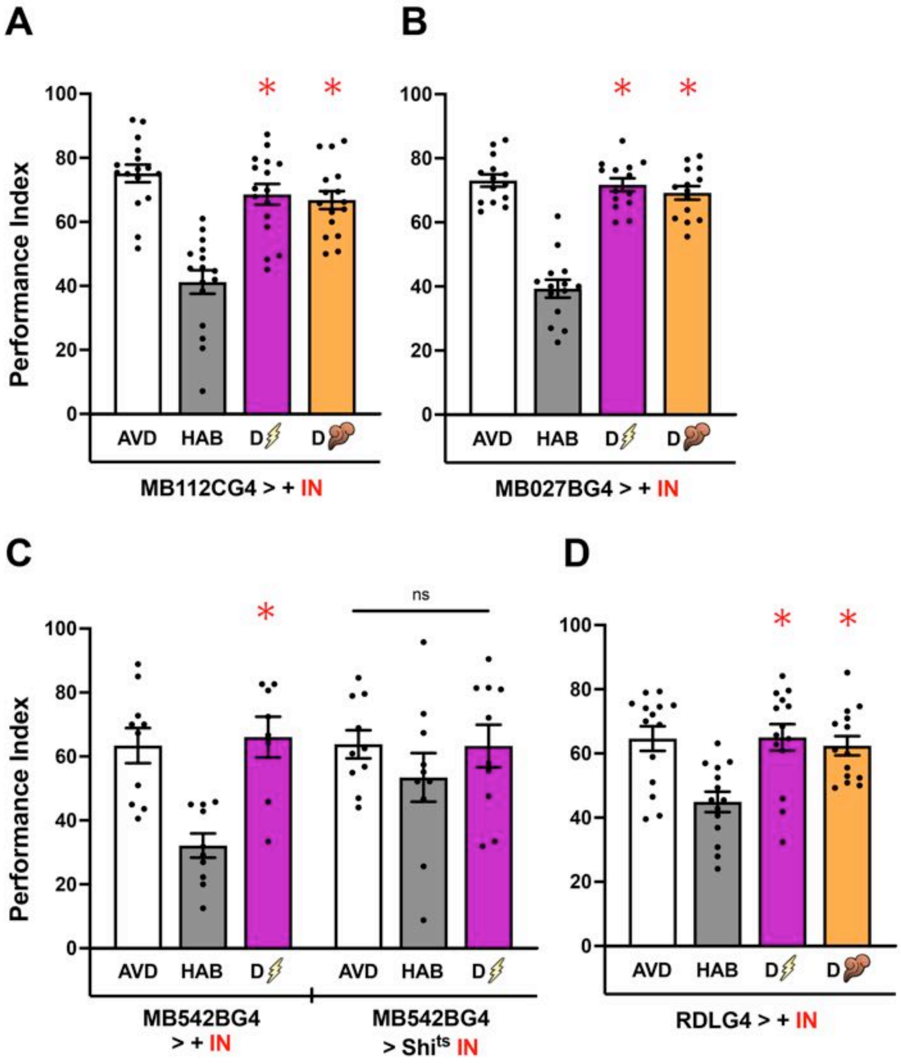

**Supplemental Figure 8: Isogenic Controls for MBONs and Ellipsoid Body Neurons**

Mean Performance Indices ± SEM are shown in all figures. The naïve response is shown in white bars and is always significantly different from the habituated response (grey bars). Cross-modal (footshock) dishabituation is indicated by the bolt (magenta bar). Stars indicate significant differences ($p<0.005$) from the habituated response and daggers significant difference ($p<0.005$) from naïve avoidance. All statistical details are presented in Supplemental Table 1.

**A)** Isogenic Control flies MB112CGal4>+ were subjected to the same 32°C induction conditions as the experimental group, but habituation or dishabituation were not impaired. n=16.

**B)** Isogenic Control flies MB027BGal4>+ were subjected to the same 32°C induction conditions as the experimental group, but habituation or dishabituation were not impaired. n=14.

**C)** Functional silencing (IN) of MBONs 15,16 & 17 with UAS-shibire$^{ts}$ under MB543B-Gal4 compared to control heterozygotes subjected to the same induction conditions. Silencing MBONs 15,16 & 17 results in abrogation of habituation to OCT. n≥8

**D)** Isogenic Control flies RDLGal4>+ were subjected to the same 32°C induction conditions as the experimental group, but habituation or dishabituation were not impaired. n=14.

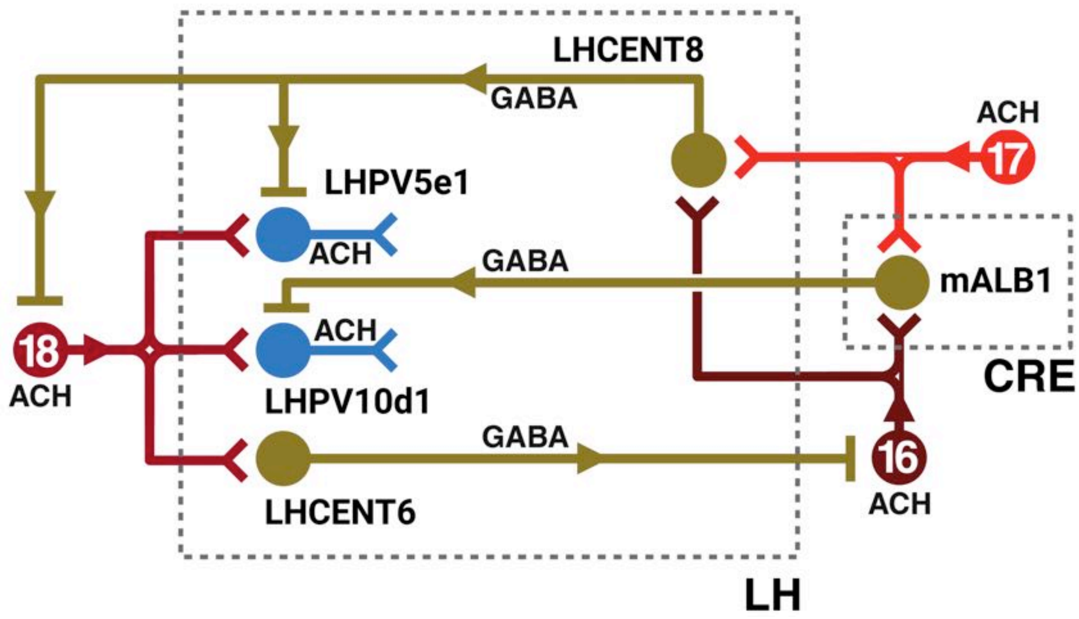

**Supplemental Figure 9: Simplified LH connectome relevant to examined circuitry underlying habituation and dishabituation**

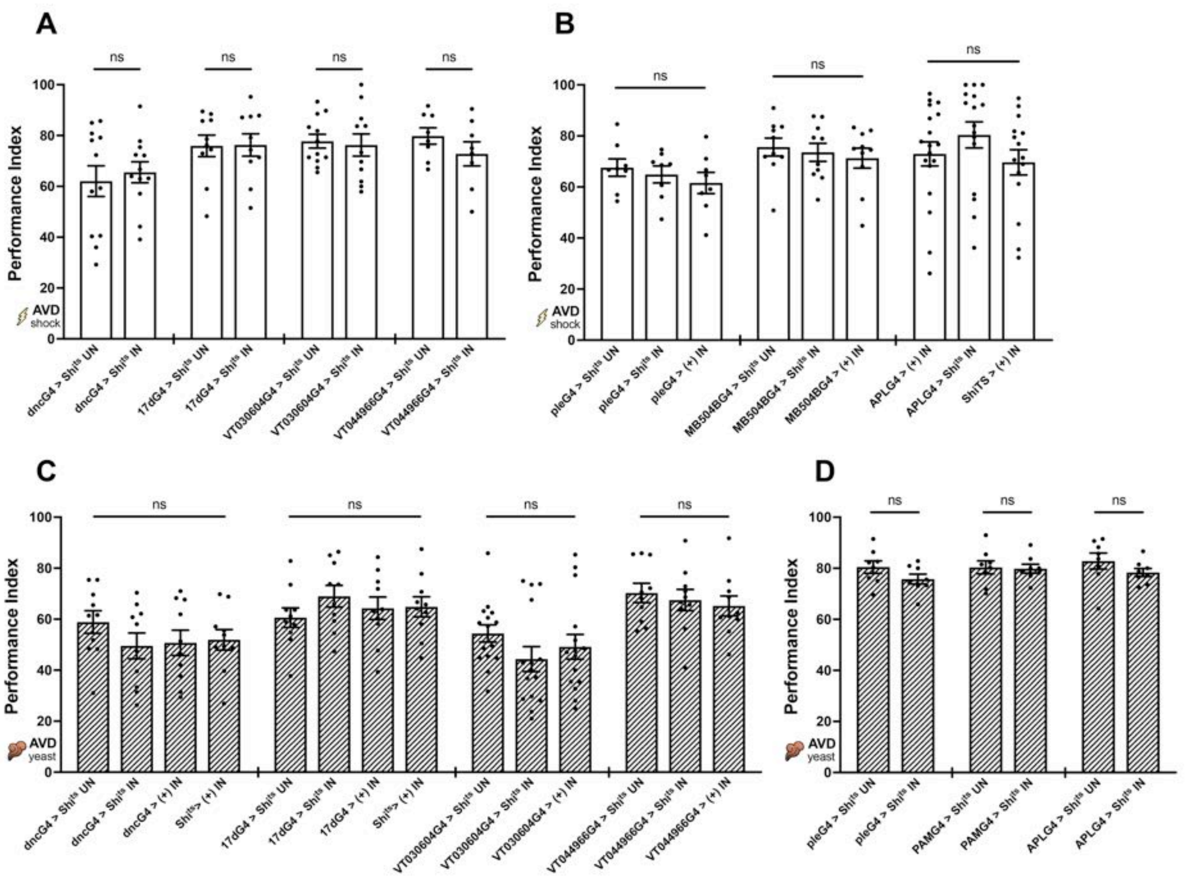

**Supplemental Figure 10: Strains expressing Shibire$^{ts}$ exhibit normal OCT avoidance under permissive and non-permissive conditions.**

Mean Performance Indices ± SEM are shown. The naïve avoidance response (AVD) is shown by the white bar for shock and white striped bar for yeast odor (YO). All statistical details are presented in Supplemental Table 1.

**A)** Blocked neurotransmission in the MB and MBns (αβ, α′β′and γ) did not impair shock avoidance (n=12 for MB, n=10 for α/β, n>11 for α′/β′ and n=8 for γ).

**B)** Blocked neurotransmission in dopaminergic neurons (all & PPL1 subset) and APL neurons did not impair shock avoidance (n=8 for all DANs, n=10 for PPL1 DANs, n>15 for APL).

**C)** Blocked neurotransmission in the MB and MBns (αβ, α′β′and γ) did not impair YO avoidance (n=10 for MB, n=10 for α/β, n=15 for α′/β′ and n=10 for γ).

**D)** Blocked neurotransmission in dopaminergic neurons (all & PAM subset) and APL neurons did not impair YO avoidance (n=8 for all DANs, n=8 for PAM DANs, n=8 for APL).

**DISHABITUATION_PHOTOACTIVATION**

**COLLECTIVE STATISTICS**

| Genotype | Mean ± SEM | HAB Control | | AVD Control | | N |
|---|---|---|---|---|---|---|
| | | F-Ratio | p-value | F-Ratio | p-value | |
| **Figure 2: Olfactory Habituation, intra-modal and cross-modal dishabituation.** | | | | | | |
| **Figure 2A: Canton S Photoactivation (Control)** | | | | | | |
| ANOVA $F_{(3,55)}$=40.6306 **p<0.0001** | | | | | | |
| Canton S AVD Octanol | 72.611 ± 3.395 | 71.673 | **<0.0001** | | | 14 |
| Canton S HAB Octanol | 36.017 ± 3.787 | | | 71.673 | **<0.0001** | 14 |
| Canton S DISHAB **SHOCK** | 77.987 ± 2.250 | 94.276 | **<0.0001** | 1.546 | **0.2192** | 14 |
| Canton S DISHAB **YEAST** | 73.492 ± 2.536 | 75.163 | **<0.0001** | 0.041 | **0.8394** | 14 |
| **Figure 2B: $w^{1118}$ Photoactivation (Control)** | | | | | | |
| ANOVA $F_{(3,47)}$=28.3436 **p<0.0001** | | | | | | |
| $w^{1118}$ AVD Octanol | 70.185 ± 2.744 | 75.300 | **<0.0001** | | | 12 |
| $w^{1118}$ HAB Octanol | 34.607 ± 2.994 | | | 74.479 | **<0.0001** | 12 |
| $w^{1118}$ DISHAB **SHOCK** | 61.847 ± 2.815 | 44.140 | **<0.0001** | 4.136 | **0.0480** | 12 |
| $w^{1118}$ DISHAB **YEAST** | 60.991 ± 3.031 | 41.412 | **<0.0001** | 5.028 | **0.0300** | 12 |
| **Figure 2C: Anosmic Mutants AVOIDANCE $Orco^2$** ($w^{1118}$ background = (+)) | | | | | | |
| ANOVA $F_{(3,47)}$=144.2059 **p<0.0001** | | | | | | |
| $w^{1118}$ AVD Octanol | 75.421 ± 2.685 | - | - | | | 12 |
| $Orco^2$ mutant AVD Octanol | 2.272 ± 2.906 | - | - | 254.394 | **<0.0001** | 12 |
| $w^{1118}$ AVD Yeast | 70.470 ± 2.945 | - | - | | | 12 |
| $Orco^2$ mutant AVD Yeast | 9.260 ± 4.210 | - | - | 178.124 | **<0.0001** | 12 |
| $w^{1118}$ AVD Octanol | 75.421 ± 2.685 | - | - | | | 12 |
| $w^{1118}$ AVD Yeast | 70.470 ± 2.945 | - | - | 1.165 | **0.2861** | 12 |
| $Orco^2$ mutant AVD Octanol | 2.272 ± 2.906 | - | - | | | 12 |
| $Orco^2$ mutant AVD Yeast | 9.260 ± 4.210 | - | - | 2.321 | **0.1347** | 12 |
| **Figure 2D: Anosmic Mutants DISHAB $Orco^2$** ($w^{1118}$ background = (+)) | | | | | | |
| ANOVA $F_{(5,95)}$=54.3988 **p<0.0001** | | | | | | |
| $w^{1118}$ AVD **Shock** | 61.302 ± 1.478 | 103.470 | **<0.0001** | | | 16 |
| $w^{1118}$ HAB **Shock** | 32.736 ± 2.338 | | | 103.470 | **<0.0001** | 16 |
| $w^{1118}$ DISHAB Yeast | 54.249 ± 2.111 | 58.687 | **<0.0001** | 6.306 | **0.0138** | 16 |
| $Orco^2$ mutant AVD **Shock** | 57.584 ± 1.935 | 70.167 | **<0.0001** | | | 16 |
| $Orco^2$ mutant HAB **Shock** | 34.060 ± 1.561 | | | 70.167 | **<0.0001** | 16 |
| $Orco^2$ mutant DISHAB Yeast | 27.866 ± 2.315 | 4.630 | **0.0299** | 111.987 | **<0.0001** | 16 |
| **Figure 2E: $w^{1118}$ Yeast Exposure** | | | | | | |
| ANOVA $F_{(2,38)}$=114.0613 **p<0.0001** | | | | | | |
| $w^{1118}$ AVD Yeast 5 sec | –28.081 ± 4.742 | - | - | | | 13 |
| $w^{1118}$ AVD Yeast 10 sec | 12.182 ± 4.827 | - | - | 42.474 | **<0.0001** | 13 |
| $w^{1118}$ AVD Yeast 90 sec | 64.952 ± 3.384 | - | - | 226.757 | **<0.0001** | 13 |
| **Figure 2F: $w^{1118}$ Octanol Exposure** | | | | | | |
| ANOVA $F_{(2,41)}$=21.5706 **p<0.0001** | | | | | | |
| $w^{1118}$ AVD OCT 5 sec | 26.027 ± 4.823 | - | - | | | 14 |
| $w^{1118}$ AVD OCT 10 sec | 33.865 ± 6.001 | - | - | 1.190 | **0.2819** | 14 |
| $w^{1118}$ AVD OCT 90 sec | 70.244 ± 4.258 | - | - | 37.880 | **<0.0001** | 14 |

| Genotype | Mean ± SEM | HAB Control | | AVD Control | | N |
|---|---|---|---|---|---|---|
| | | F-Ratio | p-value | F-Ratio | p-value | |
| **Figure 2G: w[1118]_HAB(Yeast)_DISHAB(Octanol)** | | | | | | |
| ANOVA $F_{(2,47)}$=33.4736 **p<0.0001** | | | | | | |
| $w^{1118}$ AVD **Yeast** | 73.788 ± 2.849 | 60.038 | **<0.0001** | | | 16 |
| $w^{1118}$ HAB **Yeast** | 39.676 ± 2.684 | | | 60.038 | **<0.0001** | 16 |
| $w^{1118}$ DIS Octanol | 66.753 ± 3.707 | 37.828 | **<0.0001** | 2.553 | **0.1170** | 16 |
| **Figure 2H: Yeast exposures with a 10sec delay prior to Octanol avoidance in w[1118] control flies** | | | | | | |
| ANOVA $F_{(14,192)}$=8.5382 **p<0.0001** | | | | | | |
| $w^{1118}$ AVD Octanol 0.1% OCT | 24.656 ± 3.317 | - | - | | | 14 |
| $w^{1118}$ Exposure **YEAST** 5sec 0.1% OCT | 27.009 ± 3.693 | - | - | 0.155 | **0.6940** | 12 |
| $w^{1118}$ Exposure **YEAST** 15sec 0.1% OCT | 28.651 ± 5.270 | - | - | 0.447 | **0.5043** | 12 |
| $w^{1118}$ AVD Octanol 1% OCT | 33.818 ± 4.759 | - | - | | | 12 |
| $w^{1118}$ Exposure **YEAST** 5sec 1% OCT | 37.036 ± 5.440 | - | - | 0.269 | **0.6041** | 12 |
| $w^{1118}$ Exposure **YEAST** 15sec 1% OCT | 34.045 ± 6.098 | - | - | 0.001 | **0.9707** | 12 |
| $w^{1118}$ AVD Octanol 10% OCT | 36.929 ± 2.920 | - | - | | | 12 |
| $w^{1118}$ Exposure **YEAST** 5sec 10% OCT | 44.912 ± 2.009 | - | - | 1.659 | **0.1993** | 12 |
| $w^{1118}$ Exposure **YEAST** 15sec 10% OCT | 39.440 ± 4.202 | - | - | 0.164 | **0.6857** | 12 |
| $w^{1118}$ AVD Octanol 50% OCT | 52.799 ± 2.576 | - | - | | | 14 |
| $w^{1118}$ Exposure **YEAST** 5sec 50% OCT | 52.507 ± 3.717 | - | - | 0.002 | **0.9609** | 12 |
| $w^{1118}$ Exposure **YEAST** 15sec 50% OCT | 57.504 ± 3.442 | - | - | 0.620 | **0.4317** | 12 |
| $w^{1118}$ AVD Octanol 100% OCT | 60.884 ± 4.125 | - | - | | | 15 |
| $w^{1118}$ Exposure **YEAST** 5sec 100% OCT | 57.052 ± 4.269 | - | - | 0.478 | **0.4902** | 15 |
| $w^{1118}$ Exposure **YEAST** 15sec 100% OCT | 53.168 ± 5.227 | - | - | 1.937 | **0.1656** | 15 |
| **Figure 3: The role of mushroom body neurons in dishabituation.** | | | | | | |
| ***Figure 3B, 3E: dncG4 > Shi[ts]** ($w^{1118}$ background = (+)) | | | | | | |
| ANOVA $F_{(15,223)}$=22.8621 **p<0.0001** | | | | | | |
| dncG4 > (+) **IN** AVD Octanol | 69.622 ± 2.549 | 54.072 | **<0.0001** | | | 14 |
| dncG4 > (+) **IN** HAB Octanol | 37.188 ± 3.354 | | | 54.072 | **<0.0001** | 14 |
| dncG4 > (+) **IN** DISHAB **shock** | 69.316 ± 3.235 | 53.056 | **<0.0001** | 0.004 | **0.9447** | 14 |
| dncG4 > (+) **IN** DISHAB **yeast** | 70.523 ± 3.179 | 57.119 | **<0.0001** | 0.041 | **0.8382** | 14 |
| dncG4 > $Shi^{ts}$ **IN** AVD Octanol | 67.673 ± 3.395 | 37.587 | **<0.0001** | | | 14 |
| dncG4 > $Shi^{ts}$ **IN** HAB Octanol | 40.631 ± 3.483 | | | 37.587 | **<0.0001** | 14 |
| dncG4 > $Shi^{ts}$ **IN** DISHAB **shock** | 42.657 ± 3.518 | 0.211 | **0.6464** | 32.166 | **<0.0001** | 14 |
| dncG4 > $Shi^{ts}$ **IN** DISHAB **yeast** | 43.330 ± 3.045 | 0.374 | **0.5413** | 30.459 | **<0.0001** | 14 |
| dncG4 > $Shi^{ts}$ UN AVD Octanol | 70.842 ± 3.523 | 46.780 | **<0.0001** | | | 14 |
| dncG4 > $Shi^{ts}$ UN HAB Octanol | 40.674 ± 2.779 | | | 46.780 | **<0.0001** | 14 |
| dncG4 > $Shi^{ts}$ UN DISHAB **shock** | 72.339 ± 3.006 | 51.538 | **<0.0001** | 0.115 | **0.7346** | 14 |
| dncG4 > $Shi^{ts}$ UN DISHAB **yeast** | 72.358 ± 3.044 | 51.599 | **<0.0001** | 0.118 | **0.7314** | 14 |
| $Shi^{ts}$ > (+) **IN** AVD Octanol | 74.383 ± 2.674 | 35.084 | **<0.0001** | | | 14 |
| $Shi^{ts}$ > (+) **IN** HAB Octanol | 48.257 ± 3.292 | | | 35.084 | **<0.0001** | 14 |
| $Shi^{ts}$ > (+) **IN** DISHAB **shock** | 73.385 ± 2.708 | 32.454 | **<0.0001** | 0.051 | **0.8211** | 14 |
| $Shi^{ts}$ > (+) **IN** DISHAB **yeast** | 73.998 ± 2.867 | 34.057 | **<0.0001** | 0.007 | **0.9305** | 14 |
| **Figure 3C: dncG4 > UAS-ChR2-XXL4 Channel Rhodopsin (ChR2)** | | | | | | |
| ANOVA $F_{(6,111)}$=8.8711 **p<0.0001** | | | | | | |
| dncG4 > ChR2 **IN** AVD Octanol | 77.777 ± 3.498 | 33.454 | **<0.0001** | | | 16 |
| dncG4 > ChR2 **IN** HAB Octanol | 51.199 ± 3.660 | | | 33.454 | **<0.0001** | 16 |
| dncG4 > ChR2 **IN** DISHAB **SHOCK** | 75.498 ± 2.751 | 27.962 | **<0.0001** | 0.246 | **0.6209** | 16 |
| dncG4 > ChR2 **IN** DISHAB **YEAST** | 78.497 ± 3.214 | 35.292 | **<0.0001** | 0.024 | **0.8757** | 16 |

| Genotype | Mean ± SEM | HAB Control | | AVD Control | | N |
|---|---|---|---|---|---|---|
| | | F-Ratio | p-value | F-Ratio | p-value | |
| dncG4 > ChR2 **IN DISHAB 2.7K** Photo | 76.111 ± 2.648 | 29.390 | **<0.0001** | 0.131 | **0.7175** | 16 |
| dncG4 > ChR2 **IN DISHAB 3.5K** Photo | 71.715 ± 3.141 | 19.934 | **<0.0001** | 1.740 | **0.1899** | 16 |
| dncG4 > ChR2 **IN DISHAB 17K** Photo | 66.431 ± 3.670 | 10.988 | **0.0012** | 6.096 | **0.0151** | 16 |
| **Figure 3D: dncG4 > UAS-ChR2-XXL4 Channel Rhodopsin (ChR2) NAIVE** | | | | | | |
| ANOVA $F_{(4,79)}$=11.6302 **p<0.0001** | | | | | | |
| dncG4 > ChR2 **IN** AVD Octanol | 77.777 ± 3.498 | 32.108 | **<0.0001** | | | 16 |
| dncG4 > ChR2 **IN** HAB Octanol | 51.199 ± 3.660 | | | 32.108 | **<0.0001** | 16 |
| dncG4 > ChR2 **IN PHOTO 2.7K** | 78.994 ± 2.694 | 35.115 | **<0.0001** | 0.067 | **0.7960** | 16 |
| dncG4 > ChR2 **IN PHOTO 3.5K** | 73.885 ± 3.271 | 23.393 | **<0.0001** | 0.688 | **0.4093** | 16 |
| dncG4 > ChR2 **IN PHOTO 17K** | 68.645 ± 3.375 | 13.833 | **<0.0004** | 3.791 | **0.0552** | 16 |
| **Figure 3F: dncG4 > $W^{1118}$ Photoactivation Control** | | | | | | |
| ANOVA $F_{(6,111)}$=24.4172 **p<0.0001** | | | | | | |
| dncG4 > (+) **IN** AVD Octanol | 80.615 ± 2.201 | 57.138 | **<0.0001** | | | 16 |
| dncG4 > (+) **IN** HAB Octanol | 49.824 ± 3.343 | | | 57.138 | **<0.0001** | 16 |
| dncG4 > (+) **IN** DISHAB **SHOCK** | 78.758 ± 2.698 | 50.453 | **<0.0001** | 0.207 | **0.6493** | 16 |
| dncG4 > (+) **IN** DISHAB **YEAST** | 78.791 ± 2.505 | 50.571 | **<0.0001** | 0.200 | **0.6553** | 16 |
| dncG4 > (+) **IN DISHAB 2.7K** Photo | 53.445 ± 2.804 | 0.790 | **0.3760** | 44.490 | **<0.0001** | 16 |
| dncG4 > (+) **IN DISHAB 3.5K** Photo | 54.274 ± 3.477 | 1.193 | **0.2771** | 41.815 | **<0.0001** | 16 |
| dncG4 > (+) **IN DISHAB 17K** Photo | 54.184 ± 2.920 | 1.145 | **0.2868** | 42.102 | **<0.0001** | 16 |
| **Figure 3G: dncG4 > $W^{1118}$ Photoactivation Control NAIVE** | | | | | | |
| ANOVA $F_{(4,79)}$=27.6518 **p<0.0001** | | | | | | |
| dncG4 > (+) **IN** AVD Octanol | 80.615 ± 2.201 | 77.524 | **<0.0001** | | | 16 |
| dncG4 > (+) **IN** HAB Octanol | 49.824 ± 3.343 | | | 77.524 | **<0.0001** | 16 |
| dncG4 > (+) **IN PHOTO 2.7K** | 76.594 ± 2.290 | 58.598 | **<0.0001** | 1.322 | **0.2538** | 16 |
| dncG4 > (+) **IN PHOTO 3.5K** | 78.748 ± 2.065 | 68.410 | **<0.0001** | 0.284 | **0.5951** | 16 |
| dncG4 > (+) **IN PHOTO 17K** | 78.939 ± 2.243 | 69.316 | **<0.0001** | 0.229 | **0.6332** | 16 |
| **Figure 4: Distinct MBns regulate olfactory habituation and dishabituation.** | | | | | | |
| **Figure 4B: (α/β)17dG4 > $Shi^{ts}$** ($w^{1118}$ background = (+)) | | | | | | |
| ANOVA $F_{(11,143)}$=13.8190 **p<0.0001** | | | | | | |
| (α/β)17dG4 > (+) **IN** AVD Octanol | 77.272 ± 3.553 | 42.787 | **<0.0001** | | | 12 |
| (α/β)17dG4 > (+) **IN** HAB Octanol | 49.258 ± 3.752 | | | 42.787 | **<0.0001** | 12 |
| (α/β)17dG4 > (+) **IN** DISHAB **shock** | 71.125 ± 2.897 | 26.071 | **<0.0001** | 2.059 | **0.1535** | 12 |
| (α/β)17dG4 > (+) **IN** DISHAB **yeast** | 72.621 ± 3.127 | 29.759 | **<0.0001** | 1.179 | **0.2794** | 12 |
| (α/β)17dG4 > $Shi^{ts}$ **IN** AVD Octanol | 80.911 ± 2.526 | 36.255 | **<0.0001** | | | 12 |
| (α/β)17dG4 > $Shi^{ts}$ **IN** HAB Octanol | 55.124 ± 2.581 | | | 36.255 | **<0.0001** | 12 |
| (α/β)17dG4 > $Shi^{ts}$ **IN** DISHAB **shock** | 82.580 ± 3.524 | 41.097 | **<0.0001** | 0.151 | **0.6975** | 12 |
| (α/β)17dG4 > $Shi^{ts}$ **IN** DISHAB **yeast** | 74.986 ± 3.223 | 21.506 | **<0.0001** | 1.914 | **0.1687** | 12 |
| (α/β)17dG4 > $Shi^{ts}$ UN AVD Octanol | 81.019 ± 2.611 | 27.764 | **<0.0001** | | | 12 |
| (α/β)17dG4 > $Shi^{ts}$ UN HAB Octanol | 58.452 ± 3.640 | | | 27.764 | **<0.0001** | 12 |
| (α/β)17dG4 > $Shi^{ts}$ UN DISHAB **shock** | 80.491 ± 2.441 | 26.481 | **<0.0001** | 0.015 | **0.9021** | 12 |
| (α/β)17dG4 > $Shi^{ts}$ UN DISHAB **yeast** | 77.159 ± 1.809 | 19.079 | **<0.0001** | 0.812 | **0.3691** | 12 |
| **Figure 4C: (α/β)17dG4 > UAS-ChR2-XXL4 Channel Rhodopsin (ChR2)** | | | | | | |
| ANOVA $F_{(6,97)}$=6.1016 **p<0.0001** | | | | | | |
| (α/β)17dG4 > ChR2 **IN** AVD Octanol | 73.465 ± 4.323 | 21.439 | **<0.0001** | | | 14 |
| (α/β)17dG4 > ChR2 **IN** HAB Octanol | 47.366 ± 3.379 | | | 21.439 | **<0.0001** | 14 |

| Genotype | Mean ± SEM | HAB Control | | AVD Control | | N |
|---|---|---|---|---|---|---|
| | | F-Ratio | p-value | F-Ratio | p-value | |
| (α/β)17dG4 > ChR2 **IN** DISHAB **SHOCK** | 72.952 ± 3.616 | 20.605 | **<0.0001** | 0.008 | **0.9277** | 14 |
| (α/β)17dG4 > ChR2 **IN** DISHAB **YEAST** | 69.593 ± 3.268 | 15.550 | **<0.0002** | 0.471 | **0.4938** | 14 |
| (α/β)17dG4 > ChR2 **IN DISHAB 2.7K** Photo | 75.650 ± 4.941 | 25.179 | **<0.0001** | 0.150 | **0.6991** | 14 |
| (α/β)17dG4 > ChR2 **IN DISHAB 3.5K** Photo | 74.726 ± 4.313 | 23.561 | **<0.0001** | 0.050 | **0.8234** | 14 |
| (α/β)17dG4 > ChR2 **IN DISHAB 17K** Photo | 68.103 ± 3.779 | 13.534 | **<0.0004** | 0.905 | **0.3439** | 14 |
| **Figure 4D: (α/β)17dG4 > UAS-ChR2-XXL4 Channel Rhodopsin (ChR2) NAIVE** | | | | | | |
| ANOVA $F_{(4,69)}$=8.5600 **p<0.0001** | | | | | | |
| (α/β)17dG4 > ChR2 **IN** AVD Octanol | 73.465 ± 4.323 | 23.641 | **<0.0001** | | | 14 |
| (α/β)17dG4 > ChR2 **IN** HAB Octanol | 47.366 ± 3.379 | | | 23.641 | **<0.0001** | 14 |
| (α/β)17dG4 > ChR2 **IN PHOTO 2.7K** | 72.144 ± 3.869 | 21.309 | **<0.0001** | 0.060 | **0.8064** | 14 |
| (α/β)17dG4 > ChR2 **IN PHOTO 3.5K** | 71.886 ± 3.578 | 20.867 | **<0.0001** | 0.086 | **0.7695** | 14 |
| (α/β)17dG4 > ChR2 **IN PHOTO 17K** | 70.970 ± 3.758 | 19.338 | **<0.0001** | 0.215 | **0.6436** | 14 |
| **Figure 4F: (α′/β′)VT030604G4 > Shi$^{ts}$** (w$^{1118}$ background = (+)) | | | | | | |
| ANOVA $F_{(11,167)}$=10.5032 **p<0.0001** | | | | | | |
| (α′/β′)VT030604G4 > (+) **IN** AVD Octanol | 78.669 ± 3.369 | 25.523 | **<0.0001** | | | 14 |
| (α′/β′)VT030604G4 > (+) **IN** HAB Octanol | 54.568 ± 3.209 | | | 25.523 | **<0.0001** | 14 |
| (α′/β′)VT030604G4 > (+) **IN** DISHAB **shock** | 70.436 ± 3.646 | 11.063 | **0.0011** | 2.978 | **0.0863** | 14 |
| (α′/β′)VT030604G4 > (+) **IN** DISHAB **yeast** | 70.144 ± 3.837 | 10.660 | **0.0013** | 3.193 | **0.0758** | 14 |
| (α′/β′)VT030604G4 > Shi$^{ts}$ **IN** AVD Octanol | 57.102 ± 4.023 | 0.714 | **0.3993** | | | 14 |
| (α′/β′)VT030604G4 > Shi$^{ts}$ **IN** HAB Octanol | 53.070 ± 4.497 | | | 0.714 | **0.3993** | 14 |
| (α′/β′)VT030604G4 > Shi$^{ts}$ **IN** DISHAB **shock** | 53.605 ± 3.481 | 0.012 | **0.9108** | 0.537 | **0.4646** | 14 |
| (α′/β′)VT030604G4 > Shi$^{ts}$ **IN** DISHAB **yeast** | 51.258 ± 3.461 | 0.144 | **0.7045** | 1.500 | **0.2224** | 14 |
| (α′/β′)VT030604G4 > Shi$^{ts}$ UN AVD Octanol | 72.581 ± 2.638 | 29.074 | **<0.0001** | | | 14 |
| (α′/β′)VT030604G4 > Shi$^{ts}$ UN HAB Octanol | 46.857 ± 2.816 | | | 29.074 | **<0.0001** | 14 |
| (α′/β′)VT030604G4 > Shi$^{ts}$ UN DISHAB **shock** | 72.848 ± 2.383 | 29.679 | **<0.0001** | 0.003 | **0.9555** | 14 |
| (α′/β′)VT030604G4 > Shi$^{ts}$ UN DISHAB **yeast** | 72.140 ± 2.402 | 28.085 | **<0.0001** | 0.008 | **0.9264** | 14 |
| **Figure 4G: (α′/β′)VT030604G4 > UAS-ChR2-XXL4 Channel Rhodopsin (ChR2)** | | | | | | |
| ANOVA $F_{(6,83)}$=24.2353 **p<0.0001** | | | | | | |
| (α′/β′)VT030604G4 > ChR2 **IN** AVD Octanol | 80.163 ± 2.407 | 67.935 | **<0.0001** | | | 12 |
| (α′/β′)VT030604G4 > ChR2 **IN** HAB Octanol | 42.560 ± 2.437 | | | 67.935 | **<0.0001** | 12 |
| (α′/β′)VT030604G4 > ChR2 **IN** DISHAB **SHOCK** | 65.385 ± 2.895 | 25.028 | **<0.0001** | 10.493 | **0.0017** | 12 |
| (α′/β′)VT030604G4 > ChR2 **IN** DISHAB **YEAST** | 63.690 ± 2.754 | 21.450 | **<0.0001** | 13.037 | **<0.0006** | 12 |
| (α′/β′)VT030604G4 > ChR2 **IN DISHAB 2.7K** Photo | 45.484 ± 3.519 | 0.410 | **0.5235** | 57.781 | **<0.0001** | 12 |
| (α′/β′)VT030604G4 > ChR2 **IN DISHAB 3.5K** Photo | 40.043 ± 4.337 | 0.304 | **0.5827** | 77.333 | **<0.0001** | 12 |
| (α′/β′)VT030604G4 > ChR2 **IN DISHAB 17K** Photo | 39.604 ± 3.733 | 0.419 | **0.5189** | 79.037 | **<0.0001** | 12 |
| **Figure 4H: (α′/β′)VT030604G4 > UAS-ChR2-XXL4 Channel Rhodopsin (ChR2) NAIVE** | | | | | | |
| ANOVA $F_{(4,59)}$=23.5507 **p<0.0001** | | | | | | |
| (α′/β′)VT030604G4 > ChR2 **IN** AVD Octanol | 80.163 ± 2.407 | 67.055 | **<0.0001** | | | 12 |
| (α′/β′)VT030604G4 > ChR2 **IN** HAB Octanol | 42.560 ± 2.437 | | | 67.055 | **<0.0001** | 12 |
| (α′/β′)VT030604G4 > ChR2 **IN PHOTO 2.7K** | 67.971 ± 3.761 | 30.621 | **<0.0001** | 7.049 | **0.0103** | 12 |
| (α′/β′)VT030604G4 > ChR2 **IN PHOTO 3.5K** | 56.239 ± 4.385 | 8.873 | **0.0042** | 27.142 | **<0.0001** | 12 |
| (α′/β′)VT030604G4 > ChR2 **IN PHOTO 17K** | 45.269 ± 2.756 | 0.347 | **0.5576** | 57.742 | **<0.0001** | 12 |
| **Figure 4J: (γ)VT044966G4 > Shi$^{ts}$** (w$^{1118}$ background = (+)) | | | | | | |
| ANOVA $F_{(11,143)}$=15.6053 **p<0.0001** | | | | | | |
| (γ)VT044966G4 > (+) **IN** AVD Octanol | 77.584 ± 2.896 | 42.873 | **<0.0001** | | | 12 |

| Genotype | Mean ± SEM | HAB Control | | AVD Control | | N |
|---|---|---|---|---|---|---|
| | | F-Ratio | p-value | F-Ratio | p-value | |
| (γ)VT044966G4 > (+) **IN** HAB Octanol | 50.137 ± 2.524 | | | 42.873 | **<0.0001** | 12 |
| (γ)VT044966G4 > (+) **IN** DISHAB **shock** | 72.758 ± 3.128 | 29.121 | **<0.0001** | 1.325 | **0.2516** | 12 |
| (γ)VT044966G4 > (+) **IN** DISHAB **yeast** | 80.932 ± 2.096 | 53.971 | **<0.0001** | 0.638 | **0.4258** | 12 |
| (γ)VT044966G4 > Shi$^{ts}$ **IN** AVD Octanol | 83.046 ± 3.043 | 51.910 | **<0.0001** | | | 12 |
| (γ)VT044966G4 > Shi$^{ts}$ **IN** HAB Octanol | 52.844 ± 3.204 | | | 51.910 | **<0.0001** | 12 |
| (γ)VT044966G4 > Shi$^{ts}$ **IN** DISHAB **shock** | 80.590 ± 3.295 | 43.808 | **<0.0001** | 0.3434 | **0.5588** | 12 |
| (γ)VT044966G4 > Shi$^{ts}$ **IN** DISHAB **yeast** | 77.315 ± 2.904 | 34.079 | **<0.0001** | 1.869 | **0.1739** | 12 |
| (γ)VT044966G4 > Shi$^{ts}$ UN AVD Octanol | 79.892 ± 2.573 | 23.903 | **<0.0001** | | | 12 |
| (γ)VT044966G4 > Shi$^{ts}$ UN HAB Octanol | 59.398 ± 3.578 | | | 23.903 | **<0.0001** | 12 |
| (γ)VT044966G4 > Shi$^{ts}$ UN DISHAB **shock** | 79.427 ± 3.481 | 22.830 | **<0.0001** | 0.0123 | **0.9118** | 12 |
| (γ)VT044966G4 > Shi$^{ts}$ UN DISHAB **yeast** | 79.684 ± 2.470 | 23.420 | **<0.0001** | 0.002 | **0.9604** | 12 |
| **Figure 4K: (γ)VT044966G4 > UAS-ChR2-XXL4 Channel Rhodopsin (ChR2)** | | | | | | |
| ANOVA $F_{(6,97)}$=13.7973 **p<0.0001** | | | | | | |
| (γ)VT044966G4 > ChR2 **IN** AVD Octanol | 70.144 ± 3.500 | 55.361 | **<0.0001** | | | 14 |
| (γ)VT044966G4 > ChR2 **IN** HAB Octanol | 37.393 ± 4.156 | | | 55.361 | **<0.0001** | 14 |
| (γ)VT044966G4 > ChR2 **IN** DISHAB **SHOCK** | 63.619 ± 2.430 | 35.498 | **<0.0001** | 2.197 | **0.1416** | 14 |
| (γ)VT044966G4 > ChR2 **IN** DISHAB **YEAST** | 65.862 ± 2.146 | 41.830 | **<0.0001** | 0.946 | **0.3331** | 14 |
| (γ)VT044966G4 > ChR2 **IN DISHAB 2.7K** Photo | 47.114 ± 3.059 | 4.877 | **0.0297** | 27.374 | **<0.0001** | 14 |
| (γ)VT044966G4 > ChR2 **IN DISHAB 3.5K** Photo | 59.362 ± 2.895 | 24.910 | **<0.0001** | 5.999 | **0.0162** | 14 |
| (γ)VT044966G4 > ChR2 **IN DISHAB 17K** Photo | 62.018 ± 3.165 | 31.297 | **<0.0001** | 3.408 | **0.0681** | 14 |
| **Figure 4L: (γ)VT044966G4 > UAS-ChR2-XXL4 Channel Rhodopsin (ChR2) NAIVE** | | | | | | |
| ANOVA $F_{(4,69)}$=10.9439 **p<0.0001** | | | | | | |
| (γ)VT044966G4 > ChR2 **IN** AVD Octanol | 70.144 ± 3.500 | 40.050 | **<0.0001** | | | 14 |
| (γ)VT044966G4 > ChR2 **IN** HAB Octanol | 37.393 ± 4.156 | | | 40.050 | **<0.0001** | 14 |
| (γ)VT044966G4 > ChR2 **IN PHOTO 2.7K** | 61.602 ± 2.220 | 21.883 | **<0.0001** | 2.724 | **0.1036** | 14 |
| (γ)VT044966G4 > ChR2 **IN PHOTO 3.5K** | 59.814 ± 4.318 | 18.769 | **<0.0001** | 3.984 | **0.0501** | 14 |
| (γ)VT044966G4 > ChR2 **IN PHOTO 17K** | 57.633 ± 3.721 | 15.296 | **<0.0003** | 5.844 | **0.0184** | 14 |
| **Figure 4N: (α/β)17dG4 ; (γ)VT044966G4 > Shi$^{ts}$** ($w^{1118}$ background = (+)) | | | | | | |
| ANOVA $F_{(11,215)}$=28.9779 **p<0.0001** | | | | | | |
| (α/β)17dG4 ; (γ)VT044966G4 > (+) **IN** AVD Octanol | 70.402 ± 2.114 | 75.474 | **<0.0001** | | | 18 |
| (α/β)17dG4 ; (γ)VT044966G4 > (+) **IN** HAB Octanol | 38.114 ± 2.917 | | | 75.474 | **<0.0001** | 18 |
| (α/β)17dG4 ; (γ)VT044966G4 > (+) **IN** DISHAB **shock** | 65.030 ± 2.611 | 52.449 | **<0.0001** | 2.089 | **0.1498** | 18 |
| (α/β)17dG4 ; (γ)VT044966G4 > (+) **IN** DISHAB **yeast** | 63.754 ± 2.501 | 47.592 | **<0.0001** | 3.200 | **0.0751** | 18 |
| (α/β)17dG4 ; (γ)VT044966G4 > Shi$^{ts}$ **IN** AVD Octanol | 72.872 ± 2.705 | 75.568 | **<0.0001** | | | 18 |
| (α/β)17dG4 ; (γ)VT044966G4 > Shi$^{ts}$ **IN** HAB Octanol | 40.563 ± 3.328 | | | 75.568 | **<0.0001** | 18 |
| (α/β)17dG4 ; (γ)VT044966G4 > Shi$^{ts}$ **IN** DISHAB **shock** | 46.618 ± 2.915 | 2.653 | **0.1048** | 49.899 | **<0.0001** | 18 |
| (α/β)17dG4 ; (γ)VT044966G4 > Shi$^{ts}$ **IN** DISHAB **yeast** | 47.263 ± 2.553 | 3.250 | **0.0728** | 47.475 | **<0.0001** | 18 |
| (α/β)17dG4 ; (γ)VT044966G4 > Shi$^{ts}$ UN AVD Octanol | 67.042 ± 1.976 | 93.155 | **<0.0001** | | | 18 |
| (α/β)17dG4 ; (γ)VT044966G4 > Shi$^{ts}$ UN HAB Octanol | 31.170 ± 2.514 | | | 93.155 | **<0.0001** | 18 |
| (α/β)17dG4 ; (γ)VT044966G4 > Shi$^{ts}$ UN DISHAB **shock** | 65.133 ± 2.543 | 83.505 | **<0.0001** | 0.263 | **0.6080** | 18 |
| (α/β)17dG4 ; (γ)VT044966G4 > Shi$^{ts}$ UN DISHAB **yeast** | 60.917 ± 2.590 | 64.057 | **<0.0001** | 2.716 | **0.1008** | 18 |

| Genotype | Mean ± SEM | HAB Control | | AVD Control | | N |
|---|---|---|---|---|---|---|
| | | F-Ratio | p-value | F-Ratio | p-value | |
| **Figure 5: Dopaminergic Neurons drive intra-modal and cross-modal dishabituation to Octanol in Drosophila.** | | | | | | |
| **Figure 5D: PLEG4 > Shi$^{ts}$** (w$^{1118}$ background = (+)) | | | | | | |
| ANOVA $F_{(11,191)}$=20.6590 **p<0.0001** | | | | | | |
| PLEG4 > (+) **IN** AVD Octanol | 80.520 ± 2.247 | 66.016 | **<0.0001** | | | 16 |
| PLEG4 > (+) **IN** HAB Octanol | 43.725 ± 3.021 | | | 66.016 | **<0.0001** | 16 |
| PLEG4 > (+) **IN** DISHAB **shock** | 74.573 ± 2.293 | 46.399 | **<0.0001** | 1.724 | **0.1907** | 16 |
| PLEG4 > (+) **IN** DISHAB **yeast** | 71.287 ± 3.553 | 37.042 | **<0.0001** | 4.156 | **0.0429** | 16 |
| PLEG4 > Shi$^{ts}$ **IN** AVD Octanol | 78.308 ± 2.388 | 37.064 | **<0.0001** | | | 16 |
| PLEG4 > Shi$^{ts}$ **IN** HAB Octanol | 50.737 ± 3.183 | | | 37.064 | **<0.0001** | 16 |
| PLEG4 > Shi$^{ts}$ **IN** DISHAB **shock** | 49.730 ± 3.428 | 0.049 | **0.8242** | 39.822 | **<0.0001** | 16 |
| PLEG4 > Shi$^{ts}$ **IN** DISHAB **yeast** | 50.719 ± 3.845 | <0.001 | **0.9968** | 37.112 | **<0.0001** | 16 |
| PLEG4 > Shi$^{ts}$ UN AVD Octanol | 80.284 ± 2.768 | 56.624 | **<0.0001** | | | 16 |
| PLEG4 > Shi$^{ts}$ UN HAB Octanol | 46.206 ± 3.296 | | | 56.624 | **<0.0001** | 16 |
| PLEG4 > Shi$^{ts}$ UN DISHAB **shock** | 71.805 ± 3.964 | 31.953 | **<0.0001** | 3.505 | **0.0628** | 16 |
| PLEG4 > Shi$^{ts}$ UN DISHAB **yeast** | 72.871 ± 3.787 | 34.669 | **<0.0001** | 2.679 | **0.1034** | 16 |
| **Figure 5E: PLEG4 > UAS-ChR2-XXL4 Channel Rhodopsin (ChR2)** | | | | | | |
| ANOVA $F_{(6,111)}$=12.2747 **p<0.0001** | | | | | | |
| PLEG4 > ChR2 **IN** AVD Octanol | 82.690 ± 2.111 | 49.838 | **<0.0001** | | | 16 |
| PLEG4 > ChR2 **IN** HAB Octanol | 55.131 ± 2.363 | | | 49.838 | **<0.0001** | 16 |
| PLEG4 > ChR2 **IN** DISHAB **SHOCK** | 76.044 ± 2.695 | 28.698 | **<0.0001** | 2.898 | **0.0916** | 16 |
| PLEG4 > ChR2 **IN** DISHAB **YEAST** | 73.806 ± 2.879 | 22.886 | **<0.0001** | 5.178 | **0.0248** | 16 |
| PLEG4 > ChR2 **IN DISHAB 2.7K** Photo | 58.835 ± 3.699 | 0.899 | **0.3449** | 37.344 | **<0.0001** | 16 |
| PLEG4 > ChR2 **IN DISHAB 3.5K** Photo | 71.939 ± 2.885 | 18.536 | **<0.0001** | 7.585 | **0.0069** | 16 |
| PLEG4 > ChR2 **IN DISHAB 17K** Photo | 69.787 ± 2.390 | 14.095 | **<0.0003** | 10.925 | **0.0012** | 16 |
| **Figure 5F: PLEG4 > UAS-ChR2-XXL4 Channel Rhodopsin (ChR2) NAIVE** | | | | | | |
| ANOVA $F_{(4,69)}$=12.3102 **p<0.0001** | | | | | | |
| PLEG4 > ChR2 **IN** AVD Octanol | 76.731 ± 1.866 | 35.869 | **<0.0001** | | | 14 |
| PLEG4 > ChR2 **IN** HAB Octanol | 50.838 ± 3.470 | | | 35.869 | **<0.0001** | 14 |
| PLEG4 > ChR2 **IN PHOTO 2.7K** | 72.561 ± 3.040 | 25.245 | **<0.0001** | 0.930 | **0.3383** | 14 |
| PLEG4 > ChR2 **IN PHOTO 3.5K** | 76.538 ± 3.149 | 35.335 | **<0.0001** | 0.002 | **0.9644** | 14 |
| PLEG4 > ChR2 **IN PHOTO 17K** | 70.380 ± 3.468 | 20.431 | **<0.0001** | 2.157 | **0.1466** | 14 |
| **Figure 5G: PAMG4 > Shi$^{ts}$** (w$^{1118}$ background = (+)) | | | | | | |
| ANOVA $F_{(15,255)}$=15.1556 **p<0.0001** | | | | | | |
| PAMG4 > (+) **IN** AVD Octanol | 82.480 ± 1.542 | 35.775 | **<0.0001** | | | 16 |
| PAMG4 > (+) **IN** HAB Octanol | 61.075 ± 2.954 | | | 35.775 | **<0.0001** | 16 |
| PAMG4 > (+) **IN** DISHAB **shock** | 81.444 ± 2.458 | 32.395 | **<0.0001** | 0.083 | **0.7724** | 16 |
| PAMG4 > (+) **IN** DISHAB **yeast** | 82.105 ± 2.013 | 34.532 | **<0.0001** | 0.011 | **0.9165** | 16 |
| PAMG4 > Shi$^{ts}$ **IN** AVD Octanol | 83.710 ± 2.498 | 29.227 | **<0.0001** | | | 16 |
| PAMG4 > Shi$^{ts}$ **IN** HAB Octanol | 64.363 ± 2.262 | | | 29.227 | **<0.0001** | 16 |
| PAMG4 > Shi$^{ts}$ **IN** DISHAB **shock** | 82.403 ± 2.066 | 25.413 | **<0.0001** | 0.133 | **0.7153** | 16 |
| PAMG4 > Shi$^{ts}$ **IN** DISHAB **yeast** | 64.114 ± 3.113 | 0.004 | **0.9447** | 29.982 | **<0.0001** | 16 |
| PAMG4 > Shi$^{ts}$ UN AVD Octanol | 84.152 ± 1.911 | 51.785 | **<0.0001** | | | 16 |
| PAMG4 > Shi$^{ts}$ UN HAB Octanol | 58.398 ± 3.174 | | | 51.785 | **<0.0001** | 16 |
| PAMG4 > Shi$^{ts}$ UN DISHAB **shock** | 83.594 ± 2.106 | 49.569 | **<0.0001** | 0.024 | **0.8764** | 16 |
| PAMG4 > Shi$^{ts}$ UN DISHAB **yeast** | 78.525 ± 2.702 | 31.629 | **<0.0001** | 2.471 | **0.1172** | 16 |
| Shi$^{ts}$ > (+) **IN** AVD Octanol | 81.581 ± 2.113 | 30.925 | **<0.0001** | | | 16 |
| Shi$^{ts}$ > (+) **IN** HAB Octanol | 61.680 ± 3.533 | | | 30.925 | **<0.0001** | 16 |

| Genotype | Mean ± SEM | HAB Control | | AVD Control | | N |
|---|---|---|---|---|---|---|
| | | F-Ratio | p-value | F-Ratio | p-value | |
| Shi$^{ts}$ > (+) **IN** DISHAB **shock** | 82.153 ± 2.550 | 32.729 | **<0.0001** | 0.025 | **0.8730** | 16 |
| Shi$^{ts}$ > (+) **IN** DISHAB **yeast** | 81.738 ± 2.644 | 31.416 | **<0.0001** | 0.001 | **0.9649** | 16 |
| **Figure 5H: PAMG4 > UAS-ChR2-XXL4 Channel Rhodopsin (ChR2)** | | | | | | |
| ANOVA $F_{(6,69)}=26.3879$ **p<0.0001** | | | | | | |
| PAMG4 > ChR2 **IN** AVD Octanol | 79.031 ± 3.037 | 48.017 | **<0.0001** | | | 10 |
| PAMG4 > ChR2 **IN** HAB Octanol | 46.276 ± 2.243 | | | 48.017 | **<0.0001** | 10 |
| PAMG4 > ChR2 **IN** DISHAB **SHOCK** | 78.576 ± 2.495 | 46.692 | **<0.0001** | 0.009 | **0.9235** | 10 |
| PAMG4 > ChR2 **IN** DISHAB **YEAST** | 70.121 ± 2.975 | 25.447 | **<0.0001** | 3.553 | **0.0640** | 10 |
| PAMG4 > ChR2 **IN DISHAB 2.7K** Photo | 39.230 ± 4.814 | 2.221 | **0.1410** | 70.897 | **<0.0001** | 10 |
| PAMG4 > ChR2 **IN DISHAB 3.5K** Photo | 47.763 ± 4.696 | 0.099 | **0.7540** | 43.755 | **<0.0001** | 10 |
| PAMG4 > ChR2 **IN DISHAB 17K** Photo | 75.048 ± 1.904 | 37.049 | **<0.0001** | 0.710 | **0.4025** | 10 |
| **Figure 5I: PAMG4 > UAS-ChR2-XXL4 Channel Rhodopsin (ChR2) NAIVE** | | | | | | |
| ANOVA $F_{(4,59)}=16.1554$ **p<0.0001** | | | | | | |
| PAMG4 > ChR2 **IN** AVD Octanol | 65.195 ± 2.560 | 44.333 | **<0.0001** | | | 12 |
| PAMG4 > ChR2 **IN** HAB Octanol | 28.751 ± 3.617 | | | 44.333 | **<0.0001** | 12 |
| PAMG4 > ChR2 **IN PHOTO 2.7K** | 61.457 ± 3.985 | 35.704 | **<0.0001** | 0.466 | **0.4974** | 12 |
| PAMG4 > ChR2 **IN PHOTO 3.5K** | 66.388 ± 4.460 | 47.282 | **<0.0001** | 0.047 | **0.8283** | 12 |
| PAMG4 > ChR2 **IN PHOTO 17K** | 57.812 ± 4.412 | 28.189 | **<0.0001** | 1.819 | **0.1828** | 12 |
| **Figure 5J: PPL1 MB504BG4 > Shi$^{ts}$** (w$^{1118}$ background = (+)) | | | | | | |
| ANOVA $F_{(11,191)}=21.4410$ **p<0.0001** | | | | | | |
| PPL1 MB504BG4 > (+) **IN** AVD Octanol | 81.347 ± 1.961 | 51.328 | **<0.0001** | | | 16 |
| PPL1 MB504BG4 > (+) **IN** HAB Octanol | 53.198 ± 3.205 | | | 51.328 | **<0.0001** | 16 |
| PPL1 MB504BG4 > (+) **IN** DISHAB **shock** | 73.878 ± 3.026 | 27.703 | **<0.0001** | 3.613 | **0.0589** | 16 |
| PPL1 MB504BG4 > (+) **IN** DISHAB **yeast** | 75.780 ± 2.738 | 33.033 | **<0.0001** | 2.007 | **0.1582** | 16 |
| PPL1 MB504BG4 > Shi$^{ts}$ **IN** AVD Octanol | 81.153 ± 2.058 | 43.217 | **<0.0001** | | | 16 |
| PPL1 MB504BG4 > Shi$^{ts}$ **IN** HAB Octanol | 55.324 ± 2.468 | | | 43.217 | **<0.0001** | 16 |
| PPL1 MB504BG4 > Shi$^{ts}$ **IN** DISHAB **shock** | 48.882 ± 3.563 | 2.688 | **0.1028** | 67.462 | **<0.0001** | 16 |
| PPL1 MB504BG4 > Shi$^{ts}$ **IN** DISHAB **yeast** | 77.369 ± 3.002 | 31.481 | **<0.0001** | 0.927 | **0.3367** | 16 |
| PPL1 MB504BG4 > Shi$^{ts}$ UN AVD Octanol | 80.913 ± 2.276 | 51.257 | **<0.0001** | | | 16 |
| PPL1 MB504BG4 > Shi$^{ts}$ UN HAB Octanol | 52.784 ± 3.722 | | | 51.257 | **<0.0001** | 16 |
| PPL1 MB504BG4 > Shi$^{ts}$ UN DISHAB **shock** | 77.944 ± 2.044 | 41.009 | **<0.0001** | 0.570 | **0.4508** | 16 |
| PPL1 MB504BG4 > Shi$^{ts}$ UN DISHAB **yeast** | 76.647 ± 2.564 | 36.887 | **<0.0001** | 1.179 | **0.2789** | 16 |
| **Figure 5K: PPL1 MB504BG4 > UAS-ChR2-XXL4 Channel Rhodopsin (ChR2)** | | | | | | |
| ANOVA $F_{(6,111)}=10.9495$ **p<0.0001** | | | | | | |
| PPL1 MB504BG4 > ChR2 **IN** AVD Octanol | 68.482 ± 3.934 | 28.544 | **<0.0001** | | | 16 |
| PPL1 MB504BG4 > ChR2 **IN** HAB Octanol | 39.661 ± 3.943 | | | 28.544 | **<0.0001** | 16 |
| PPL1 MB504BG4 > ChR2 **IN** DISHAB **SHOCK** | 64.333 ± 2.823 | 20.917 | **<0.0001** | 0.591 | **0.4435** | 16 |
| PPL1 MB504BG4 > ChR2 **IN** DISHAB **YEAST** | 61.664 ± 2.631 | 16.637 | **<0.0001** | 1.597 | **0.2090** | 16 |
| PPL1 MB504BG4 > ChR2 **IN DISHAB 2.7K** Photo | 56.158 ± 4.087 | 9.352 | **0.0028** | 5.218 | **0.0243** | 16 |
| PPL1 MB504BG4 > ChR2 **IN DISHAB 3.5K** Photo | 45.460 ± 4.607 | 1.155 | **0.2848** | 18.214 | **<0.0001** | 16 |
| PPL1 MB504BG4 > ChR2 **IN DISHAB 17K** Photo | 36.624 ± 4.240 | 0.317 | **0.5745** | 34.878 | **<0.0001** | 16 |
| **Figure 5L: PPL1 MB504BG4 > UAS-ChR2-XXL4 Channel Rhodopsin (ChR2) NAIVE** | | | | | | |
| ANOVA $F_{(4,79)}=11.0758$ **p<0.0001** | | | | | | |
| PPL1 MB504BG4 > ChR2 **IN** AVD Octanol | 73.927 ± 2.361 | 34.448 | **<0.0001** | | | 16 |
| PPL1 MB504BG4 > ChR2 **IN** HAB Octanol | 41.608 ± 3.754 | | | 34.448 | **<0.0001** | 16 |

| Genotype | Mean ± SEM | HAB Control | | AVD Control | | N |
|---|---|---|---|---|---|---|
| | | F-Ratio | p-value | F-Ratio | p-value | |
| PPL1 MB504BG4 > ChR2 **IN PHOTO 2.7K** | 69.956 ± 4.713 | 26.503 | **<0.0001** | 0.520 | **0.4730** | 16 |
| PPL1 MB504BG4 > ChR2 **IN PHOTO 3.5K** | 68.150 ± 4.388 | 23.234 | **<0.0001** | 1.100 | **0.2974** | 16 |
| PPL1 MB504BG4 > ChR2 **IN PHOTO 17K** | 68.460 ± 3.827 | 23.779 | **<0.0001** | 0.985 | **0.3239** | 16 |
| **Figure 6: PPL1 Dopaminergic Neurons drive intra-modal and cross-modal dishabituation to yeast odor in Drosophila.** | | | | | | |
| **Figure 6A: PPL1 MB504BG4 >** $Shi^{ts}$ ($w^{1118}$ background = (+)) | | | | | | |
| ANOVA $F_{(7,127)}$=23.9017 **p<0.0001** | | | | | | |
| PPL1 MB504BG4 > (+) **IN** AVD yeast | 63.148 ± 3.153 | 70.551 | **<0.0001** | | | 16 |
| PPL1 MB504BG4 > (+) **IN** HAB yeast | 24.791 ± 3.101 | | | 70.551 | **<0.0001** | 16 |
| PPL1 MB504BG4 > (+) **IN** DISHAB **shock** | 58.440 ± 3.334 | 54.296 | **<0.0001** | 1.062 | **0.3046** | 16 |
| PPL1 MB504BG4 > (+) **IN** DISHAB **Octanol** | 59.878 ± 3.218 | 59.036 | **<0.0001** | 0.5126 | **0.4754** | 16 |
| PPL1 MB504BG4 > $Shi^{ts}$ **IN** AVD yeast | 57.773 ± 3.174 | 30.611 | **<0.0001** | | | 16 |
| PPL1 MB504BG4 > $Shi^{ts}$ **IN** HAB yeast | 32.508 ± 2.999 | | | 30.611 | **<0.0001** | 16 |
| PPL1 MB504BG4 > $Shi^{ts}$ **IN** DISHAB **shock** | 32.125 ± 3.574 | 0.007 | **0.9333** | 31.546 | **<0.0001** | 16 |
| PPL1 MB504BG4 > $Shi^{ts}$ **IN** DISHAB **Octanol** | 34.016 ± 3.242 | 0.109 | **0.7417** | 27.066 | **<0.0001** | 16 |
| **Figure 6B: PPL1 MB504BG4 > UAS-ChR2-XXL4 Channel Rhodopsin (ChR2) YEAST HAB** | | | | | | |
| ANOVA $F_{(5,83)}$=17.0167 **p<0.0001** | | | | | | |
| PPL1 MB504BG4 > ChR2 **IN** AVD Yeast | 53.826 ± 3.397 | 36.417 | **<0.0001** | | | 14 |
| PPL1 MB504BG4 > ChR2 **IN** HAB Yeast | 22.165 ± 3.992 | | | 36.417 | **<0.0001** | 14 |
| PPL1 MB504BG4 > ChR2 **IN** DISHAB **OCTANOL** | 57.941 ± 3.802 | 46.498 | **<0.0001** | 0.615 | **0.4352** | 14 |
| PPL1 MB504BG4 > ChR2 **IN DISHAB 2.7K** Photo | 37.909 ± 3.656 | 9.004 | **0.0036** | 9.204 | **0.0032** | 14 |
| PPL1 MB504BG4 > ChR2 **IN DISHAB 3.5K** Photo | 31.035 ± 3.199 | 2.858 | **0.0948** | 18.869 | **<0.0001** | 14 |
| PPL1 MB504BG4 > ChR2 **IN DISHAB 17K** Photo | 23.021 ± 4.126 | 0.026 | **0.8708** | 34.475 | **<0.0001** | 14 |
| **Figure 6C: PAMG4 >** $Shi^{ts}$ ($w^{1118}$ background = (+)) | | | | | | |
| ANOVA $F_{(7,95)}$=16.1573 **p<0.0001** | | | | | | |
| PAMG4 > (+) **IN** AVD yeast | 60.256 ± 4.162 | 40.400 | **<0.0001** | | | 12 |
| PAMG4 > (+) **IN** HAB yeast | 34.119 ± 2.976 | | | 40.400 | **<0.0001** | 12 |
| PAMG4 > (+) **IN** DISHAB **shock** | 60.413 ± 2.777 | 40.889 | **<0.0001** | 0.001 | **0.9694** | 12 |
| PAMG4 > (+) **IN** DISHAB **Octanol** | 56.914 ± 2.249 | 30.729 | **<0.0001** | 0.660 | **0.4185** | 12 |
| PAMG4 > $Shi^{ts}$ **IN** AVD yeast | 59.251 ± 2.778 | 42.268 | **<0.0001** | | | 12 |
| PAMG4 > $Shi^{ts}$ **IN** HAB yeast | 32.517 ± 2.695 | | | 42.268 | **<0.0001** | 12 |
| PAMG4 > $Shi^{ts}$ **IN** DISHAB **shock** | 57.337 ± 2.588 | 36.434 | **<0.0001** | 0.216 | **0.6428** | 12 |
| PAMG4 > $Shi^{ts}$ **IN** DISHAB **Octanol** | 55.570 ± 2.642 | 31.431 | **<0.0001** | 0.801 | **0.3731** | 12 |
| **Figure 7: The role of PPL1 neuronal subsets in cross modal dishabituation.** | | | | | | |
| **Figure 7B: PPL1-γ2a′1 MB296BG4 > UAS-ChR2-XXL4 Channel Rhodopsin (ChR2)** | | | | | | |
| ANOVA $F_{(6,125)}$=10.0663 **p<0.0001** | | | | | | |
| MB296BG4 > ChR2 **IN** AVD Octanol | 70.854 ± 2.452 | 26.033 | **<0.0001** | | | 18 |
| MB296BG4 > ChR2 **IN** HAB Octanol | 51.826 ± 2.617 | | | 26.033 | **<0.0001** | 18 |
| MB296BG4 > ChR2 **IN** DISHAB **SHOCK** | 71.333 ± 2.160 | 27.359 | **<0.0001** | 0.016 | **0.8981** | 18 |
| MB296BG4 > ChR2 **IN** DISHAB **YEAST** | 73.330 ± 2.700 | 33.249 | **<0.0001** | 0.440 | **0.5080** | 18 |
| MB296BG4 > ChR2 **IN DISHAB 2.7K** Photo | 65.337 ± 2.736 | 13.125 | **<0.0005** | 2.188 | **0.1416** | 18 |
| MB296BG4 > ChR2 **IN DISHAB 3.5K** Photo | 61.762 ± 2.699 | 7.098 | **0.0087** | 5.943 | **0.0162** | 18 |
| MB296BG4 > ChR2 **IN DISHAB 17K** Photo | 55.192 ± 3.011 | 0.814 | **0.3685** | 17.637 | **<0.0001** | 18 |
| **Figure 7C: PPL1-γ2a′1 MB296BG4 > UAS-ChR2-XXL4 Channel Rhodopsin (ChR2) NAIVE** | | | | | | |
| ANOVA $F_{(4,84)}$=11.5001 **p<0.0001** | | | | | | |
| MB296BG4 > ChR2 **IN** AVD Octanol | 70.739 ± 2.519 | 30.226 | **<0.0001** | | | 17 |

| Genotype | Mean ± SEM | HAB Control | | AVD Control | | N |
|---|---|---|---|---|---|---|
| | | F-Ratio | p-value | F-Ratio | p-value | |
| MB296BG4 > ChR2 **IN** HAB Octanol | 41.322 ± 4.061 | | | 30.226 | **<0.0001** | 17 |
| MB296BG4 > ChR2 **IN PHOTO 2.7K** | 63.545 ± 4.262 | 17.250 | **<0.0001** | 1.807 | **0.1825** | 17 |
| MB296BG4 > ChR2 **IN PHOTO 3.5K** | 55.781 ± 3.505 | 7.301 | **0.0084** | 7.815 | **0.0064** | 17 |
| MB296BG4 > ChR2 **IN PHOTO 17K** | 42.744 ± 4.274 | 0.070 | **0.7910** | 27.374 | **<0.0001** | 17 |
| **Figure 7E: PPL1-γ1pedc MB320CG4 > UAS-ChR2-XXL4 Channel Rhodopsin (ChR2)** | | | | | | |
| ANOVA $F_{(6,111)}$=22.2699 **p<0.0001** | | | | | | |
| MB320CG4 > ChR2 **IN** AVD Octanol | 65.994 ± 2.863 | 53.546 | **<0.0001** | | | 16 |
| MB320CG4 > ChR2 **IN** HAB Octanol | 37.046 ± 2.949 | | | 53.546 | **<0.0001** | 16 |
| MB320CG4 > ChR2 **IN** DISHAB **SHOCK** | 70.528 ± 2.174 | 71.630 | **<0.0001** | 1.313 | **0.2544** | 16 |
| MB320CG4 > ChR2 **IN** DISHAB **YEAST** | 68.693 ± 2.870 | 63.996 | **<0.0001** | 0.465 | **0.4965** | 16 |
| MB320CG4 > ChR2 **IN DISHAB 2.7K** Photo | 52.853 ± 2.259 | 15.965 | **<0.0002** | 11.034 | **0.0012** | 16 |
| MB320CG4 > ChR2 **IN DISHAB 3.5K** Photo | 52.286 ± 3.265 | 14.840 | **<0.0003** | 12.007 | **<0.0008** | 16 |
| MB320CG4 > ChR2 **IN DISHAB 17K** Photo | 42.165 ± 3.022 | 1.674 | **0.1985** | 36.283 | **<0.0001** | 16 |
| **Figure 7F: PPL1-γ1pedc MB320CG4 > UAS-ChR2-XXL4 Channel Rhodopsin (ChR2) NAIVE** | | | | | | |
| ANOVA $F_{(4,59)}$=14.0823 **p<0.0001** | | | | | | |
| MB320CG4 > ChR2 **IN** AVD Octanol | 70.811 ± 2.389 | 36.423 | **<0.0001** | | | 12 |
| MB320CG4 > ChR2 **IN** HAB Octanol | 38.993 ± 3.539 | | | 36.423 | **<0.0001** | 12 |
| MB320CG4 > ChR2 **IN PHOTO 2.7K** | 68.587 ± 4.618 | 31.509 | **<0.0001** | 0.177 | **0.6747** | 12 |
| MB320CG4 > ChR2 **IN PHOTO 3.5K** | 50.823 ± 4.086 | 5.035 | **0.0288** | 14.373 | **<0.0004** | 12 |
| MB320CG4 > ChR2 **IN PHOTO 17K** | 46.361 ± 3.636 | 1.953 | **0.1678** | 21.507 | **<0.0001** | 12 |
| **Figure 7H: PPL1-α2α′2 MB058BG4 > UAS-ChR2-XXL4 Channel Rhodopsin (ChR2)** | | | | | | |
| ANOVA $F_{(6,97)}$=21.9961 **p<0.0001** | | | | | | |
| MB058BG4 > ChR2 **IN** AVD Octanol | 65.612 ± 3.428 | 37.196 | **<0.0001** | | | 14 |
| MB058BG4 > ChR2 **IN** HAB Octanol | 36.876 ± 3.867 | | | 37.196 | **<0.0001** | 14 |
| MB058BG4 > ChR2 **IN** DISHAB **SHOCK** | 69.155 ± 2.466 | 46.936 | **<0.0001** | 0.565 | **0.4539** | 14 |
| MB058BG4 > ChR2 **IN** DISHAB **YEAST** | 71.403 ± 2.162 | 53.699 | **<0.0001** | 1.510 | **0.2221** | 14 |
| MB058BG4 > ChR2 **IN DISHAB 2.7K** Photo | 38.018 ± 4.060 | 0.058 | **0.8090** | 34.299 | **<0.0001** | 14 |
| MB058BG4 > ChR2 **IN DISHAB 3.5K** Photo | 38.858 ± 3.841 | 0.176 | **0.6750** | 32.243 | **<0.0001** | 14 |
| MB058BG4 > ChR2 **IN DISHAB 17K** Photo | 48.214 ± 2.996 | 5.791 | **0.0181** | 13.633 | **<0.0004** | 14 |
| **Figure 7I: PPL1-α2α′2 MB058BG4 > UAS-ChR2-XXL4 Channel Rhodopsin (ChR2) NAIVE** | | | | | | |
| ANOVA $F_{(4,59)}$=16.7817 **p<0.0001** | | | | | | |
| MB058BG4 > ChR2 **IN** AVD Octanol | 63.227 ± 3.533 | 37.852 | **<0.0001** | | | 12 |
| MB058BG4 > ChR2 **IN** HAB Octanol | 31.211 ± 3.480 | | | 37.852 | **<0.0001** | 12 |
| MB058BG4 > ChR2 **IN PHOTO 2.7K** | 63.306 ± 4.579 | 38.039 | **<0.0001** | <0.001 | **0.9879** | 12 |
| MB058BG4 > ChR2 **IN PHOTO 3.5K** | 63.152 ± 3.650 | 37.674 | **<0.0001** | <0.001 | **0.9885** | 12 |
| MB058BG4 > ChR2 **IN PHOTO 17K** | 68.454 ± 2.966 | 51.222 | **<0.0001** | 1.009 | **0.3195** | 12 |
| **Figure 7K: PPL1-a3 MB630BG4 > UAS-ChR2-XXL4 Channel Rhodopsin (ChR2)** | | | | | | |
| ANOVA $F_{(6,111)}$=9.1773 **p<0.0001** | | | | | | |
| MB630BG4 > ChR2 **IN** AVD Octanol | 70.023 ± 3.162 | 15.507 | **<0.0002** | | | 16 |
| MB630BG4 > ChR2 **IN** HAB Octanol | 52.907 ± 3.003 | | | 15.507 | **<0.0002** | 16 |
| MB630BG4 > ChR2 **IN** DISHAB **SHOCK** | 71.233 ± 1.749 | 17.777 | **<0.0001** | 0.077 | **0.7812** | 16 |
| MB630BG4 > ChR2 **IN** DISHAB **YEAST** | 72.633 ± 2.203 | 20.597 | **<0.0001** | 0.360 | **0.5494** | 16 |
| MB630BG4 > ChR2 **IN DISHAB 2.7K** Photo | 53.043 ± 3.411 | 0.001 | **0.9751** | 15.262 | **<0.0002** | 16 |
| MB630BG4 > ChR2 **IN DISHAB 3.5K** Photo | 55.273 ± 3.726 | 0.296 | **0.5873** | 11.516 | **<0.0010** | 16 |
| MB630BG4 > ChR2 **IN DISHAB 17K** Photo | 54.813 ± 3.696 | 0.192 | **0.6618** | 12.245 | **<0.0007** | 16 |

| Genotype | Mean ± SEM | HAB Control | | AVD Control | | N |
|---|---|---|---|---|---|---|
| | | F-Ratio | p-value | F-Ratio | p-value | |
| **Figure 7L: PPL1-a3 MB630BG4 > UAS-ChR2-XXL4 Channel Rhodopsin (ChR2) NAIVE** | | | | | | |
| ANOVA $F_{(4,69)}$=10.8266 **p<0.0001** | | | | | | |
| MB630BG4 > ChR2 **IN** AVD Octanol | 67.071 ± 2.996 | 37.977 | **<0.0001** | | | 14 |
| MB630BG4 > ChR2 **IN** HAB Octanol | 39.840 ± 3.269 | | | 37.977 | **<0.0001** | 14 |
| MB630BG4 > ChR2 **IN PHOTO 2.7K** | 61.982 ± 2.820 | 25.108 | **<0.0001** | 1.326 | **0.2536** | 14 |
| MB630BG4 > ChR2 **IN PHOTO 3.5K** | 58.443 ± 3.605 | 17.722 | **<0.0001** | 3.813 | **0.0551** | 14 |
| MB630BG4 > ChR2 **IN PHOTO 17K** | 57.211 ± 2.861 | 15.452 | **<0.0003** | 4.979 | **0.0290** | 14 |
| **Figure 8: APL neurons regulate Homosensory Dishabituation.** | | | | | | |
| **Figure 8B: APLG4 > Shi$^{ts}$** ($w^{1118}$ background = (+)) | | | | | | |
| ANOVA $F_{(11,143)}$=19.1816 **p<0.0001** | | | | | | |
| APLG4 > (+) **IN** AVD Octanol | 78.797 ± 2.804 | 71.397 | **<0.0001** | | | 12 |
| APLG4 > (+) **IN** HAB Octanol | 33.802 ± 4.533 | | | 71.397 | **<0.0001** | 12 |
| APLG4 > (+) **IN** DISHAB **shock** | 63.433 ± 4.859 | 30.963 | **<0.0001** | 8.324 | **0.0045** | 12 |
| APLG4 > (+) **IN** DISHAB **yeast** | 64.711 ± 3.554 | 33.691 | **<0.0001** | 6.997 | **0.0091** | 12 |
| APLG4 > Shi$^{ts}$ **IN** AVD Octanol | 71.605 ± 2.545 | 39.846 | **<0.0001** | | | 12 |
| APLG4 > Shi$^{ts}$ **IN** HAB Octanol | 37.991 ± 3.996 | | | 39.846 | **<0.0001** | 12 |
| APLG4 > Shi$^{ts}$ **IN** DISHAB **shock** | 68.377 ± 5.176 | 32.560 | **<0.0001** | 0.367 | **0.5454** | 12 |
| APLG4 > Shi$^{ts}$ **IN** DISHAB **yeast** | 43.811 ± 4.564 | 1.194 | **0.2764** | 27.243 | **<0.0001** | 12 |
| APLG4 > Shi$^{ts}$ UN AVD Octanol | 81.563 ± 2.116 | 39.565 | **<0.0001** | | | 12 |
| APLG4 > Shi$^{ts}$ UN HAB Octanol | 48.068 ± 3.753 | | | 39.565 | **<0.0001** | 12 |
| APLG4 > Shi$^{ts}$ UN DISHAB **shock** | 77.949 ± 2.419 | 31.487 | **<0.0001** | 0.460 | **0.4984** | 12 |
| APLG4 > Shi$^{ts}$ UN DISHAB **yeast** | 68.029 ± 3.312 | 14.050 | **<0.0003** | 6.460 | **0.0121** | 12 |
| **Figure 8C: APLG4 > UAS-ChR2-XXL4 Channel Rhodopsin (ChR2)** | | | | | | |
| ANOVA $F_{(6,97)}$=11.3064 **p<0.0001** | | | | | | |
| APLG4 > ChR2 **IN** AVD Octanol | 81.503 ± 2.670 | 42.634 | **<0.0001** | | | 14 |
| APLG4 > ChR2 **IN** HAB Octanol | 48.726 ± 3.414 | | | 42.634 | **<0.0001** | 14 |
| APLG4 > ChR2 **IN** DISHAB **SHOCK** | 77.274 ± 2.897 | 32.343 | **<0.0001** | 0.709 | **0.4017** | 14 |
| APLG4 > ChR2 **IN** DISHAB **YEAST** | 67.466 ± 3.487 | 13.936 | **<0.0004** | 7.819 | **0.0063** | 14 |
| APLG4 > ChR2 **IN DISHAB 2.7K** Photo | 56.155 ± 3.910 | 2.189 | **0.1423** | 25.499 | **<0.0001** | 14 |
| APLG4 > ChR2 **IN DISHAB 3.5K** Photo | 58.350 ± 4.624 | 3.675 | **0.0583** | 21.273 | **<0.0001** | 14 |
| APLG4 > ChR2 **IN DISHAB 17K** Photo | 71.602 ± 3.488 | 20.767 | **<0.0001** | 3.890 | **0.0516** | 14 |
| **Figure 8D: APLG4 > UAS-ChR2-XXL4 Channel Rhodopsin (ChR2) NAIVE** | | | | | | |
| ANOVA $F_{(4,69)}$=16.5584 **p<0.0001** | | | | | | |
| APLG4 > ChR2 **IN** AVD Octanol | 71.085 ± 3.518 | 51.575 | **<0.0001** | | | 14 |
| APLG4 > ChR2 **IN** HAB Octanol | 34.713 ± 2.934 | | | 51.575 | **<0.0001** | 14 |
| APLG4 > ChR2 **IN PHOTO 2.7K** | 62.775 ± 3.651 | 30.700 | **<0.0001** | 2.692 | **0.1056** | 14 |
| APLG4 > ChR2 **IN PHOTO 3.5K** | 68.993 ± 4.127 | 45.813 | **<0.0001** | 0.170 | **0.6809** | 14 |
| APLG4 > ChR2 **IN PHOTO 17K** | 59.131 ± 3.572 | 23.245 | **<0.0001** | 5.570 | **0.0212** | 14 |
| **Figure 8E: APLG4 > Shi$^{ts}$** ($w^{1118}$ background = (+)) | | | | | | |
| ANOVA $F_{(7,127)}$=11.8296 **p<0.0001** | | | | | | |
| APLG4 > (+) **IN** AVD yeast | 58.374 ± 4.089 | 23.676 | **<0.0001** | | | 16 |
| APLG4 > (+) **IN** HAB yeast | 28.834 ± 4.052 | | | 23.676 | **<0.0001** | 16 |
| APLG4 > (+) **IN** DISHAB **shock** | 54.735 ± 5.048 | 18.202 | **<0.0001** | 0.359 | **0.5500** | 16 |
| APLG4 > (+) **IN** DISHAB **Octanol** | 56.601 ± 4.579 | 20.918 | **<0.0001** | 0.085 | **0.7707** | 16 |
| APLG4 > Shi$^{ts}$ **IN** AVD yeast | 56.982 ± 3.778 | 24.074 | **<0.0001** | | | 16 |

| Genotype | Mean ± SEM | HAB Control | | AVD Control | | N |
|---|---|---|---|---|---|---|
| | | F-Ratio | p-value | F-Ratio | p-value | |
| APLG4 > Shi$^{ts}$ **IN** HAB yeast | 27.193 ± 4.033 | | | 24.074 | **<0.0001** | 16 |
| APLG4 > Shi$^{ts}$ **IN** DISHAB **shock** | 51.483 ± 4.030 | 16.006 | **<0.0002** | 0.820 | **0.3668** | 16 |
| APLG4 > Shi$^{ts}$ **IN** DISHAB **Octanol** | 26.189 ± 4.584 | 0.027 | **0.8689** | 25.724 | **<0.0001** | 16 |
| **Figure 8F: G80$^{ts}$ ; APLG4 > GAD-RNAi** (w$^{1118}$ background = (+)) VDRC 32344 | | | | | | |
| ANOVA $F_{(15,223)}$=21.4876 **p<0.0001** | | | | | | |
| G80$^{ts}$ ; APLG4 > (+) **IN** AVD Octanol | 78.688 ± 2.492 | 47.606 | **<0.0001** | | | 14 |
| G80$^{ts}$ ; APLG4 > (+) **IN** HAB Octanol | 50.683 ± 3.222 | | | 47.606 | **<0.0001** | 14 |
| G80$^{ts}$ ; APLG4 > (+) **IN** DISHAB **shock** | 77.160 ± 2.132 | 42.555 | **<0.0001** | 0.141 | **0.7070** | 14 |
| G80$^{ts}$ ; APLG4 > (+) **IN** DISHAB **yeast** | 75.060 ± 2.862 | 36.071 | **<0.0001** | 0.798 | **0.3724** | 14 |
| G80$^{ts}$ ; APLG4 > GAD-RNAi **IN** AVD Octanol | 82.617 ± 2.403 | 41.591 | **<0.0001** | | | 14 |
| G80$^{ts}$ ; APLG4 > GAD-RNAi **IN** HAB Octanol | 56.441 ± 2.389 | | | 41.591 | **<0.0001** | 14 |
| G80$^{ts}$ ; APLG4 > GAD-RNAi **IN** DISHAB **shock** | 73.239 ± 2.830 | 17.126 | **<0.0001** | 5.339 | **0.0218** | 14 |
| G80$^{ts}$ ; APLG4 > GAD-RNAi **IN** DISHAB **yeast** | 57.365 ± 2.782 | 0.051 | **0.8202** | 38.709 | **<0.0001** | 14 |
| G80$^{ts}$ ; APLG4 > GAD-RNAi UN AVD Octanol | 88.540 ± 2.219 | 45.157 | **<0.0001** | | | 14 |
| G80$^{ts}$ ; APLG4 > GAD-RNAi UN HAB Octanol | 61.264 ± 3.052 | | | 45.157 | **<0.0001** | 14 |
| G80$^{ts}$ ; APLG4 > GAD-RNAi UN DISHAB **shock** | 83.130 ± 1.798 | 29.022 | **<0.0001** | 1.776 | **0.1840** | 14 |
| G80$^{ts}$ ; APLG4 > GAD-RNAi UN DISHAB **yeast** | 78.562 ± 3.443 | 18.161 | **<0.0001** | 6.043 | **0.0147** | 14 |
| GAD-RNAI **IN** > (+) AVD Octanol | 73.233 ± 3.155 | 67.842 | **<0.0001** | | | 14 |
| GAD-RNAI **IN** > (+) HAB Octanol | 39.802 ± 3.618 | | | 67.842 | **<0.0001** | 14 |
| GAD-RNAI **IN** > (+) DISHAB **shock** | 66.345 ± 3.544 | 42.768 | **<0.0001** | 2.879 | **0.0912** | 14 |
| GAD-RNAI **IN** > (+) DISHAB **yeast** | 64.987 ± 3.204 | 38.504 | **<0.0001** | 4.126 | **0.0434** | 14 |
| **Figure 8G: G80$^{ts}$ ; APLG4 > TBH-RNAi A** (y;v background = (+)) BL 27667 | | | | | | |
| ANOVA $F_{(11,143)}$=21.4635 **p<0.0001** | | | | | | |
| G80$^{ts}$ ; APLG4 > (+) **IN** AVD Octanol | 68.310 ± 3.072 | 56.192 | **<0.0001** | | | 12 |
| G80$^{ts}$ ; APLG4 > (+) **IN** HAB Octanol | 35.396 ± 3.299 | | | 56.192 | **<0.0001** | 12 |
| G80$^{ts}$ ; APLG4 > (+) **IN** DISHAB **shock** | 64.047 ± 3.266 | 42.579 | **<0.0001** | 0.942 | **0.3333** | 12 |
| G80$^{ts}$ ; APLG4 > (+) **IN** DISHAB **yeast** | 59.235 ± 2.658 | 29.477 | **<0.0001** | 4.271 | **0.0407** | 12 |
| G80$^{ts}$ ; APLG4 > TBH-RNAi **IN** AVD Octanol | 73.537 ± 3.308 | 50.226 | **<0.0001** | | | 12 |
| G80$^{ts}$ ; APLG4 > TBH-RNAi **IN** HAB Octanol | 42.419 ± 3.954 | | | 50.226 | **<0.0001** | 12 |
| G80$^{ts}$ ; APLG4 > TBH-RNAi **IN** DISHAB **shock** | 64.516 ± 3.556 | 25.326 | **<0.0001** | 4.221 | **0.0418** | 12 |
| G80$^{ts}$ ; APLG4 > TBH-RNAi **IN** DISHAB **yeast** | 61.440 ± 3.301 | 18.765 | **<0.0001** | 7.590 | **0.0066** | 12 |
| G80$^{ts}$ ; APLG4 > TBH-RNAi UN AVD Octanol | 84.465 ± 2.666 | 26.780 | **<0.0001** | | | 12 |
| G80$^{ts}$ ; APLG4 > TBH-RNAi UN HAB Octanol | 61.743 ± 3.088 | | | 26.780 | **<0.0001** | 12 |
| G80$^{ts}$ ; APLG4 > TBH-RNAi UN DISHAB **shock** | 78.871 ± 2.223 | 15.216 | **<0.0002** | 1.623 | **0.2048** | 12 |
| G80$^{ts}$ ; APLG4 > TBH-RNAi UN DISHAB **yeast** | 78.474 ± 2.424 | 14.520 | **<0.0003** | 1.861 | **0.1747** | 12 |
| **Figure 9: MBONs 11,16,17 regulate olfactory Habituation** | | | | | | |
| **Figure 9B: MBON11 MB112CG4 > Shi$^{ts}$** (w$^{1118}$ background = (+)) | | | | | | |
| ANOVA $F_{(11,191)}$=20.5081 **p<0.0001** | | | | | | |
| **MBON11** MB112CG4 > (+) **IN** AVD Octanol | 75.123 ± 2.771 | 73.848 | **<0.0001** | | | 16 |
| **MBON11** MB112CG4 > (+) **IN** HAB Octanol | 41.195 ± 3.659 | | | 73.848 | **<0.0001** | 16 |
| **MBON11** MB112CG4 > (+) **IN** DISHAB **shock** | 68.593 ± 3.234 | 48.159 | **<0.0001** | 2.735 | **0.0999** | 16 |
| **MBON11** MB112CG4 > (+) **IN** DISHAB **yeast** | 66.798 ± 2.801 | 42.053 | **<0.0001** | 4.446 | **0.0363** | 16 |
| **MBON11** MB112CG4 > Shi$^{ts}$ **IN** AVD Octanol | 71.733 ± 2.466 | 1.349 | **0.2468** | | | 16 |
| **MBON11** MB112CG4 > Shi$^{ts}$ **IN** HAB Octanol | 67.147 ± 2.968 | | | 1.349 | **0.2468** | 16 |
| **MBON11** MB112CG4 > Shi$^{ts}$ **IN** DISHAB **shock** | 66.654 ± 2.682 | 0.015 | **0.9007** | 1.655 | **0.1998** | 16 |
| **MBON11** MB112CG4 > Shi$^{ts}$ **IN** DISHAB **yeast** | 67.352 ± 2.086 | 0.002 | **0.9585** | 1.231 | **0.2686** | 16 |
| **MBON11** MB112CG4 > Shi$^{ts}$ UN AVD Octanol | 73.669 ± 2.160 | 97.548 | **<0.0001** | | | 16 |

| Genotype | Mean ± SEM | HAB Control | | AVD Control | | N |
|---|---|---|---|---|---|---|
| | | F-Ratio | p-value | F-Ratio | p-value | |
| **MBON11** MB112CG4 > $Shi^{ts}$ UN HAB Octanol | 34.676 ± 3.255 | | | 97.548 | **<0.0001** | 16 |
| **MBON11** MB112CG4 > $Shi^{ts}$ UN DISHAB **shock** | 69.764 ± 2.085 | 78.986 | **<0.0001** | 0.978 | **0.3238** | 16 |
| **MBON11** MB112CG4 > $Shi^{ts}$ UN DISHAB **yeast** | 67.340 ± 2.841 | 68.453 | **<0.0001** | 2.569 | **0.1106** | 16 |
| **Figure 9C: MBON11 MB112CG4 > UAS-ChR2-XXL4 Channel Rhodopsin (ChR2)** | | | | | | |
| ANOVA $F_{(6,139)}=23.5253$ **p<0.0001** | | | | | | |
| **MBON11** MB112CG4 > ChR2 **IN** AVD Octanol | 66.610 ± 3.088 | 37.306 | **<0.0001** | | | 20 |
| **MBON11** MB112CG4 > ChR2 **IN** HAB Octanol | 39.005 ± 3.311 | | | 37.306 | **<0.0001** | 20 |
| **MBON11** MB112CG4 > ChR2 **IN** DISHAB **SHOCK** | 67.263 ± 2.804 | 39.094 | **<0.0001** | 0.020 | **0.8852** | 20 |
| **MBON11** MB112CG4 > ChR2 **IN** DISHAB **YEAST** | 64.507 ± 2.804 | 31.840 | **<0.0001** | 0.216 | **0.6425** | 20 |
| **MBON11** MB112CG4 > ChR2 **IN DISHAB 2.7K** Photo | 40.000 ± 4.102 | 0.048 | **0.8260** | 34.665 | **<0.0001** | 20 |
| **MBON11** MB112CG4 > ChR2 **IN DISHAB 3.5K** Photo | 32.940 ± 2.935 | 1.800 | **0.1819** | 55.499 | **<0.0001** | 20 |
| **MBON11** MB112CG4 > ChR2 **IN DISHAB 17K** Photo | 37.929 ± 3.132 | 0.056 | **0.8121** | 40.271 | **<0.0001** | 20 |
| **Figure 9D: MBON11 MB112CG4 > UAS-ChR2-XXL4 Channel Rhodopsin (ChR2) NAIVE** | | | | | | |
| ANOVA $F_{(4,99)}=13.9121$ **p<0.0001** | | | | | | |
| **MBON11** MB112CG4 > ChR2 **IN** AVD Octanol | 66.610 ± 3.088 | 32.593 | **<0.0001** | | | 20 |
| **MBON11** MB112CG4 > ChR2 **IN** HAB Octanol | 39.005 ± 3.311 | | | 32.593 | **<0.0001** | 20 |
| **MBON11** MB112CG4 > ChR2 **IN PHOTO 2.7K** | 44.848 ± 3.698 | 1.460 | **0.2298** | 20.254 | **<0.0001** | 20 |
| **MBON11** MB112CG4 > ChR2 **IN PHOTO 3.5K** | 39.858 ± 3.022 | 0.031 | **0.8602** | 30.609 | **<0.0001** | 20 |
| **MBON11** MB112CG4 > ChR2 **IN PHOTO 17K** | 33.989 ± 3.890 | 1.076 | **0.3022** | 45.513 | **<0.0001** | 20 |
| **Figure 9F: MBON16&17 MB027BG4 > $Shi^{ts}$** ($w^{1118}$ background = (+)) | | | | | | |
| ANOVA $F_{(11,167)}=25.0176$ **p<0.0001** | | | | | | |
| **MBON16&17** MB027BG4 > (+) **IN** AVD Octanol | 73.048 ± 1.924 | 109.587 | **<0.0001** | | | 14 |
| **MBON16&17** MB027BG4 > (+) **IN** HAB Octanol | 39.301 ± 2.790 | | | 109.587 | **<0.0001** | 14 |
| **MBON16&17** MB027BG4 > (+) **IN** DISHAB **shock** | 71.731 ± 2.005 | 101.448 | **<0.0001** | 0.167 | **0.6830** | 14 |
| **MBON16&17** MB027BG4 > (+) **IN** DISHAB **yeast** | 69.205 ± 2.120 | 86.260 | **<0.0001** | 1.424 | **0.2344** | 14 |
| **MBON16&17** MB027BG4 > $Shi^{ts}$ **IN** AVD Octanol | 69.415 ± 1.948 | 0.089 | **0.7657** | | | 14 |
| **MBON16&17** MB027BG4 > $Shi^{ts}$ **IN** HAB Octanol | 68.454 ± 2.602 | | | 0.089 | **0.7657** | 14 |
| **MBON16&17** MB027BG4 > $Shi^{ts}$ **IN** DISHAB **shock** | 67.573 ± 2.608 | 0.074 | **0.7846** | 0.327 | **0.5680** | 14 |
| **MBON16&17** MB027BG4 > $Shi^{ts}$ **IN** DISHAB **yeast** | 65.679 ± 2.023 | 0.742 | **0.3900** | 1.346 | **0.2476** | 14 |
| **MBON16&17** MB027BG4 > $Shi^{ts}$ UN AVD Octanol | 70.005 ± 2.611 | 80.194 | **<0.0001** | | | 14 |
| **MBON14&18** MB027BG4 > $Shi^{ts}$ UN HAB Octanol | 41.171 ± 2.401 | | | 80.194 | **<0.0001** | 14 |
| **MBON16&17** MB027BG4 > $Shi^{ts}$ UN DISHAB **shock** | 68.378 ± 1.939 | 71.400 | **<0.0001** | 0.255 | **0.6140** | 14 |
| **MBON16&17** MB027BG4 > $Shi^{ts}$ UN DISHAB **yeast** | 66.672 ± 2.093 | 62.726 | **<0.0001** | 1.071 | **0.3021** | 14 |
| **Figure 9G: MBON16&17 MB027BG4 > UAS-ChR2-XXL4 Channel Rhodopsin (ChR2)** | | | | | | |
| ANOVA $F_{(6,97)}=29.6637$ **p<0.0001** | | | | | | |
| **MBON16&17** MB027BG4 > ChR2 **IN** AVD Octanol | 74.963 ± 2.450 | 63.587 | **<0.0001** | | | 14 |
| **MBON16&17** MB027BG4 > ChR2 **IN** HAB Octanol | 44.817 ± 3.001 | | | 63.587 | **<0.0001** | 14 |
| **MBON16&17** MB027BG4 > ChR2 **IN** DISHAB **SHOCK** | 71.301 ± 2.559 | 49.077 | **<0.0001** | 0.938 | **0.3353** | 14 |
| **MBON16&17** MB027BG4 > ChR2 **IN** DISHAB **YEAST** | 69.734 ± 2.500 | 43.441 | **<0.0001** | 1.913 | **0.1700** | 14 |
| **MBON16&17** MB027BG4 > ChR2 **IN DISHAB 2.7K** Photo | 50.531 ± 2.432 | 2.284 | **0.1341** | 41.766 | **<0.0001** | 14 |

| Genotype | Mean ± SEM | HAB Control | | AVD Control | | N |
|---|---|---|---|---|---|---|
| | | F-Ratio | p-value | F-Ratio | p-value | |
| **MBON16&17** MB027BG4 > ChR2 **IN DISHAB 3.5K** Photo | 44.374 ± 3.220 | 0.013 | **0.9071** | 65.466 | **<0.0001** | 14 |
| **MBON16&17** MB027BG4 > ChR2 **IN DISHAB 17K** Photo | 41.729 ± 2.432 | 0.666 | **0.4163** | 77.276 | **<0.0001** | 14 |
| **Figure 9H: MBON16&17 MB027BG4 > UAS-ChR2-XXL4 Channel Rhodopsin (ChR2) NAIVE** | | | | | | |
| ANOVA $F_{(4,69)}$=23.6692 **p<0.0001** | | | | | | |
| **MBON16&17** MB027BG4 > ChR2 **IN** AVD Octanol | 74.963 ± 2.450 | 53.343 | **<0.0001** | | | 14 |
| **MBON16&17** MB027BG4 > ChR2 **IN** HAB Octanol | 44.817 ± 3.001 | | | 53.343 | **<0.0001** | 14 |
| **MBON16&17** MB027BG4 > ChR2 **IN PHOTO 2.7K** | 46.225 ± 2.988 | 0.116 | **0.7340** | 48.475 | **<0.0001** | 14 |
| **MBON16&17** MB027BG4 > ChR2 **IN PHOTO 3.5K** | 42.051 ± 3.012 | 0.449 | **0.5051** | 63.581 | **<0.0001** | 14 |
| **MBON16&17** MB027BG4 > ChR2 **IN PHOTO 17K** | 41.102 ± 3.093 | 0.809 | **0.3714** | 67.299 | **<0.0001** | 14 |
| **Figure 10: MBONs 8,9,18 regulate olfactory Dishabituation** | | | | | | |
| **Figure 10B: MBON15&18 MB050BG4 > UAS-ChR2-XXL4 Channel Rhodopsin (ChR2)** | | | | | | |
| ANOVA $F_{(6,97)}$=18.2249 **p<0.0001** | | | | | | |
| **MBON15&18** MB050BG4 > ChR2 **IN** AVD Octanol | 70.276 ± 2.702 | 54.903 | **<0.0001** | | | 14 |
| **MBON15&18** MB050BG4 > ChR2 **IN** HAB Octanol | 37.798 ± 3.101 | | | 54.903 | **<0.0001** | 14 |
| **MBON15&18** MB050BG4 > ChR2 **IN** DISHAB **SHOCK** | 69.716 ± 2.770 | 53.026 | **<0.0001** | 0.016 | **0.8985** | 14 |
| **MBON15&18** MB050BG4 > ChR2 **IN** DISHAB **YEAST** | 65.527 ± 2.618 | 40.021 | **<0.0001** | 1.173 | **0.2814** | 14 |
| **MBON15&18** MB050BG4 > ChR2 **IN DISHAB 2.7K** Photo | 42.413 ± 3.670 | 1.108 | **0.2951** | 40.409 | **<0.0001** | 14 |
| **MBON15&18** MB050BG4 > ChR2 **IN DISHAB 3.5K** Photo | 52.925 ± 3.705 | 11.910 | **<0.0009** | 15.670 | **<0.0001** | 14 |
| **MBON15&18** MB050BG4 > ChR2 **IN DISHAB 17K** Photo | 63.568 ± 2.930 | 34.566 | **<0.0001** | 2.342 | **0.1293** | 14 |
| **Figure 10C: MBON15&18 MB050BG4 > UAS-ChR2-XXL4 Channel Rhodopsin (ChR2) NAIVE** | | | | | | |
| ANOVA $F_{(4,69)}$=18.5997 **p<0.0001** | | | | | | |
| **MBON15&18** MB050BG4 > ChR2 **IN** AVD Octanol | 70.276 ± 2.702 | 60.679 | **<0.0001** | | | 14 |
| **MBON15&18** MB050BG4 > ChR2 **IN** HAB Octanol | 37.798 ± 3.101 | | | 60.679 | **<0.0001** | 14 |
| **MBON15&18** MB050BG4 > ChR2 **IN PHOTO 2.7K** | 65.364 ± 3.502 | 43.712 | **<0.0001** | 1.388 | **0.2430** | 14 |
| **MBON15&18** MB050BG4 > ChR2 **IN PHOTO 3.5K** | 64.481 ± 2.613 | 40.958 | **<0.0001** | 1.931 | **0.1693** | 14 |
| **MBON15&18** MB050BG4 > ChR2 **IN PHOTO 17K** | 60.149 ± 2.727 | 28.738 | **<0.0001** | 5.899 | **0.0179** | 14 |
| **Figure 10E: MB110CG4 > UAS-ChR2-XXL4 Channel Rhodopsin (ChR2) (MBON8&9)** | | | | | | |
| ANOVA $F_{(6,97)}$=14.4155 **p<0.0001** | | | | | | |
| MB110CG4 > ChR2 **IN** AVD Octanol | 71.101 ± 3.002 | 65.767 | **<0.0001** | | | 14 |
| MB110CG4 > ChR2 **IN** HAB Octanol | 30.511 ± 3.806 | | | 65.767 | **<0.0001** | 14 |
| MB110CG4 > ChR2 **IN** DISHAB **SHOCK** | 68.262 ± 3.979 | 56.888 | **<0.0001** | 0.321 | **0.5719** | 14 |
| MB110CG4 > ChR2 **IN** DISHAB **YEAST** | 61.862 ± 2.843 | 39.235 | **<0.0001** | 3.407 | **0.0681** | 14 |
| MB110CG4 > ChR2 **IN DISHAB 2.7K** Photo | 57.094 ± 3.646 | 28.207 | **<0.0001** | 7.832 | **0.0062** | 14 |
| MB110CG4 > ChR2 **IN DISHAB 3.5K** Photo | 51.865 ± 3.994 | 18.202 | **<0.0001** | 14.771 | **<0.0003** | 14 |
| MB110CG4 > ChR2 **IN DISHAB 17K** Photo | 54.071 ± 3.315 | 22.156 | **<0.0001** | 11.577 | **<0.0010** | 14 |
| **Figure 10F: MB110CG4 > UAS-ChR2-XXL4 Channel Rhodopsin (ChR2) NAIVE (MBON8&9)** | | | | | | |
| ANOVA $F_{(4,59)}$=11.5890 **p<0.0001** | | | | | | |
| MB110CG4 > ChR2 **IN** AVD Octanol | 70.239 ± 4.436 | 25.928 | **<0.0001** | | | 12 |
| MB110CG4 > ChR2 **IN** HAB Octanol | 36.624 ± 4.769 | | | 25.928 | **<0.0001** | 12 |

| Genotype | Mean ± SEM | HAB Control | | AVD Control | | N |
|---|---|---|---|---|---|---|
| | | F-Ratio | p-value | F-Ratio | p-value | |
| MB110CG4 > ChR2 **IN PHOTO 2.7K** | 73.208 ± 4.615 | 30.711 | **<0.0001** | 0.202 | **0.6546** | 12 |
| MB110CG4 > ChR2 **IN PHOTO 3.5K** | 75.076 ± 5.202 | 33.927 | **<0.0001** | 0.536 | **0.4668** | 12 |
| MB110CG4 > ChR2 **IN PHOTO 17K** | 68.035 ± 4.259 | 22.640 | **<0.0001** | 0.111 | **0.7398** | 12 |
| **Figure 11 : Ellipsoid Body Neurons do not participate in Habituation nor Dishabituation.** | | | | | | |
| **Figure 11B: RDLG4 > $Shi^{ts}$** ($w^{1118}$ background = (+)) | | | | | | |
| ANOVA $F_{(11,167)}$=9.9745 **$p<0.0001$** | | | | | | |
| RDLG4 > (+) **IN** AVD Octanol | 64.643 ± 3.845 | 16.242 | **<0.0001** | | | 14 |
| RDLG4 > (+) **IN** HAB Octanol | 44.913 ± 3.176 | | | 16.242 | **<0.0001** | 14 |
| RDLG4 > (+) **IN** DISHAB **shock** | 65.023 ± 4.112 | 16.873 | **<0.0001** | 0.006 | **0.9383** | 14 |
| RDLG4 > (+) **IN** DISHAB **yeast** | 62.394 ± 3.003 | 12.750 | **<0.0005** | 0.211 | **0.6465** | 14 |
| RDLG4 > $Shi^{ts}$ **IN** AVD Octanol | 65.277 ± 4.465 | 28.225 | **<0.0001** | | | 14 |
| RDLG4 > $Shi^{ts}$ **IN** HAB Octanol | 39.268 ± 3.129 | | | 28.225 | **<0.0001** | 14 |
| RDLG4 > $Shi^{ts}$ **IN** DISHAB **shock** | 60.454 ± 3.723 | 18.727 | **<0.0001** | 0.970 | **0.3260** | 14 |
| RDLG4 > $Shi^{ts}$ **IN** DISHAB **yeast** | 56.862 ± 3.044 | 12.916 | **<0.0005** | 2.954 | **0.0876** | 14 |
| RDLG4 > $Shi^{ts}$ UN AVD Octanol | 68.271 ± 3.943 | 16.558 | **<0.0001** | | | 14 |
| RDLG4 > $Shi^{ts}$ UN HAB Octanol | 48.350 ± 3.530 | | | 16.558 | **<0.0001** | 14 |
| RDLG4 > $Shi^{ts}$ UN DISHAB **shock** | 76.095 ± 2.769 | 32.119 | **<0.0001** | 2.554 | **0.1120** | 14 |
| RDLG4 > $Shi^{ts}$ UN DISHAB **yeast** | 70.200 ± 2.108 | 19.920 | **<0.0001** | 0.155 | **0.6940** | 14 |
| **Figure 11C: RDLG4 > UAS-ChR2-XXL4 Channel Rhodopsin (ChR2)** | | | | | | |
| ANOVA $F_{(6,125)}$=10.8361 **$p<0.0001$** | | | | | | |
| RDLG4 > ChR2 **IN** AVD Octanol | 72.083 ± 2.886 | 25.814 | **<0.0001** | | | 18 |
| RDLG4 > ChR2 **IN** HAB Octanol | 42.854 ± 4.340 | | | 25.814 | **<0.0001** | 18 |
| RDLG4 > ChR2 **IN** DISHAB **SHOCK** | 68.828 ± 2.984 | 20.383 | **<0.0001** | 0.320 | **0.5725** | 18 |
| RDLG4 > ChR2 **IN** DISHAB **YEAST** | 67.261 ± 2.900 | 17.998 | **<0.0001** | 0.702 | **0.4035** | 18 |
| RDLG4 > ChR2 **IN DISHAB 2.7K** Photo | 48.825 ± 4.312 | 1.077 | **0.3013** | 16.344 | **<0.0001** | 18 |
| RDLG4 > ChR2 **IN DISHAB 3.5K** Photo | 44.076 ± 5.525 | 0.045 | **0.8322** | 23.701 | **<0.0001** | 18 |
| RDLG4 > ChR2 **IN DISHAB 17K** Photo | 43.244 ± 4.712 | 0.004 | **0.9460** | 25.130 | **<0.0001** | 18 |
| **Figure 11D: RDLG4 > UAS-ChR2-XXL4 Channel Rhodopsin (ChR2) NAIVE** | | | | | | |
| ANOVA $F_{(4,59)}$=11.5517 **$p<0.0001$** | | | | | | |
| RDLG4 > ChR2 **IN** AVD Octanol | 71.796 ± 3.449 | 32.090 | **<0.0001** | | | 12 |
| RDLG4 > ChR2 **IN** HAB Octanol | 36.135 ± 4.056 | | | 32.090 | **<0.0001** | 12 |
| RDLG4 > ChR2 **IN PHOTO 2.7K** | 63.214 ± 5.339 | 18.503 | **<0.0001** | 1.858 | **0.1783** | 12 |
| RDLG4 > ChR2 **IN PHOTO 3.5K** | 73.302 ± 4.335 | 34.858 | **<0.0001** | 0.057 | **0.8117** | 12 |
| RDLG4 > ChR2 **IN PHOTO 17K** | 66.650 ± 4.839 | 23.497 | **<0.0001** | 0.668 | **0.4171** | 12 |
| **Figure 12: Continuous exposure to Octanol following shock-induced dishabituation reinstates habituation.** | | | | | | |
| ANOVA $F_{(7,111)}$=12.2232 **$p<0.0001$** | | | | | | |
| $w^{1118}$ AVD Octanol 100% OCT | 55.733 ± 3.152 | 39.322 | **<0.0001** | | | 14 |
| $w^{1118}$ HAB Octanol 100% OCT | 27.391 ± 2.939 | | | 39.322 | **<0.0001** | 14 |
| $w^{1118}$ DISHAB **SHOCK** 100% OCT | 49.837 ± 3.303 | 24.662 | **<0.0001** | 1.701 | **0.1949** | 14 |
| $w^{1118}$ DISHAB **SHOCK** 0min **OCT** 100% OCT | 51.188 ± 2.791 | 27.722 | **<0.0001** | 1.011 | **0.3169** | 14 |
| $w^{1118}$ DISHAB **SHOCK** 1min **OCT** 100% OCT | 42.702 ± 3.317 | 11.475 | **<0.0010** | 8.312 | **0.0047** | 14 |
| $w^{1118}$ DISHAB **SHOCK** 2min **OCT** 100% OCT | 38.184 ± 3.020 | 5.702 | **0.0187** | 15.076 | **<0.0002** | 14 |
| $w^{1118}$ DISHAB **SHOCK** 3min **OCT** 100% OCT | 33.249 ± 3.556 | 1.679 | **0.1978** | 24.747 | **<0.0001** | 14 |
| $w^{1118}$ DISHAB **SHOCK** 4min **OCT** 100% OCT | 26.152 ± 3.413 | 0.075 | **0.7845** | 42.834 | **<0.0001** | 14 |
| **Supplemental Figure 1: Octanol-habituated flies cannot be dishabituated by a high-potency Octanol concentration.** | | | | | | |

| Genotype | Mean ± SEM | HAB Control | | AVD Control | | N |
|---|---|---|---|---|---|---|
| | | F-Ratio | p-value | F-Ratio | p-value | |
| **Supplemental Figure 1A: Habituation to 0.1,1,10% Octanol dilutions and dishabituation to 100% Octanol in $w^{1118}$ flies** | | | | | | |
| ANOVA $F_{(9,119)}$=17.9900 **$p<0.0001$** | | | | | | |
| $w^{1118}$ AVD Octanol 0.1% OCT | 22.056 ± 5.197 | 16.008 | **<0.0002** | | | 12 |
| $w^{1118}$ HAB Octanol 0.1% OCT | -1.935 ± 6.168 | | | 16.008 | **<0.0002** | 12 |
| $w^{1118}$ DISHAB **100% Octanol** 0.1% OCT | -4.400 ± 4.971 | 0.169 | **0.6817** | 19.467 | **<0.0001** | 12 |
| $w^{1118}$ AVD Octanol 1% OCT | 30.891 ± 3.554 | 16.096 | **<0.0002** | | | 12 |
| $w^{1118}$ HAB Octanol 1% OCT | 6.833 ± 3.858 | | | 16.096 | **<0.0002** | 12 |
| $w^{1118}$ DISHAB **100% Octanol** 1% OCT | 10.378 ± 3.358 | 0.349 | **0.5556** | 11.702 | **<0.0009** | 12 |
| $w^{1118}$ AVD Octanol 10% OCT | 47.933 ± 3.501 | 22.740 | **<0.0001** | | | 12 |
| $w^{1118}$ HAB Octanol 10% OCT | 19.338 ± 4.080 | | | 22.740 | **<0.0001** | 12 |
| $w^{1118}$ DISHAB **100% Octanol** 10% OCT | 23.638 ± 3.152 | 0.514 | **0.4748** | 16.415 | **<0.0001** | 12 |
| $w^{1118}$ DISHAB **SHOCK** 10% OCT | 45.525 ± 3.513 | 19.071 | **<0.0001** | 0.161 | **0.6888** | 12 |
| **Supplemental Figure 1B: Habituation to 20,50,100% Octanol dilutions and dishabituation to 100% Octanol in $w^{1118}$ flies** | | | | | | |
| ANOVA $F_{(2,35)}$=27.0659 **$p<0.0001$** | | | | | | |
| $w^{1118}$ AVD Octanol 20% OCT | 58.319 ± 3.294 | 44.261 | **<0.0001** | | | 12 |
| $w^{1118}$ HAB Octanol 20% OCT | 26.133 ± 3.917 | | | 44.261 | **<0.0001** | 12 |
| $w^{1118}$ DISHAB **100% Octanol** 20% OCT | 29.064 ± 2.985 | 0.366 | **0.5488** | 36.569 | **<0.0001** | 12 |
| ANOVA $F_{(2,53)}$=24.2193 **$p<0.0001$ (Full experiment shown in S. Figure 8B below)** | | | | | | |
| $w^{1118}$ AVD Octanol 50% OCT | 61.223 ± 4.584 | 35.357 | **<0.0001** | | | 18 |
| $w^{1118}$ HAB Octanol 50% OCT | 23.774 ± 4.095 | | | 35.357 | **<0.0001** | 18 |
| $w^{1118}$ DISHAB **100% Octanol** 50% OCT | 22.772 ± 4.659 | 0.025 | **0.8741** | 37.275 | **<0.0001** | 18 |
| ANOVA $F_{(2,35)}$=39.0738 **$p<0.0001$** | | | | | | |
| $w^{1118}$ AVD Octanol 100% OCT | 63.244 ± 2.997 | 58.345 | **<0.0001** | | | 12 |
| $w^{1118}$ HAB Octanol 100% OCT | 20.117 ± 3.634 | | | 58.345 | **<0.0001** | 12 |
| $w^{1118}$ DISHAB **100% Octanol** 100% OCT | 19.922 ± 5.061 | 0.001 | **0.9726** | 58.874 | **<0.0001** | 12 |
| **Supplemental Figure 2: $Orco^1$ mutant is unable to sense the yeast odor stimulus.** | | | | | | |
| **Supplemental Figure 2A: Anosmic Mutants AVOIDANCE Orco1** ($w^{1118}$ background = (+)) | | | | | | |
| ANOVA $F_{(3,39)}$=139.1718 **$p<0.0001$** | | | | | | |
| $w^{1118}$ AVD Octanol | 70.011 ± 1.942 | - | - | | | 10 |
| $Orco^1$ mutant AVD Octanol | 19.314 ± 2.843 | - | - | 250.462 | **<0.0001** | 10 |
| $w^{1118}$ AVD Yeast | 58.446 ± 1.855 | - | - | | | 10 |
| $Orco^1$ mutant AVD Yeast | 18.049 ± 2.285 | - | - | 159.032 | **<0.0001** | 10 |
| $w^{1118}$ AVD Octanol | 70.011 ± 1.942 | - | - | | | 10 |
| $w^{1118}$ AVD Yeast | 58.446 ± 1.855 | - | - | 13.033 | **<0.0001** | 10 |
| $Orco^1$ mutant AVD Octanol | 19.314 ± 2.843 | - | - | | | 10 |
| $Orco^1$ mutant AVD Yeast | 18.049 ± 2.285 | - | - | 0.156 | **0.6951** | 10 |
| **Supplemental Figure 2B: Anosmic Mutants DISHAB $Orco^1$** ($w^{1118}$ background = (+)) | | | | | | |
| ANOVA $F_{(5,95)}$=48.8938 **$p<0.0001$** | | | | | | |
| $w^{1118}$ AVD **Shock** | 61.302 ± 1.478 | 106.263 | **<0.0001** | | | 16 |
| $w^{1118}$ HAB **Shock** | 32.736 ± 2.338 | | | 106.263 | **<0.0001** | 16 |
| $w^{1118}$ DISHAB Yeast | 54.249 ± 2.111 | 60.271 | **<0.0001** | 6.476 | **0.0126** | 16 |
| $Orco^1$ mutant AVD **Shock** | 57.994 ± 1.849 | 74.703 | **<0.0001** | | | 16 |
| $Orco^1$ mutant HAB **Shock** | 34.044 ± 1.913 | | | 74.703 | **<0.0001** | 16 |
| $Orco^1$ mutant DISHAB Yeast | 32.768 ± 1.961 | 0.211 | **0.6463** | 82.874 | **<0.0001** | 16 |
| **Supplemental Figure 2C: Yeast exposures with a 30sec delay prior to Octanol avoidance in $w^{1118}$ control flies** | | | | | | |
| ANOVA $F_{(14,161)}$=3.3531 **$p<0.0001$** | | | | | | |

| Genotype | Mean ± SEM | HAB Control | | AVD Control | | N |
|---|---|---|---|---|---|---|
| | | F-Ratio | p-value | F-Ratio | p-value | |
| $w^{1118}$ AVD Octanol 0.1% OCT | 32.774 ± 4.447 | - | - | | | 10 |
| $w^{1118}$ Exposure **YEAST** 5sec 0.1% OCT | 32.361 ± 5.713 | - | - | 0.003 | **0.9509** | 10 |
| $w^{1118}$ Exposure **YEAST** 15sec 0.1% OCT | 29.549 ± 5.285 | - | - | 0.232 | **0.6303** | 10 |
| $w^{1118}$ AVD Octanol 1% OCT | 35.999 ± 5.161 | - | - | | | 10 |
| $w^{1118}$ Exposure **YEAST** 5sec 1% OCT | 45.941 ± 4.541 | - | - | 2.210 | **0.1392** | 10 |
| $w^{1118}$ Exposure **YEAST** 15sec 1% OCT | 39.778 ± 5.855 | - | - | 0.319 | **0.5728** | 10 |
| $w^{1118}$ AVD Octanol 10% OCT | 39.952 ± 4.147 | - | - | | | 10 |
| $w^{1118}$ Exposure **YEAST** 5sec 10% OCT | 47.042 ± 3.765 | - | - | 1.124 | **0.2907** | 10 |
| $w^{1118}$ Exposure **YEAST** 15sec 10% OCT | 50.700 ± 2.855 | - | - | 2.583 | **0.1101** | 10 |
| $w^{1118}$ AVD Octanol 50% OCT | 51.814 ± 3.357 | - | - | | | 10 |
| $w^{1118}$ Exposure **YEAST** 5sec 50% OCT | 46.637 ± 3.787 | - | - | 0.599 | **0.4400** | 10 |
| $w^{1118}$ Exposure **YEAST** 15sec 50% OCT | 51.837 ± 4.542 | - | - | <0.001 | **0.9973** | 10 |
| $w^{1118}$ AVD Octanol 100% OCT | 56.083 ± 5.124 | - | - | | | 14 |
| $w^{1118}$ Exposure **YEAST** 5sec 100% OCT | 51.194 ± 3.755 | - | - | 0.748 | **0.3883** | 14 |
| $w^{1118}$ Exposure **YEAST** 15sec 100% OCT | 47.922 ± 4.208 | - | - | 2.085 | **0.1508** | 14 |
| **Supplemental Figure 3: Photoactivation Controls** | | | | | | |
| **Supplemental Figure 3A: Canton S Photoactivation (Control)** | | | | | | |
| ANOVA $F_{(6,97)}=43.9040$ **p<0.0001** | | | | | | |
| Canton S AVD Octanol | 72.611 ± 3.395 | 66.902 | **<0.0001** | | | 14 |
| Canton S HAB Octanol | 36.017 ± 3.787 | | | 66.902 | **<0.0001** | 14 |
| Canton S DISHAB **SHOCK** | 77.987 ± 2.250 | 88.001 | **<0.0001** | 1.443 | **0.2326** | 14 |
| Canton S DISHAB **YEAST** | 73.492 ± 2.536 | 70.160 | **<0.0001** | 0.038 | **0.8444** | 14 |
| Canton S **DISHAB 2.7K** Photo | 37.451 ± 2.668 | 0.102 | **0.7492** | 61.761 | **<0.0001** | 14 |
| Canton S **DISHAB 3.5K** Photo | 34.006 ± 3.178 | 0.202 | **0.6541** | 74.458 | **<0.0001** | 14 |
| Canton S **DISHAB 17K** Photo | 35.134 ± 3.932 | 0.039 | **0.8438** | 70.172 | **<0.0001** | 14 |
| **Supplemental Figure 3B: Canton S Photoactivation (Control) NAÏVE** | | | | | | |
| ANOVA $F_{(4,69)}=28.5560$ **p<0.0001** | | | | | | |
| Canton S AVD Octanol | 72.611 ± 3.395 | 67.624 | **<0.0001** | | | 14 |
| Canton S HAB Octanol | 36.017 ± 3.787 | | | 67.624 | **<0.0001** | 14 |
| Canton S **PHOTO 2.7K** | 76.778 ± 2.870 | 83.901 | **<0.0001** | 0.876 | **0.3525** | 14 |
| Canton S **PHOTO 3.5K** | 72.761 ± 2.698 | 68.180 | **<0.0001** | 0.001 | **0.9732** | 14 |
| Canton S **PHOTO 17K** | 71.146 ± 2.848 | 62.319 | **<0.0001** | 0.108 | **0.7430** | 14 |
| **Supplemental Figure 3C: $w^{1118}$ Photoactivation (Control)** | | | | | | |
| ANOVA $F_{(6,83)}=22.3832$ **p<0.0001** | | | | | | |
| $w^{1118}$ AVD Octanol | 70.185 ± 2.744 | 74.479 | **<0.0001** | | | 12 |
| $w^{1118}$ HAB Octanol | 34.607 ± 2.994 | | | 74.479 | **<0.0001** | 12 |
| $w^{1118}$ DISHAB **SHOCK** | 61.847 ± 2.815 | 43.659 | **<0.0001** | 4.091 | **0.0465** | 12 |
| $w^{1118}$ DISHAB **YEAST** | 60.991 ± 3.031 | 40.960 | **<0.0001** | 4.973 | **0.0286** | 12 |
| $w^{1118}$ **DISHAB 2.7K** Photo | 42.380 ± 2.467 | 3.555 | **0.0631** | 45.489 | **<0.0001** | 12 |
| $w^{1118}$ **DISHAB 3.5K** Photo | 42.085 ± 3.021 | 3.290 | **0.0735** | 46.460 | **<0.0001** | 12 |
| $w^{1118}$ **DISHAB 17K** Photo | 39.264 ± 3.262 | 1.276 | **0.2620** | 56.255 | **<0.0001** | 12 |
| **Supplemental Figure 3D: $w^{1118}$ Photoactivation (Control) NAÏVE** | | | | | | |
| ANOVA $F_{(4,59)}=57.1462$ **p<0.0001** | | | | | | |
| $w^{1118}$ AVD Octanol | 70.185 ± 2.744 | 113.502 | **<0.0001** | | | 12 |
| $w^{1118}$ HAB Octanol | 34.607 ± 2.994 | | | 113.502 | **<0.0001** | 12 |
| $w^{1118}$ **PHOTO 2.7K** | 75.961 ± 2.178 | 153.344 | **<0.0001** | 2.991 | **0.0893** | 12 |

| Genotype | Mean ± SEM | HAB Control | | AVD Control | | N |
|---|---|---|---|---|---|---|
| | | F-Ratio | p-value | F-Ratio | p-value | |
| $w^{1118}$ **PHOTO 3.5K** | 74.580 ± 1.622 | 143.280 | **<0.0001** | 1.732 | **0.1935** | 12 |
| $w^{1118}$ **PHOTO 17K** | 75.953 ± 2.001 | 153.288 | **<0.0001** | 2.983 | **0.0897** | 12 |
| **Supplemental Figure 3E: Isogenic Photoactivation sibling Control (ChR2 > TM3sb) A** | | | | | | |
| ANOVA $F_{(6,97)}$=23.9811 **p<0.0001** | | | | | | |
| ChR2 > TM3sb **IN** AVD Octanol | 72.360 ± 2.788 | 58.252 | **<0.0001** | | | 14 |
| ChR2 > TM3sb **IN** HAB Octanol | 37.706 ± 3.016 | | | 58.252 | **<0.0001** | 14 |
| ChR2 > TM3sb **IN** DISHAB **SHOCK** | 68.939 ± 3.220 | 47.319 | **<0.0001** | 0.567 | **0.4531** | 14 |
| ChR2 > TM3sb **IN** DISHAB **YEAST** | 67.521 ± 2.836 | 43.120 | **<0.0001** | 1.135 | **0.2893** | 14 |
| ChR2 > TM3sb **IN DISHAB 2.7K** Photo | 38.018 ± 3.286 | 0.004 | **0.9453** | 57.208 | **<0.0001** | 14 |
| ChR2 > TM3sb **IN DISHAB 3.5K** Photo | 44.472 ± 3.188 | 2.220 | **0.1396** | 37.727 | **<0.0001** | 14 |
| ChR2 > TM3sb **IN DISHAB 17K** Photo | 42.401 ± 3.985 | 1.069 | **0.3038** | 43.538 | **<0.0001** | 14 |
| **Supplemental Figure 3F: Isogenic Photoactivation sibling Control (ChR2 > TM3sb) A NAÏVE** | | | | | | |
| ANOVA $F_{(4,59)}$=16.6188 **p<0.0001** | | | | | | |
| ChR2 > TM3sb **IN** AVD Octanol | 66.733 ± 3.068 | 40.890 | **<0.0001** | | | 12 |
| ChR2 > TM3sb **IN** HAB Octanol | 36.862 ± 3.406 | | | 40.890 | **<0.0001** | 12 |
| ChR2 > TM3sb **IN PHOTO 2.7K** | 68.971 ± 2.893 | 47.247 | **<0.0001** | 0.229 | **0.6337** | 12 |
| ChR2 > TM3sb **IN PHOTO 3.5K** | 67.021 ± 3.071 | 41.683 | **<0.0001** | 0.003 | **0.9510** | 12 |
| ChR2 > TM3sb **IN PHOTO 17K** | 64.205 ± 3.966 | 34.261 | **<0.0001** | 0.292 | **0.5905** | 12 |
| **Supplemental Figure 3G: Isogenic Photoactivation sibling Control (ChR2 > CyO) B** | | | | | | |
| ANOVA $F_{(6,97)}$=17.6586 **p<0.0001** | | | | | | |
| ChR2 > CyO **IN** AVD Octanol | 76.618 ± 3.000 | 26.373 | **<0.0001** | | | 14 |
| ChR2 > CyO **IN** HAB Octanol | 53.366 ± 3.465 | | | 26.373 | **<0.0001** | 14 |
| ChR2 > CyO **IN** DISHAB **SHOCK** | 77.557 ± 2.802 | 28.547 | **<0.0001** | 0.043 | **0.8360** | 14 |
| ChR2 > CyO **IN** DISHAB **YEAST** | 82.961 ± 1.978 | 42.724 | **<0.0001** | 1.962 | **0.1646** | 14 |
| ChR2 > CyO **IN DISHAB 2.7K** Photo | 57.218 ± 3.473 | 0.723 | **0.3970** | 18.358 | **<0.0001** | 14 |
| ChR2 > CyO **IN DISHAB 3.5K** Photo | 54.741 ± 3.600 | 0.092 | **0.7621** | 23.347 | **<0.0001** | 14 |
| ChR2 > CyO **IN DISHAB 17K** Photo | 51.976 ± 3.733 | 0.094 | **0.7595** | 29.621 | **<0.0001** | 14 |
| **Supplemental Figure 3H: Isogenic Photoactivation sibling Control (ChR2 > CyO) B NAÏVE** | | | | | | |
| ANOVA $F_{(4,69)}$=33.7809 **p<0.0001** | | | | | | |
| ChR2 > CyO **IN** AVD Octanol | 77.750 ± 1.869 | 83.917 | **<0.0001** | | | 14 |
| ChR2 > CyO **IN** HAB Octanol | 51.055 ± 2.819 | | | 83.917 | **<0.0001** | 14 |
| ChR2 > CyO **IN PHOTO 2.7K** | 81.343 ± 1.742 | 108.026 | **<0.0001** | 1.520 | **0.2220** | 14 |
| ChR2 > CyO **IN PHOTO 3.5K** | 75.580 ± 1.683 | 70.826 | **<0.0001** | 0.554 | **0.4590** | 14 |
| ChR2 > CyO **IN PHOTO 17K** | 73.420 ± 1.979 | 58.902 | **<0.0001** | 2.207 | **0.1421** | 14 |
| ***Supplemental Figure 4: Sibling Controls for MB neuronal subsets.** | | | | | | |
| Sibling control data are presented in Figure 4 and Supplemental Figure 5 tables. These data were separated from the respective figures to enhance the representation of behavioral experiments. **(S. Figure 4B: 17dG4 > (+), 4C: c739G4 > (+), 4E: VT030604G4 > (+), 4F: c305aG4 > (+), 4H: VT044966G4 > (+), 4I: MB131BG4 > (+), 4K: 17dG4 ; VT044966G4 > (+)** | | | | | | |
| **Supplemental Figure 5: Distinct MBns regulate olfactory habituation and dishabituation.** | | | | | | |
| **Supplemental Figure 5B: (α/β)c739G4 > $Shi^{ts}$** ($w^{1118}$ background = (+)) | | | | | | |
| ANOVA $F_{(11,167)}$=9.9821 **p<0.0001** | | | | | | |
| (α/β)c739G4 > (+) **IN** AVD Octanol | 80.635 ± 2.469 | 35.399 | **<0.0001** | | | 14 |
| (α/β)c739G4 > (+) **IN** HAB Octanol | 53.370 ± 4.171 | | | 35.399 | | 14 |
| (α/β)c739G4 > (+) **IN** DISHAB **shock** | 73.539 ± 2.381 | 19.370 | **<0.0001** | 2.397 | **0.1235** | 14 |

| Genotype | Mean ± SEM | HAB Control | | AVD Control | | N |
|---|---|---|---|---|---|---|
| | | F-Ratio | p-value | F-Ratio | p-value | |
| (α/β)c739G4 > (+) **IN** DISHAB **yeast** | 76.220 ± 2.990 | 24.863 | **<0.0001** | 0.928 | **0.3368** | 14 |
| (α/β)c739G4 > $Shi^{ts}$ **IN** AVD Octanol | 75.877 ± 2.838 | 25.587 | **<0.0001** | | | 14 |
| (α/β)c739G4 > $Shi^{ts}$ **IN** HAB Octanol | 52.697 ± 3.228 | | | 25.587 | **<0.0001** | 14 |
| (α/β)c739G4 > $Shi^{ts}$ **IN** DISHAB **shock** | 71.724 ± 3.070 | 17.238 | **<0.0001** | 0.821 | **0.3661** | 14 |
| (α/β)c739G4 > $Shi^{ts}$ **IN** DISHAB **yeast** | 69.870 ± 3.228 | 14.043 | **<0.0003** | 1.718 | **0.1918** | 14 |
| (α/β)c739G4 > $Shi^{ts}$ UN AVD Octanol | 71.753 ± 4.096 | 22.300 | **<0.0001** | | | 14 |
| (α/β)c739G4 > $Shi^{ts}$ UN HAB Octanol | 50.113 ± 3.884 | | | 22.300 | **<0.0001** | 14 |
| (α/β)c739G4 > $Shi^{ts}$ UN DISHAB **shock** | 73.778 ± 3.150 | 26.668 | **<0.0001** | 0.195 | **0.6592** | 14 |
| (α/β)c739G4 > $Shi^{ts}$ UN DISHAB **yeast** | 68.630 ± 2.787 | 16.328 | **<0.0001** | 0.464 | **0.4965** | 14 |
| **Supplemental Figure 5C: (α/β)c739G4 > UAS-ChR2-XXL4 Channel Rhodopsin (ChR2)** | | | | | | |
| ANOVA $F_{(6,97)}$=7.4878 **p<0.0001** | | | | | | |
| (α/β)c739G4 > ChR2 **IN** AVD Octanol | 88.252 ± 2.506 | 41.759 | **<0.0001** | | | 14 |
| (α/β)c739G4 > ChR2 **IN** HAB Octanol | 64.305 ± 2.813 | | | 41.759 | **<0.0001** | 14 |
| (α/β)c739G4 > ChR2 **IN** DISHAB **SHOCK** | 78.727 ± 1.608 | 15.146 | **<0.0002** | 6.606 | **0.0117** | 14 |
| (α/β)c739G4 > ChR2 **IN** DISHAB **YEAST** | 79.751 ± 1.784 | 17.373 | **<0.0001** | 5.262 | **0.0240** | 14 |
| (α/β)c739G4 > ChR2 **IN DISHAB 2.7K** Photo | 74.756 ± 2.776 | 7.953 | **0.0058** | 13.264 | **<0.0005** | 14 |
| (α/β)c739G4 > ChR2 **IN DISHAB 3.5K** Photo | 79.664 ± 3.654 | 17.178 | **<0.0001** | 5.370 | **0.0227** | 14 |
| (α/β)c739G4 > ChR2 **IN DISHAB 17K** Photo | 77.453 ± 2.650 | 12.588 | **<0.0007** | 8.492 | **0.0044** | 14 |
| **Supplemental Figure 5D: (α/β)c739G4 > UAS-ChR2-XXL4 Channel Rhodopsin (ChR2) NAIVE** | | | | | | |
| ANOVA $F_{(4,69)}$=10.7846 **p<0.0001** | | | | | | |
| (α/β)c739G4 > ChR2 **IN** AVD Octanol | 82.378 ± 1.629 | 37.467 | **<0.0001** | | | 14 |
| (α/β)c739G4 > ChR2 **IN** HAB Octanol | 57.545 ± 4.033 | | | 37.467 | **<0.0001** | 14 |
| (α/β)c739G4 > ChR2 **IN PHOTO 2.7K** | 77.895 ± 2.045 | 25.160 | **<0.0001** | 1.221 | **0.2732** | 14 |
| (α/β)c739G4 > ChR2 **IN PHOTO 3.5K** | 74.099 ± 3.060 | 16.650 | **<0.0002** | 4.163 | **0.0453** | 14 |
| (α/β)c739G4 > ChR2 **IN PHOTO 17K** | 74.930 ± 2.945 | 18.361 | **<0.0001** | 3.370 | **0.0709** | 14 |
| **Supplemental Figure 5F: (α´/β´)c305aG4 > $Shi^{ts}$** ($w^{1118}$ background = (+)) | | | | | | |
| ANOVA $F_{(11,167)}$=18.4602 **p<0.0001** | | | | | | |
| (α´/β´)c305aG4 > (+) **IN** AVD Octanol | 80.772 ± 2.450 | 47.859 | **<0.0001** | | | 14 |
| (α´/β´)c305aG4 > (+) **IN** HAB Octanol | 51.641 ± 4.566 | | | 47.859 | **<0.0001** | 14 |
| (α´/β´)c305aG4 > (+) **IN** DISHAB **shock** | 78.000 ± 3.085 | 39.186 | **<0.0001** | 0.433 | **0.5113** | 14 |
| (α´/β´)c305aG4 > (+) **IN** DISHAB **yeast** | 70.271 ± 3.273 | 19.573 | **<0.0001** | 6.219 | **0.0136** | 14 |
| (α´/β´)c305aG4 > $Shi^{ts}$ **IN** AVD Octanol | 53.313 ± 2.794 | 1.360 | **0.2452** | | | 14 |
| (α´/β´)c305aG4 > $Shi^{ts}$ **IN** HAB Octanol | 48.402 ± 2.431 | | | 1.360 | **0.2452** | 14 |
| (α´/β´)c305aG4 > $Shi^{ts}$ **IN** DISHAB **shock** | 58.681 ± 2.948 | 5.959 | **0.0157** | 1.625 | **0.2042** | 14 |
| (α´/β´)c305aG4 > $Shi^{ts}$ **IN** DISHAB **yeast** | 52.460 ± 3.835 | 0.928 | **0.3366** | 0.041 | **0.8397** | 14 |
| (α´/β´)c305aG4 > $Shi^{ts}$ UN AVD Octanol | 77.101 ± 1.340 | 41.331 | **<0.0001** | | | 14 |
| (α´/β´)c305aG4 > $Shi^{ts}$ UN HAB Octanol | 50.030 ± 2.863 | | | 41.331 | **<0.0001** | 14 |
| (α´/β´)c305aG4 > $Shi^{ts}$ UN DISHAB **shock** | 75.938 ± 2.444 | 37.856 | **<0.0001** | 0.076 | **0.7827** | 14 |
| (α´/β´)c305aG4 > $Shi^{ts}$ UN DISHAB **yeast** | 74.160 ± 2.488 | 32.836 | **<0.0001** | 0.488 | **0.4858** | 14 |
| **Supplemental Figure 5G: (α´/β´)c305aG4 > UAS-ChR2-XXL4 Channel Rhodopsin (ChR2)** | | | | | | |
| ANOVA $F_{(6,97)}$=14.9077 **p<0.0001** | | | | | | |
| (α´/β´)c305aG4 > ChR2 **IN** AVD Octanol | 77.752 ± 2.292 | 34.623 | **<0.0001** | | | 14 |
| (α´/β´)c305aG4 > ChR2 **IN** HAB Octanol | 56.377 ± 2.518 | | | 34.623 | **<0.0001** | 14 |
| (α´/β´)c305aG4 > ChR2 **IN** DISHAB **SHOCK** | 78.142 ± 1.953 | 35.899 | **<0.0001** | 0.011 | **0.9146** | 14 |
| (α´/β´)c305aG4 > ChR2 **IN** DISHAB **YEAST** | 69.200 ± 2.406 | 12.461 | **<0.0007** | 5.541 | **0.0207** | 14 |
| (α´/β´)c305aG4 > ChR2 **IN DISHAB 2.7K** Photo | 59.743 ± 2.720 | 0.858 | **0.3564** | 24.575 | **<0.0001** | 14 |

| Genotype | Mean ± SEM | HAB Control | | AVD Control | | N |
|---|---|---|---|---|---|---|
| | | F-Ratio | p-value | F-Ratio | p-value | |
| (α′/β′)c305aG4 > ChR2 **IN DISHAB 3.5K** Photo | 59.568 ± 2.751 | 0.771 | **0.3819** | 25.055 | **<0.0001** | 14 |
| (α′/β′)c305aG4 > ChR2 **IN DISHAB 17K** Photo | 54.717 ± 3.163 | 0.208 | **0.6487** | 40.210 | **<0.0001** | 14 |
| **Supplemental Figure 5H: (α′/β′)c305aG4 > UAS-ChR2-XXL4 Channel Rhodopsin (ChR2) NAIVE** | | | | | | |
| ANOVA $F_{(4,69)}$=13.1673 **p<0.0001** | | | | | | |
| (α′/β′)c305aG4 > ChR2 **IN** AVD Octanol | 77.752 ± 2.292 | 36.390 | **<0.0001** | | | 14 |
| (α′/β′)c305aG4 > ChR2 **IN** HAB Octanol | 56.377 ± 2.518 | | | 36.390 | **<0.0001** | 14 |
| (α′/β′)c305aG4 > ChR2 **IN PHOTO 2.7K** | 70.424 ± 2.419 | 15.717 | **<0.0002** | 4.276 | **0.0426** | 14 |
| (α′/β′)c305aG4 > ChR2 **IN PHOTO 3.5K** | 72.304 ± 2.895 | 20.205 | **<0.0001** | 2.363 | **0.1290** | 14 |
| (α′/β′)c305aG4 > ChR2 **IN PHOTO 17K** | 59.094 ± 2.356 | 0.588 | **0.4459** | 27.725 | **<0.0001** | 14 |
| **Supplemental Figure 5J: (γ)MB131BG4 > $Shi^{ts}$** ($w^{1118}$ background = (+)) | | | | | | |
| ANOVA $F_{(11,167)}$=13.4243 **p<0.0001** | | | | | | |
| (γ) MB131BG4 > (+) **IN** AVD Octanol | 74.133 ± 3.812 | 42.174 | **<0.0001** | | | 14 |
| (γ) MB131BG4 > (+) **IN** HAB Octanol | 39.538 ± 5.473 | | | 42.174 | **<0.0001** | 14 |
| (γ) MB131BG4 > (+) **IN** DISHAB **shock** | 67.522 ± 3.885 | 27.596 | **<0.0001** | 1.540 | **0.2164** | 14 |
| (γ) MB131BG4 > (+) **IN** DISHAB **yeast** | 68.884 ± 4.000 | 30.348 | **<0.0001** | 0.970 | **0.3260** | 14 |
| (γ) MB131BG4 > $Shi^{ts}$ **IN** AVD Octanol | 68.840 ± 3.901 | 31.049 | **<0.0001** | | | 14 |
| (γ) MB131BG4 > $Shi^{ts}$ **IN** HAB Octanol | 39.157 ± 3.523 | | | 31.049 | **<0.0001** | 14 |
| (γ) MB131BG4 > $Shi^{ts}$ **IN** DISHAB **shock** | 66.759 ± 4.631 | 26.847 | **<0.0001** | 0.152 | **0.6965** | 14 |
| (γ) MB131BG4 > $Shi^{ts}$ **IN** DISHAB **yeast** | 61.707 ± 4.163 | 17.919 | **<0.0001** | 1.793 | **0.1824** | 14 |
| (γ) MB131BG4 > $Shi^{ts}$ UN AVD Octanol | 80.583 ± 2.279 | 34.488 | **<0.0001** | | | 14 |
| (γ) MB131BG4 > $Shi^{ts}$ UN HAB Octanol | 49.299 ± 3.440 | | | 34.488 | **<0.0001** | 14 |
| (γ) MB131BG4 > $Shi^{ts}$ UN DISHAB **shock** | 75.302 ± 1.980 | 23.828 | **<0.0001** | 0.982 | **0.3230** | 14 |
| (γ) MB131BG4 > $Shi^{ts}$ UN DISHAB **yeast** | 73.288 ± 2.700 | 20.280 | **<0.0001** | 1.875 | **0.1728** | 14 |
| **Supplemental Figure 5K: (γ)MB131BG4 > UAS-ChR2-XXL4 Channel Rhodopsin (ChR2)** | | | | | | |
| ANOVA $F_{(6,118)}$=21.4291 **p<0.0001** | | | | | | |
| (γ) MB131BG4 > ChR2 **IN** AVD Octanol | 82.242 ± 1.751 | 94.445 | **<0.0001** | | | 17 |
| (γ) MB131BG4 > ChR2 **IN** HAB Octanol | 44.244 ± 3.099 | | | 94.445 | **<0.0001** | 17 |
| (γ) MB131BG4 > ChR2 **IN** DISHAB **SHOCK** | 78.315 ± 1.832 | 75.933 | **<0.0001** | 1.008 | **0.3173** | 17 |
| (γ) MB131BG4 > ChR2 **IN** DISHAB **YEAST** | 78.342 ± 2.369 | 76.057 | **<0.0001** | 0.994 | **0.3207** | 17 |
| (γ) MB131BG4 > ChR2 **IN DISHAB 2.7K** Photo | 64.900 ± 3.360 | 27.910 | **<0.0001** | 19.671 | **<0.0001** | 17 |
| (γ) MB131BG4 > ChR2 **IN DISHAB 3.5K** Photo | 69.576 ± 3.529 | 41.976 | **<0.0001** | 10.493 | **0.0015** | 17 |
| (γ) MB131BG4 > ChR2 **IN DISHAB 17K** Photo | 72.152 ± 2.846 | 50.948 | **<0.0001** | 6.659 | **0.0111** | 17 |
| **Supplemental Figure 5L: (γ)MB131BG4 > UAS-ChR2-XXL4 Channel Rhodopsin (ChR2) NAIVE** | | | | | | |
| ANOVA $F_{(4,69)}$=16.0731 **p<0.0001** | | | | | | |
| (γ) MB131BG4 > ChR2 **IN** AVD Octanol | 75.162 ± 2.519 | 46.739 | **<0.0001** | | | 14 |
| (γ) MB131BG4 > ChR2 **IN** HAB Octanol | 47.180 ± 3.225 | | | 46.739 | **<0.0001** | 14 |
| (γ) MB131BG4 > ChR2 **IN PHOTO 2.7K** | 71.952 ± 3.124 | 36.628 | **<0.0001** | 0.615 | **0.4356** | 14 |
| (γ) MB131BG4 > ChR2 **IN PHOTO 3.5K** | 72.621 ± 2.790 | 38.635 | **<0.0001** | 0.385 | **0.5368** | 14 |
| (γ) MB131BG4 > ChR2 **IN PHOTO 17K** | 72.121 ± 2.754 | 37.131 | **<0.0001** | 0.552 | **0.4601** | 14 |
| ***Supplemental Figure 6: Sibling Controls for Dopaminergic Neurons.** | | | | | | |
| Sibling control data are presented in Figure 5 table and were separated to enhance the representation of behavioral experiments. **(S. Figure 5A: pleG4 > (+), 5B: PAMG4 > (+) & $Shi^{ts}$ > (+), 5C: PPL1G4 > (+)** | | | | | | |
| ***Supplemental Figure 7: Isogenic Controls for APL Neurons.** | | | | | | |
| Sibling control data are presented in Figure 8 table and were separated to enhance the representation of behavioral experiments. **(S. Figure 7A: APLG4 > (+), 7B: APLG4G80$^{ts}$ > (+) & GAD-RNAi > (+), 7C: APLG4G80$^{ts}$ > (+) (A)** | | | | | | |

| Genotype | Mean ± SEM | HAB Control | | AVD Control | | N |
|---|---|---|---|---|---|---|
| | | F-Ratio | p-value | F-Ratio | p-value | |
| **Supplemental Figure 7D, 7E: G80$^{ts}$ ; APLG4 > TBH-RNAi B** (w$^{1118}$ background = (+)) VDRC 51667 | | | | | | |
| ANOVA $F_{(11,155)}$=9.8344 **p<0.0001** | | | | | | |
| G80$^{ts}$ ; APLG4 > (+) **IN** AVD Octanol | 74.998 ± 2.915 | 20.362 | **<0.0001** | | | 13 |
| G80$^{ts}$ ; APLG4 > (+) **IN** HAB Octanol | 52.216 ± 4.315 | | | 20.362 | **<0.0001** | 13 |
| G80$^{ts}$ ; APLG4 > (+) **IN** DISHAB **shock** | 71.877 ± 4.414 | 15.166 | **<0.0002** | 0.3820 | **0.5374** | 13 |
| G80$^{ts}$ ; APLG4 > (+) **IN** DISHAB **yeast** | 70.167 ± 3.633 | 12.642 | **<0.0006** | 0.9153 | **0.3402** | 13 |
| G80$^{ts}$ ; APLG4 > TBH-RNAi **IN** AVD Octanol | 82.445 ± 2.855 | 40.058 | **<0.0001** | | | 13 |
| G80$^{ts}$ ; APLG4 > TBH-RNAi **IN** HAB Octanol | 50.491 ± 3.667 | | | 40.058 | **<0.0001** | 13 |
| G80$^{ts}$ ; APLG4 > TBH-RNAi **IN** DISHAB **shock** | 67.335 ± 3.602 | 11.129 | **0.0011** | 8.958 | **0.0032** | 13 |
| G80$^{ts}$ ; APLG4 > TBH-RNAi **IN** DISHAB **yeast** | 67.930 ± 4.432 | 11.930 | **<0.0008** | 8.266 | **0.0046** | 13 |
| G80$^{ts}$ ; APLG4 > TBH-RNAi UN AVD Octanol | 76.780 ± 3.223 | 23.321 | **<0.0001** | | | 13 |
| G80$^{ts}$ ; APLG4 > TBH-RNAi UN HAB Octanol | 52.399 ± 4.157 | | | 23.321 | **<0.0001** | 13 |
| G80$^{ts}$ ; APLG4 > TBH-RNAi UN DISHAB **shock** | 79.692 ± 2.186 | 29.225 | **<0.0001** | 0.332 | **0.5649** | 13 |
| G80$^{ts}$ ; APLG4 > TBH-RNAi UN DISHAB **yeast** | 77.660 ± 2.535 | 25.035 | **<0.0001** | 0.030 | **0.8618** | 13 |
| **Supplemental Figure 7F: GADG4 > UAS-ChR2-XXL4 Channel Rhodopsin (ChR2)** | | | | | | |
| ANOVA $F_{(6,139)}$=11.0064 **p<0.0001** | | | | | | |
| GADG4 > ChR2 **IN** AVD Octanol | 73.612 ± 2.632 | 25.586 | **<0.0001** | | | 20 |
| GADG4 > ChR2 **IN** HAB Octanol | 47.371 ± 3.479 | | | 25.586 | **<0.0001** | 20 |
| GADG4 > ChR2 **IN** DISHAB **SHOCK** | 71.068 ± 3.100 | 20.865 | **<0.0001** | 0.240 | **0.6246** | 20 |
| GADG4 > ChR2 **IN** DISHAB **YEAST** | 70.841 ± 3.388 | 20.468 | **<0.0001** | 0.285 | **0.5941** | 20 |
| GADG4 > ChR2 **IN DISHAB 2.7K** Photo | 65.093 ± 4.167 | 11.669 | **<0.0009** | 2.697 | **0.1028** | 20 |
| GADG4 > ChR2 **IN DISHAB 3.5K** Photo | 59.138 ± 4.442 | 5.144 | **0.0249** | 7.784 | **0.0060** | 20 |
| GADG4 > ChR2 **IN DISHAB 17K** Photo | 42.925 ± 4.118 | 0.734 | **0.3929** | 34.991 | **<0.0001** | 20 |
| **Supplemental Figure 7G: GADG4 > UAS-ChR2-XXL4 Channel Rhodopsin (ChR2) NAIVE** | | | | | | |
| ANOVA $F_{(4,99)}$=11.7010 **p<0.0001** | | | | | | |
| GADG4 > ChR2 **IN** AVD Octanol | 73.612 ± 2.632 | 24.756 | **<0.0001** | | | 20 |
| GADG4 > ChR2 **IN** HAB Octanol | 47.371 ± 3.479 | | | 24.756 | **<0.0001** | 20 |
| GADG4 > ChR2 **IN PHOTO 2.7K** | 66.346 ± 4.328 | 12.944 | **<0.0006** | 1.898 | **0.1715** | 20 |
| GADG4 > ChR2 **IN PHOTO 3.5K** | 66.135 ± 3.857 | 12.658 | **<0.0006** | 2.009 | **0.1595** | 20 |
| GADG4 > ChR2 **IN PHOTO 17K** | 44.875 ± 4.109 | 0.224 | **0.6370** | 29.691 | **<0.0001** | 20 |
| ***Supplemental Figure 8: Isogenic Controls for MBONs and Ellipsoid Body Neurons** | | | | | | |
| Sibling control data are presented in Figure 9,10 and 11 table and were separated to enhance the representation of behavioral experiments. **(S. Figure 8A: MB112CG4 > (+), 8B: MB027BG4 > (+), 8D: RDLG4 > (+)** | | | | | | |
| **Supplemental Figure 8C: MB543BG4 > Shi$^{ts}$** (w$^{1118}$ background = (+)) | | | | | | |
| ANOVA $F_{(5,57)}$=4.9556 **p<0.001** | | | | | | |
| MB543BG4 > (+) IN AVD Octanol | 63.396 ± 5.491 | 14.720 | **<0.0004** | | | 10 |
| MB543BG4 > (+) IN HAB Octanol | 32.125 ± 3.798 | | | 14.720 | **<0.0004** | 10 |
| MB543BG4 > (+) IN DISHAB **SHOCK** | 66.049 ± 6.387 | 15.399 | **<0.0003** | 0.094 | **0.7601** | 8 |
| MB543BG4 > Shi$^{ts}$ IN AVD Octanol | 63.803 ± 4.413 | 1.618 | **0.2089** | | | 10 |
| MB543BG4 > Shi$^{ts}$ IN HAB Octanol | 53.434 ± 7.614 | | | 1.618 | **0.2089** | 10 |
| MB543BG4 > Shi$^{ts}$ IN DISHAB **SHOCK** | 63.271 ± 6.669 | 1.456 | **0.2329** | 0.004 | **0.9481** | 10 |
| **Supplemental Figure 10: Strains expressing Shibire$^{ts}$ exhibit normal OCT avoidance under permissive and non-permissive conditions.** | | | | | | |
| **Supplemental Figure 10A: dncG4 > Shi$^{ts}$ AVOIDANCE** (w$^{1118}$ background = (+)) | | | | | | |
| ANOVA $F_{(1,23)}$=0.2279 **p<0.0001** | | | | | | |

| Genotype | Mean ± SEM | HAB Control | | AVD Control | | N |
|---|---|---|---|---|---|---|
| | | F-Ratio | p-value | F-Ratio | p-value | |
| dncG4 > Shi$^{ts}$ UN AVD shock | 62,046 ± 6,037 | - | - | 0.227 | **0.6378** | 12 |
| dncG4 > Shi$^{ts}$ **IN** AVD shock | 65,537 ± 4,130 | - | - | | | 12 |
| **Supplemental Figure 10A: 17dG4 > Shi$^{ts}$ AVOIDANCE** (w$^{1118}$ background = (+)) | | | | | | |
| ANOVA $F_{(1,19)}$=0.0036 **p<0.0001** | | | | | | |
| 17dG4 > Shi$^{ts}$ UN AVD shock | 75.914 ± 4.227 | - | - | 0.003 | **0.9526** | 10 |
| 17dG4 > Shi$^{ts}$ **IN** AVD shock | 76.281 ± 4.387 | - | - | | | 10 |
| **Supplemental Figure 10A: VT030604G4 > Shi$^{ts}$ AVOIDANCE** (w$^{1118}$ background = (+)) | | | | | | |
| ANOVA $F_{(1,22)}$=0.0888 **p<0.0001** | | | | | | |
| VT030604G4 > Shi$^{ts}$ UN AVD shock | 77.752 ± 2.717 | - | - | 0.088 | **0.7686** | 12 |
| VT030604G4 > Shi$^{ts}$ **IN** AVD shock | 76.249 ± 4.359 | - | - | | | 11 |
| **Supplemental Figure 10A: VT044966G4 > Shi$^{ts}$ AVOIDANCE** (w$^{1118}$ background = (+)) | | | | | | |
| ANOVA $F_{(1,15)}$=1.4892 **p<0.0001** | | | | | | |
| VT044966G4 > Shi$^{ts}$ UN AVD shock | 79.789 ± 3.232 | - | - | 1.489 | **0.2424** | 8 |
| VT044966G4 > Shi$^{ts}$ **IN** AVD shock | 72.777 ± 4.750 | - | - | | | 8 |
| **Supplemental Figure 10B: pleG4 > Shi$^{ts}$ AVOIDANCE** (w$^{1118}$ background = (+)) | | | | | | |
| ANOVA $F_{(2,23)}$=0.6815 **p<0.0001** | | | | | | |
| pleG4 > Shi$^{ts}$ UN AVD shock | 67.590 ± 3.420 | - | - | 0.271 | **0.6081** | 8 |
| pleG4 > Shi$^{ts}$ **IN** AVD shock | 64.900 ± 3.316 | - | - | | | 8 |
| pleG4 > (+) **IN** AVD shock | 61.570 ± 4.163 | - | - | 0.415 | **0.5261** | 8 |
| **Supplemental Figure 10B: MB504BG4 > Shi$^{ts}$ AVOIDANCE** (w$^{1118}$ background = (+)) | | | | | | |
| ANOVA $F_{(2,29)}$=0.3510 **p<0.0001** | | | | | | |
| MB504BG4 > Shi$^{ts}$ UN AVD shock | 75.623 ± 3.518 | - | - | 0.158 | **0.6936** | 10 |
| MB504BG4 > Shi$^{ts}$ **IN** AVD shock | 73.562 ± 3.544 | - | - | | | 10 |
| MB504BG4 > (+) **IN** AVD shock | 71.288 ± 3.904 | - | - | 0.193 | **0.6639** | 10 |
| **Supplemental Figure 10B: APLG4 > Shi$^{ts}$ AVOIDANCE** (w$^{1118}$ background = (+)) | | | | | | |
| ANOVA $F_{(2,48)}$=1.2055 **p<0.0001** | | | | | | |
| APLG4 > (+) **IN** AVD shock | 72.944 ± 4.717 | - | - | 1.191 | **0.2807** | 18 |
| APLG4 > Shi$^{ts}$ **IN** AVD shock | 80.396 ± 5.101 | - | - | | | 16 |
| Shi$^{ts}$ > (+) **IN** AVD shock | 69.647 ± 4.932 | - | - | 2.264 | **0.1391** | 15 |
| **Supplemental Figure 10C: dncG4 > Shi$^{ts}$ AVOIDANCE** (w$^{1118}$ background = (+)) | | | | | | |
| ANOVA $F_{(3,39)}$=0.8190 **p<0.0001** | | | | | | |
| dncG4 > Shi$^{ts}$ UN AVD yeast | 58.866 ± 4.431 | - | - | 2.036 | **0.1621** | 10 |
| dncG4 > Shi$^{ts}$ **IN** AVD yeast | 49.515 ± 5.045 | - | - | | | 10 |
| dncG4 > (+) **IN** AVD yeast | 50.708 ± 4.968 | - | - | 0.033 | **0.8564** | 10 |
| Shi$^{ts}$ > (+) **IN** AVD yeast | 51.907 ± 4.012 | - | - | 0.133 | **0.7171** | 10 |
| **Supplemental Figure 10C: 17dG4 > Shi$^{ts}$ AVOIDANCE** (w$^{1118}$ background = (+)) | | | | | | |
| ANOVA $F_{(3,39)}$=0.7012 **p<0.0001** | | | | | | |
| 17dG4 > Shi$^{ts}$ UN AVD yeast | 60.595 ± 3.830 | - | - | 2.090 | **0.1568** | 10 |
| 17dG4 > Shi$^{ts}$ **IN** AVD yeast | 68.985 ± 4.230 | - | - | | | 10 |
| 17dG4 > (+) **IN** AVD yeast | 64.275 ± 4.405 | - | - | 0.658 | **0.4222** | 10 |
| Shi$^{ts}$ > (+) **IN** AVD yeast | 64.844 ± 3.919 | - | - | 0.509 | **0.4800** | 10 |
| **Supplemental Figure 10C: VT030604G4 > Shi$^{ts}$ AVOIDANCE** (w$^{1118}$ background = (+)) | | | | | | |
| ANOVA $F_{(2,44)}$=1.2917 **p<0.0001** | | | | | | |
| VT030604G4 > Shi$^{ts}$ UN AVD yeast | 54.396 ± 3.371 | - | - | 2.581 | **0.1155** | 15 |
| VT030604G4 > Shi$^{ts}$ **IN** AVD yeast | 44.346 ± 4.888 | - | - | | | 15 |
| VT030604G4 > (+) **IN** AVD yeast | 49.157 ± 4.839 | - | - | 0.591 | **0.4461** | 15 |
| **Supplemental Figure 10C: VT044966G4 > Shi$^{ts}$ AVOIDANCE** (w$^{1118}$ background = (+)) | | | | | | |
| ANOVA $F_{(2,29)}$=0.4134 **p<0.0001** | | | | | | |

| Genotype | Mean ± SEM | HAB Control | | AVD Control | | N |
|---|---|---|---|---|---|---|
| | | F-Ratio | p-value | F-Ratio | p-value | |
| VT044966G4 > Shi$^{ts}$ UN AVD yeast | 70.273 ± 3.782 | - | - | 0.247 | **0.6226** | 10 |
| VT044966G4 > Shi$^{ts}$ **IN** AVD yeast | 67.491 ± 4.147 | - | - | | | 10 |
| VT044966G4 > (+) **IN** AVD yeast | 65.199 ± 3.917 | - | - | 0.168 | **0.6850** | 10 |
| **Supplemental Figure 10D: pleG4 > Shi$^{ts}$ AVOIDANCE** ($w^{1118}$ background = (+)) | | | | | | |
| ANOVA $F_{(1,15)}$=2.3083 **p<0.0001** | | | | | | |
| pleG4 > Shi$^{ts}$ UN AVD yeast | 80.470 ± 2.431 | - | - | 2.308 | **0.1509** | 8 |
| pleG4 > Shi$^{ts}$ **IN** AVD yeast | 75.755 ± 1.929 | - | - | | | 8 |
| **Supplemental Figure 10D: PAMG4 > Shi$^{ts}$ AVOIDANCE** ($w^{1118}$ background = (+)) | | | | | | |
| ANOVA $F_{(1,15)}$=0.0248 **p<0.0001** | | | | | | |
| PAMG4 > Shi$^{ts}$ UN AVD yeast | 80.334 ± 2.571 | - | - | 0.024 | **0.8771** | 8 |
| PAMG4 > Shi$^{ts}$ **IN** AVD yeast | 79.845 ± 1.739 | - | - | | | 8 |
| **Supplemental Figure 10D: APLG4 > Shi$^{ts}$ AVOIDANCE** ($w^{1118}$ background = (+)) | | | | | | |
| ANOVA $F_{(1,15)}$=1.6109 **p<0.0001** | | | | | | |
| APLG4 > Shi$^{ts}$ UN AVD yeast | 82.817 ± 3.141 | - | - | 1.610 | **0.2250** | 8 |
| APLG4 > Shi$^{ts}$ **IN** AVD yeast | 78.343 ± 1.597 | - | - | | | 8 |
